# Coordination between ECM and cell-cell adhesion regulates the development of islet aggregation, architecture, and functional maturation

**DOI:** 10.1101/2022.04.27.489466

**Authors:** Wilma Tixi, Maricela Maldonado, Ya-Ting Chang, Amy Chiu, Wilson Yeung, Nazia Parveen, Michael Nelson, Ryan Hart, Shihao Wang, Wu Jih Hsu, Patrick Fueger, Janel L. Kopp, Mark O. Huising, Sangeeta Dhawan, Hung-Ping Shih

## Abstract

Pancreatic islets are 3-dimensional cell aggregates consisting of unique cellular composition, cell-to-cell contacts, and interactions with blood vessels. Cell aggregation is essential for islet endocrine function; however, it remains unclear how developing islets establish aggregation. By combining genetic animal models, imaging tools, and gene expression profiling, we demonstrate that islet aggregation is regulated by extracellular matrix signaling and cell-cell adhesion. Islet endocrine cell-specific inactivation of extracellular matrix receptor Integrin β1 disrupted blood vessel interactions but promoted cell-cell adhesion and the formation of larger islets. In contrast, ablation of cell-cell adhesion molecule α-Catenin promoted blood vessel interactions yet compromised islet clustering. Simultaneous removal of Integrin β1 and α-Catenin disrupts islet aggregation and the endocrine cell maturation process, demonstrating that establishment of islet aggregates is essential for functional maturation. Our study provides new insights into understanding the fundamental self-organizing mechanism for islet aggregation, architecture, and functional maturation.

**HIGHLIGHTS:**

- Islet vascularization and aggregation are regulated via ECM-Itgb1 signaling
- ECM-Itgb1 signaling negatively controls islet aggregation via regulation of cell-cell adhesion during development
- Cell-cell adhesion negatively regulates the interaction of endocrine cell-vasculature in islets
- Differential cell adhesion regulates the establishment of islet architecture
- Endocrine functional maturation depends on islet aggregation regulated by the coordination of ECM-Itgb1 signaling and cell-cell adhesion

## INTRODUCTION

To control various physiological responses, groups of endocrine cells cluster together with vasculature, mesenchymal cells, and neuronal cells to form a highly organized “mini-organ” in the pancreatic epithelium: the islet of Langerhans. The islet endocrine cells emerge from a domain of multipotent progenitors in the embryonic pancreatic epithelium. Initially, the multipotent pancreatic progenitors resolve into a pre-acinar domain, and a bipotential endocrine/ductal progenitor domain of the developing pancreas (Larsen and Grapin-Botton, 2017; Romer and Sussel, 2015; Shih et al., 2013). Endocrine cell progenitors are specified by the expression of the pro-endocrine gene *Neurogenin 3* (*Ngn3*) (Gu et al., 2002). Following endocrine cell specification, the Ngn3^+^ precursors undergo delamination (Bechard et al., 2016; Gouzi et al., 2011; Rosenberg et al., 2010). Delaminated endocrine precursors remain associated with the ducts as they migrate and join other endocrine cells and aggregate into immature islets (Sharon et al., 2019a). These immature islets form long interconnected cords along the ductal epithelium. Later the islet cords change morphology and undergo fission to form distinct spherical islets (Jo et al., 2011; Seymour et al., 2004; Sznurkowska et al., 2020). Throughout the differentiation and morphological changes, endocrine cells are intimately associated with the vasculature, which facilitates functional control of hormone release (Reinert et al., 2014). To coordinate the hormone release of endocrine cells within each islet, autocrine and paracrine interactions, as well as direct cell contacts are required (Campbell and Newgard, 2021). These interactions are established by the aggregation process during development (Adams and Blum, 2022). Defects in the aggregation processes may lead to defects in endocrine cell development, and eventually, islet dysfunction. Supporting this notion, human pluripotent stem cell (hPSC)-derived β-cells acquire more functional maturity *in vitro* if clustered together (Cottle et al., 2021; Nair et al., 2019). However, the mechanism by which this process is regulated *in vivo* has largely remained elusive.

Self-organization of a group of cells into a higher-order, multicellular functional unit relies on an array of cell interactions regulated by cell-cell adhesions (Fagotto, 2014). For example, a calcium-dependent adhesion molecule, Neural Cell Adhesion Molecule (N-CAM), has been shown to be critical for murine endocrine cells to cluster (Esni et al., 1999). In rodents, islets form a stereotypical core-mantle architecture in which the Insulin (Ins) secreting β-cells are located mainly in the central core, with Glucagon (Gcg) secreting α-cells and Somatostatin (Sst) secreting δ-cells localizing in the periphery to form a mantle (Adams and Blum, 2022; Arrojo e Drigo et al., 2015; Kim et al., 2009). In culture, dissociated rat endocrine cells can self-organize into the core– mantle architecture, suggesting that the islet structure is established by factors intrinsic to endocrine cells (Halban et al., 1987). N-CAM deleted mice show abnormal islet architecture, with α-cells spread throughout the islet core and abnormal assembly of adherens junction proteins N- and E-cadherin (Ncad and Ecad) (Esni et al., 1999). In addition, ectopic expression of a dominant negative Ecad in mouse β-cells interferes with their clustering and islet formation (Dahl et al., 1996); and loss of *Ecad* results in abnormal blood glucose homeostasis (Wakae-Takada et al., 2013). Intracellularly, adherens junctions are stabilized by β-Catenin (*Ctnnb1*) and α-Catenin (*Ctnna1*) (Jamora and Fuchs, 2002). The function of *Ctnnb1* has been extensively studied in the developing pancreas (Elghazi et al., 2012; Murtaugh et al., 2005; Rulifson et al., 2007) and in hPSC-derived β-cells, because of its role in cell proliferation via Wnt signaling (Sharon et al., 2019b); yet genetic studies show that *Ctnnb1* is not essential for islet clustering and architecture (Elghazi et al., 2012; Murtaugh et al., 2005). In the developing pancreas, inactivation of *Ctnna1* results in defects in endocrine progenitor differentiation due to an expansion of pancreatic ductal progenitors (Jimenez-Caliani et al., 2017). Whether the process of islet clustering and architecture requires *Ctnna1* is unknown.

The regulation of cell-cell adhesion is often coupled with cell-extracellular matrix (ECM) interaction and Integrin signaling (Mui et al., 2016; Weber et al., 2011). Upon ECM binding, Integrins activate signal transduction pathways and regulate 1) cytoskeleton dynamics, 2) cell movement, 3) cell differentiation, and 4) functional maturation of epithelial cells, vasculature, and neurons (Barros et al., 2011; Li et al., 2008; Scheppke et al., 2012). During angiogenesis, ECM receptor Integrin β1 (Itgb1) is required to control the localization of cell-cell adhesion molecule VE-cadherin and maintain cell–cell junction integrity (Yamamoto et al., 2015). In salivary glands and early embryonic pancreas, Integrin β1-mediated ECM adhesions regulate Ecad to control cell clustering and regulate budding epithelial branching (Shih et al., 2016; Wang et al., 2021; Wang et al., 2017). Whether islet morphogenesis also requires ECM-Integrin signaling remains to be investigated.

In the present study, we examine how loss of *Itgb1* and *Ctnna1* in Ngn3-expressing endocrine progenitors affects islet morphogenesis. We show that *Itgb1* and *Ctnna1* have collaborative roles during islet aggregation: ECM-Itgb1 signaling modulates islet aggregation by negatively regulating cell-cell adhesions; and α-Catenin promotes endocrine cell clustering and modulates vascularization during islet development. We also provide evidence that the islet architecture is regulated by differential adhesion in endocrine subtypes.

## RESULTS

### The morphogenesis and functional maturation of islet endocrine cells depend on ECM-Itgb1 signaling during development

To investigate the process of islet aggregation, embryonic day 15.5 (E15.5) pancreas was stained for blood vessel maker PECAM, ductal marker Spp1, and endocrine cell marker Chromogranin A. Confocal microscopy and 3D whole-mount immunofluorescence imaging revealed that at E15.5 endocrine cells had started aggregating, and the vascular and ductal networks were already highly similar (Figure S1A). This close relation between the ducts and blood vessels at the stage prior to islet aggregation suggests that blood vessels could readily interact and provide instructive cues to regulate endocrine progenitor aggregation into islets. Blood vessels in islets produce ECM-rich basal membrane to support and organize endocrine cells (Lammert et al., 2003; Roscioni et al., 2016). Thus, we hypothesized that ECM signaling, provided by the vasculature, plays a key role in the initiation steps of islet aggregation. Supporting this notion, published work using single-cell gene expression profiling shown that the ECM-Itgb1 signaling gene ontology (GO) categories are most significantly enriched in the Ngn3^+^ endocrine progenitors (Bastidas-Ponce et al., 2019). Immunofluorescence (IF) analysis validated that both the Ngn3^+^ endocrine progenitors and their progenies (eYFP^+^ cells in *Ngn3-Cre; Rosa26-eYFP* pancreas) expressed Itgb1 (Figure 1A). To investigate the role of Itgb1 during endocrine cell aggregation, we used the *Ngn3-Cre* allele to recombine the *Itgb1-flox* alleles in endocrine progenitors (*Ngn3-Cre*; *Itgb1^f/f^*; hereafter *Itgb1*^Δ*Endo/*Δ*Endo*^). Expression of Itgb1 was almost entirely absent throughout the islet endocrine cells in comparison to control littermates at 6 weeks old (Figure 1B,1C). We first examined whether inactivation of ECM-Itgb1 signaling affected vascular interaction. Electron microscope ultrastructure analysis revealed tight attachment in basal membranes between vascular and endocrine cells in control islets (Figure S1B). In contrast, in the *Itgb1*^Δ*Endo/*Δ*Endo*^ islets, the endocrine cells were detached from the basal membranes of blood vessels (Figure S1C, red arrows). Despite the vascular defects with Itgb1 loss, all four endocrine cell subtypes, α-, β-, δ-, and Ppy-(pancreatic polypeptide) cells, were present at similar proportions in control and *Itgb1*^Δ*Endo/*Δ*Endo*^ islets (Figure 1D,1E;S1D-S1G). In addition, the expression of β-cell lineage markers Nkx6.1 and Pdx1 were present in the Ins^+^ cells of control and *Itgb1*^Δ*Endo/*Δ*Endo*^ islets (Figure 1F,1G). Although the populations of endocrine cell lineages were unaffected by the loss of *Itgb1*, morphometrical analysis revealed drastic morphological changes. In control islets, β-cells were organized in a central core and surrounded by α- and δ- cells in the periphery (Figure 1D). Conversely, in *Itgb1*^Δ*Endo/*Δ*Endo*^ mice, several individual islets appeared to agglomerate, resulting in larger islet sizes and aberrant distribution of α- and δ- cells in the central core (Figure 1E,1G; S1I). Thus, the inactivation of Itgb1 has a profound impact on islet morphogenesis, but not endocrine cell fate choice.

**Figure 1.**
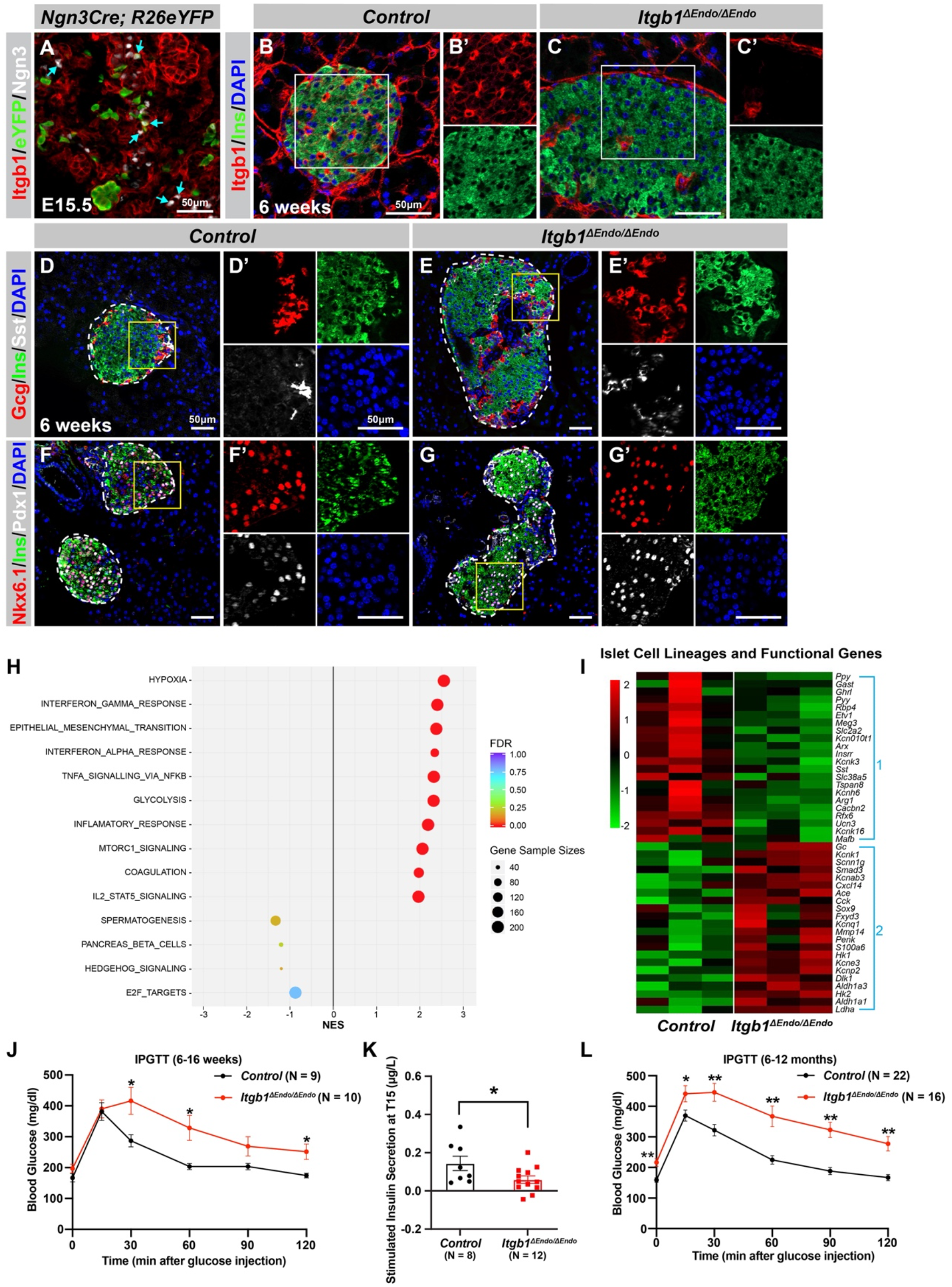
ECM-Itgb1 signaling in endocrine progenitors is required for normal islet morphology and function. A) Immunofluorescence staining for Itgb1 and Ngn3 in pancreas sections of *Ngn3-Cre; Rosa26eYFP* reporter mice at E15.5. Cyan arrows indicate Ngn3^+^ cells and lineage-traced endocrine cells expressing Itgb1. (B-C) Immunofluorescence staining for Itgb1, Ins, and DAPI on pancreas sections, demonstrating significant *Itgb1* deletion in an *Itgb1^ΔEndo/ΔEndo^* islet at 6 weeks old. Fields demarcated by white boxes in B-C are shown with individual color channels in B’-C’ side panels. (D-G) Immunofluorescence staining for Gcg, Ins, Sst, Nkx6.1, Pdx1, and DAPI on pancreas sections of (D,F) control and (E,G) *Itgb1^ΔEndo/ΔEndo^* mice at 6 weeks old. Individual islets are outlined by dotted lines. Fields demarcated by yellow boxes in D-G are shown at higher magnification in D’-G’ side panels. (H) Gene-set enrichment analysis of the differentially expressed genes, indicating top enriched pathways for control versus *Itgb1^ΔEndo/ΔEndo^* islets at 6 weeks old. Gene sample sizes and FDRs are indicated by the size and color of the dots. (I) Heatmap demonstrating down-regulation (Group 1) and up-regulation (Group 2) of functional maturation genes in *Itgb1^ΔEndo/ΔEndo^* islets compared to controls. (J-L) Intraperitoneal glucose tolerance test (J,L) and stimulated insulin secretion at T15 (K) on 6 to 16 weeks old (J,K) and (L) 6 to 12 months old control and *Itgb1^ΔEndo/ΔEndo^* mice. Ngn3, Neurogenin 3; Gcg, Glucagon; Ins, Insulin; DAPI, 4’,6-diamidino-2-phenylindole; Sst, Somatostatin; IPGTT, intraperitoneal glucose tolerance test; FDR, false discovery rate; E15.5, embryonic day 15.5 Data are shown as mean ± SEM. * p < 0.05 and ** p < 0.01.

We next investigated whether the loss of *Itgb1* had any impact on the expression of genes related to islet function. Islets from 6 weeks old control and *Itgb1*^Δ*Endo/*Δ*Endo*^ mice were isolated and subjected to RNA-seq and differential gene expression analysis. A comparison of gene-expression profiles between *Itgb1*^Δ*Endo/*Δ*Endo*^ and control islets revealed 417 differentially expressed genes (DEGs) with false discovery rate (FDR) < 0.05 and fold change (FC)>2 (Table S1), of which 332 were up-regulated and 85 were down-regulated. Gene-set enrichment analysis (GSEA) of DEGs between control and *Itgb1*^Δ*Endo/*Δ*Endo*^ islets showed up-regulation of pathways associated with hypoxia, epithelial mesenchymal transition, glycolysis, and inflammatory response (Figure 1H). The analysis of Gene Ontology (GO) of Differentially Expressed Genes (DEGs) related to hypoxia showed that the genes responsible for hypoxia response were up-regulated in *Itgb1*^Δ*Endo/*Δ*Endo*^ islets when compared to the control group (as shown in Figure S1L). The immunofluorescence (IF) analysis indicated that the *Itgb1*^Δ*Endo/*Δ*Endo*^ islets at 2-3 months old displayed a significant reduction in blood vessels (as shown in Figure S1M-S1O), which suggests that the lack of vascular interaction could lead to hypoxia responses. On the other hand, the down-regulated genes were associated with spermatogenesis, pancreas β-cells, and E2F targets (Figure 1H). GO analysis of DEGs in the pancreas β-cells category further revealed that functional maturation genes for β-cells (e.g., *Slc2a2* (*Glut2*), *Ucn3*, and *Mafa*) and for α-cells (e.g., *Arx*, *Mafb*, and *Etv1*), were drastically down-regulated in the *Itgb1*^Δ*Endo/*Δ*Endo*^ islets (Figure 1I, Group 1, S2A). IF analysis also validated the reduced expression of Mafa, Ucn3, and Glut2 in β-cells of *Itgb1*^Δ*Endo/*Δ*endo*^ islets (Figure S2B-E). Furthermore, several “disallowed” genes for β-cells, including *Ldha1*, *Aldh1a3*, and *Hk1* (Lemaire et al., 2016), were significantly up-regulated in the *Itgb1*^Δ*Endo/*Δ*Endo*^ islets (Figure 1I, Group 2, S2A). Hence, it is likely that the functional maturation of endocrine cells within *Itgb1*^Δ*Endo/*Δ*Endo*^ islets is compromised. This notion is supported by the observation that, by 6 weeks of age, these mice exhibit glucose intolerance and reduced insulin secretion in response to glucose stimulation (Figure 1J-K). The impairment in glycemic regulation further deteriorates in mice older than 6 months (Figure 1L). Together, these observations indicate that ECM-integrin interactions are critical for islet vascularization, morphogenesis, and the functional maturation of endocrine cells.

### ECM signaling regulates endocrine progenitor cell migration and cell-cell adhesion to control islet aggregation during development

Our initial analysis of the pancreatic sections from *Itgb1*^Δ*Endo/*Δ*Endo*^ mice revealed larger islet sizes (Figure S1J). Consistently, the islets isolated from *Itgb1*^Δ*Endo/*Δ*Endo*^ mice appeared larger in size as well (Figure 2A-2C); however, the total islet area was similar between control and *Itgb1*^Δ*Endo/*Δ*Endo*^ mice (Figure 2D). The increased size was mainly attributed to a greater number of large islets (>50 µm^2^ x 10^3^, as shown in Figure S1K,2E). Therefore, the enlarged islet size is likely a result of a morphological change independent of the expansion of total endocrine cells. Furthermore, in the control pancreata, the islets were located near the ducts, but contained a very small number of ductal cells (Figure 2F,2H,S3A). In contrast, the islets in the *Itgb1*^Δ*Endo/*Δ*Endo*^ pancreata contained a substantial number of ductal cells (Figure 2G,2H,S3B). As the endocrine progenitors arise and migrate away from the ducts, we hypothesized that there might be deficiencies in this process for Itgb1-deficient endocrine cells. To investigate this, we conducted segmentation analysis of 3D whole-mount immunofluorescence on E15.5 pancreata, the stage at which the majority of endocrine progenitor migration occurs (Rosenberg et al., 2010). Our results showed that most of the Itgb1-deleted endocrine progenitor cell aggregates had a greater tendency to cluster within the ductal cords during development (Figure 2I,2J and Movie S1; the green color represents zero distance from ducts). These findings imply that there might be deficiencies in both endocrine cell migration and premature islet aggregation in *Itgb1*-deleted endocrine cells. To further confirm this, we conducted live-imaging analysis on pancreas explants obtained from E16.5 *Ngn3-Cre; R26mT/mG; Itgb1^+/+^* (control) and *Ngn3-Cre; R26mT/mG; Itgb1^f/f^*(*Itgb1^ΔEndo/ΔEndo^*) mice. We tracked the movement and aggregation of Ngn3-progeny (mGFP^+^) endocrine progenitor cells in explant cultures for up to 3 days (Figure S3C,S3D; Movie S2,S3). Our observations showed that in contrast to the controls, where endocrine progenitor cells migrated and aggregated outside of ductal cords, the mGFP^+^ cells in *Itgb1^ΔEndo/ΔEndo^* explants readily aggregated before moving away from the cords (Figure S3D white arrows). In contrast to our findings, previous studies using *Ins-Cre* or *RIP-Cre* to ablate *Itgb1* after endocrine progenitors have differentiated into insulin-producing cells did not report any observable islet aggregation or migration defects (Diaferia et al., 2013; Win et al., 2020). However, *Ngn3-Cre* mediated *Itgb1* deletion would occur much earlier in embryonic stages (Figure 1A), suggesting that Itgb1 signaling regulates the migration and aggregation of progenitor cells in the endocrine lineage during the islet developmental process.

**Figure 2.**
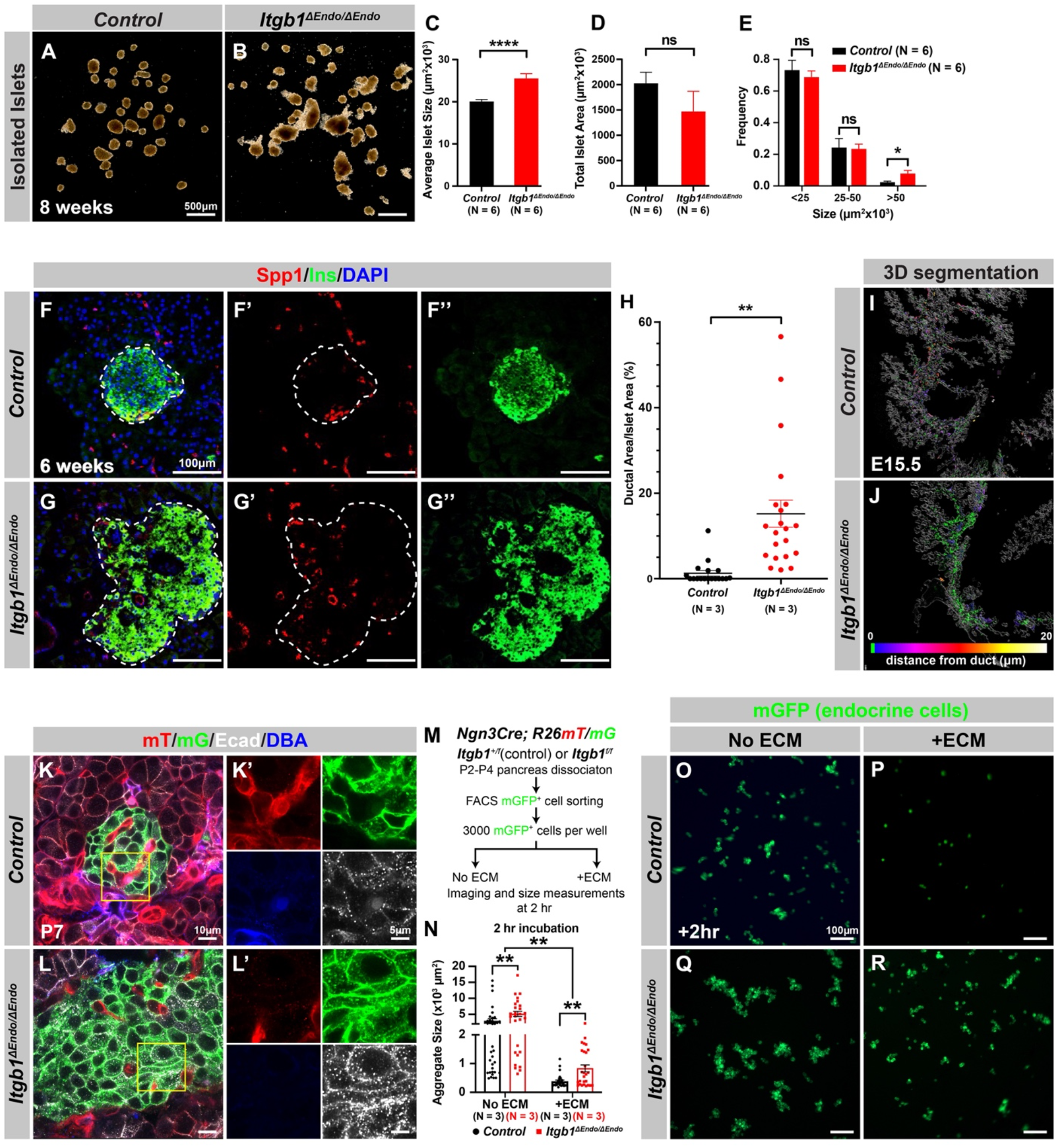
Itgb1 signaling is required for proper endocrine progenitor aggregation. (A-B) Brightfield images of isolated islets from control and *Itgb1^ΔEndo/ΔEndo^*mice at 8 weeks old. (C-E) Quantification of (C) average islet size, (D) total islet area, and (E) islet size distribution based on morphometric analysis of 8 weeks old control and *Itgb1^ΔEndo/ΔEndo^*isolated islets. (F-G) Immunofluorescence staining for Spp1, Ins, and DAPI on 6 weeks old control and *Itgb1^ΔEndo/ΔEndo^* ancreas sections. Islets are outlined by white dotted lines. (H) The ratio of ductal cell areas within the islets is quantified based on Spp1 and ChrA staining of pancreas sections from 6-week-old control and *Itgb1^ΔEndo/ΔEndo^*mice. (I-J) 3D segmentation analysis on whole mount immunofluorescence images of E15.5 control and *Itgb1^ΔEndo/ΔEndo^* pancreata stained for ChrA and Spp1 (white colored) reveals that the endocrine cells in the *Itgb1^ΔEndo/ΔEndo^* pancreas are mostly retained within ducts, and the distance between endocrine cells and ducts is represented by a color scale. (K-L) Airyscan super-resolution images depict immunofluorescence staining for Ecad and ductal marker DBA in P7 mT/mG reporter mice showing up-regulation of Ecad in the *Itgb1^ΔEndo/ΔEndo^* pancreas. The dot-like Ecad expression pattern in super-resolution imaging represents adherens junction structures. The yellow boxes in K-L demarcate fields shown at higher magnification in K’-L’ side panels. (M) The schematic depicts the experimental design for the *in vitro* aggregation assay. (N) Aggregate size is quantified after a 2-hour incubation with or without ECM. (O-R) Fluorescence images of mGFP+ endocrine progenitor aggregates after 2 hours of culture show that mGFP+ control endocrine progenitors clustered (O) without ECM but remained single cells (P) with ECM. Alternatively, *Itgb1^ΔEndo/ΔEndo^* endocrine progenitors clustered (Q) without ECM or (R) with ECM. Spp1, Secreted Phosphoprotein 1; Ins, Insulin; DAPI, 4’,6-diamidino-2-phenylindole; Ecad, E-cadherin; DBA, Dolichos Biflorus Agglutinin; P7, postnatal day 7; E15.5, embryonic day 15.5 Data are shown as mean ± SEM. * p < 0.05, ** p < 0.01 and **** p < 0.0001.

To further investigate this notion, we sought to inactivate Itgb1 only after the endocrine cells had fully differentiated and matured. Urocortin 3 (Ucn3) is a late β-cell-specific maturity marker in mouse islets that is expressed after *Ins1* and *Ins2* and starts perinatally. Most β-cells express Ucn3 by 3 weeks of age (Blum et al., 2014; van der Meulen and Huising, 2014). The expression pattern of Cre recombinase under the control of the *Ucn3* promoter aligns with this timeline (van der Meulen et al., 2017). Therefore, *Ucn3-Cre* mediated deletion of *Itgb1* is not expected to occur until islet development has completed in the presence of functional Itgb1 alleles and normal Itgb1 expression. To test whether Itgb1 signaling also maintains islet cell aggregation after β-cell maturation and islet formation, we crossed *Ucn3-Cre* with *Itgb1^f/f^* mice to inactivate Itgb1 exclusively in mature β-cells (referred to as *Itgb1^ΔMatureβ/ΔMatureβ^*). Our analysis revealed that, similar to *Itgb1^ΔEndo/ΔEndo^* mice, *Itgb1^ΔMatureβ/ΔMatureβ^* mice exhibited vasculature exclusion from islets (Figure S3E’,S3F’), but their islet size remained similar to control mice (Figure S3E-S3I). These results suggest that Itgb1 signaling specifically regulates islet cell aggregation during pancreatic development.

During the early stages of pancreas development, the expression of Itgb1 in pancreatic progenitor cells has been identified as a crucial factor in controlling pancreas branching morphogenesis through its regulation of cell-cell adhesion (Shih et al., 2016). As the process of endocrine progenitor cell differentiation and islet formation commences, cell-cell adhesion is typically downregulated to facilitate the migration and interaction of endocrine cells with non-endocrine cells (Gouzi et al., 2011). Given the established role of Itgb1 in regulating cell-cell adhesion in the context of pancreas development, we were prompted to investigate whether Itgb1 signaling also plays a role in instructing endocrine progenitor cells to modulate their adhesion behavior and thereby control the islet aggregation process. To address this question, we took Airyscan super-resolution images of Ecad IF on endocrine cells in *Ngn3-Cre; R26mT/mG; Itgb1^f/f^* islets at P7, since islet aggregation is complete by the neonatal stages (Sznurkowska et al., 2020). The Airyscan images provide sufficient resolution to show adherens junctions between cells by visualizing Ecad^+^ clusters ((Gonschior et al., 2020); white dots in Figure 2K,2L). The lineage-traced endocrine cells (mGFP^+^) in control islets displayed low amounts of Ecad^+^ clusters (Figure 2K’). However, the amount of Ecad^+^ clusters was drastically increased in mutant islets (Figure 2L’), suggesting that Itgb1 signaling negatively regulates cell adhesion in developing islets. To further test whether the increased cell adhesiveness is responsible for the increased islet size when Itgb1 signaling is disrupted, we performed an *in vitro* cell aggregation assay. Live mGFP^+^ endocrine cells, from neonatal (P2-P4) *Ngn3-Cre; R26mT/mG; Itgb1^+/f^* or *Ngn3-Cre; R26mT/mG; Itgb1^f/f^* pancreata, were completely dissociated, then collected by FACS. 3000 mGFP^+^ cells were allowed to aggregate in wells of a 96-well plate that were coated with bovine serum albumin (no ECM) or Matrigel and fibronectin (+ECM) (Figure 2M). Within 2 hours of cell seeding, the control mGFP^+^ endocrine cells rapidly aggregated in wells with no ECM coating, but formed no aggregates in the ECM-coated wells (Figure 2N-2P). Within the same amount of time, the mutant mGFP^+^ endocrine cells clustered into larger aggregates in wells without ECM coating and maintained aggregation in the ECM-coated wells (Figure 2N,2Q,2R). After 48 hours, the mGFP^+^ cells isolated from both the controls and mutants progressively clustered into larger aggregates in wells without ECM coating (Figure S4A-S4C,S4E). In contrast, in the ECM-coated wells, only the mutant mGFP^+^ cells clustered, while control mGFP^+^ cells remained unaggregated (Figure S4D,S4F). Together, these findings demonstrated that ECM-Itgb1 signaling negatively regulates islet aggregation. As Integrins play a crucial role in linking the actin cytoskeleton, we examined whether the inactivation of Itgb1 in endocrine progenitors affects the organization of the actin cytoskeleton in islet cells. In *Itgb1*^Δ*Endo/*Δ*Endo*^ islets, we observed pronounced foci of condensed F-actin between the endocrine cells (Figure S4G-S4J), indicating increased cell-cell adhesion. These findings suggest that the loss of Itgb1 in endocrine cells leads to alterations in the actin cytoskeleton, resulting in enhanced cell-cell adhesion within the islet microenvironment. The concentrated F-actin foci between the endocrine cells indicate a potential mechanism through which Itgb1 regulates cell-cell interactions and influences islet development and aggregation.

### α-Catenin promotes endocrine cell aggregation and regulates vascularization during islet development

Our study revealed that Itgb1 plays a critical role in islet development by regulating endocrine cell-cell adhesion. However, the mechanism by which cell-cell adhesion regulates islet formation remains unclear. Although Ecad and Ncad-mediated adherens junctions are believed to be the primary cell-cell adhesion mechanisms linking islet cells into aggregates, our findings and other studies have shown that deletions of Ecad or Ncad alone do not affect islet aggregation, suggesting functional redundancy between these cell adhesion molecules.

α-Catenin (Ctnna1) is an essential hub protein linking Ecad-mediated adherens junctions to the cytoskeleton and has been demonstrated to be indispensable for the formation of adherens junctions(Jamora and Fuchs, 2002). Therefore, to investigate the requirement of cell adhesion in islet formation, we conditionally ablated Ctnna1 in endocrine progenitors by generating *Ngn3-Cre*; *Ctnna1^f/f^* mice (hereafter referred to as *Ctnna1*^Δ*Endo/*Δ*Endo*^). To determine whether *Ctnna1* deletion affects islet development, Chromogranin A (ChrA) staining on pancreas sections from 8 weeks old mice were imaged, and the total relative islet area was measured (Figure 3A,3B). Compared to their wild-type littermates, islet area (relative to the total pancreas area) in *Ctnna1*^Δ*Endo/*Δ*Endo*^ mice was unchanged (Figure 3C), but there was a significant reduction in average islet size (Figure 3D). The proportion of larger islets was significantly reduced, while the proportion of small islets was increased (Figure S5A-S5C), suggesting that *Ctnna1* is required for proper islet aggregation, but not for the development of total islet cell mass. In *Ctnna1*^Δ*Endo/*Δ*Endo*^ islets, we observe diminished Ecad expression and a decrease in the density of F-actin at cell junctions. These findings imply a disruption of cell junctions and a reduction in cell adhesion among Ctnna1-deficient endocrine cells (Figure 3E, 3F).

**Figure 3.**
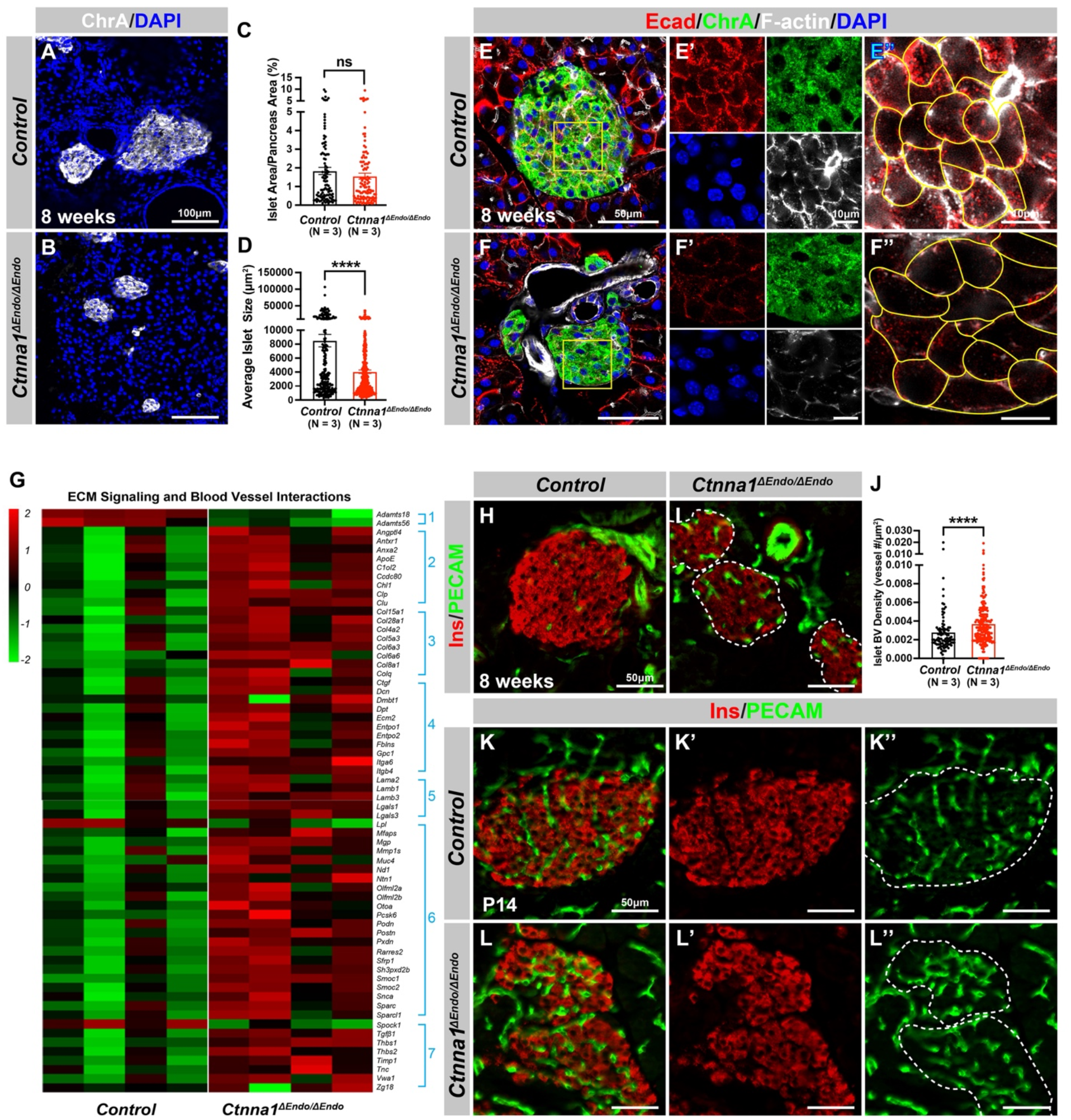
Ctnna1-mediated cell-cell adhesion is required for normal islet morphology and endocrine cell aggregation. (A-B) Immunofluorescence staining for ChrA and DAPI in 8 weeks old pancreas sections showing smaller islet sizes in *Ctnna1^ΔEndo/ΔEndo^* mice. (C-D) Quantification of (C) the percentage of islet area in pancreas area and (D) average islet size based on ChrA staining on 8 weeks old control and *Ctnna1^ΔEndo/ΔEndo^*pancreas sections. (E-F) Airyscan images of immunofluorescence staining for Ecad, ChrA, F-actin, and DAPI for control and *Ctnna1^ΔEndo/ΔEndo^* pancreas sections. Fields demarcated by yellow boxes are shown at higher magnification with individual color channels in E’-F’ middle panels, showing reduced expression of Ecad and F-actin in *Ctnna1^ΔEndo/ΔEndo^*islets. Individual endocrine cell shape is delineated by yellow lines in E”-F”, demonstrating the enlarged endocrine cell sizes in the islets of *Ctnna1^ΔEndo/ΔEndo^* mice. (G) Heatmap showing genes for ECM signaling and blood vessel interactions are up-regulated in *Ctnna1^ΔEndo/ΔEndo^* islets at 8 weeks old. Groups 1-7 represent gene ontologies of: 1) Metalloproteases, 2) Vascular Adhesion 3) Collagens, 4) Integrins, 5) Laminins, 6) Glycoproteins and 7) Thrombospondins. (H-I) Immunofluorescence staining of Ins and PECAM on 8 weeks old control and *Ctnna1^ΔEndo/ΔEndo^* pancreas sections. (J) Quantification of islet blood vessel density based on Ins and PECAM staining on 8 weeks old control and *Ctnna1^ΔEndo/ΔEndo^* pancreas sections. (K-L) Immunofluorescence staining of Ins and PECAM on P14 control and *Ctnna1^ΔEndo/ΔEndo^*pancreas sections showing increased vasculature in *Ctnna1^ΔEndo/ΔEndo^* islets. Islet area is outlined by dotted lines. ChrA, Chromogranin A; DAPI, 4’,6-diamidino-2-phenylindole; Ecad, E-cadherin; Ins, Insulin; PECAM, Platelet endothelial cell adhesion molecule-1; P14, postnatal day 14 Data are shown as mean ± SEM. **** p < 0.0001.

To investigate how *Ctnna1*-deletion impacts islet development, we compared transcriptional profiling of *Ctnna1*^Δ*Endo/*Δ*Endo*^ and control islets at 8 weeks. A comparison of gene expression profiles between *Ctnna1*^Δ*Endo/*Δ*Endo*^ and control islets revealed significant differences in the expression of 319 genes with FDR < 0.05 and FC>2, of which 245 were up-regulated and 74 were down-regulated (Table S2). Among all GO categories, one of the most up-regulated groups in *Ctnna1*^Δ*Endo/*Δ*Endo*^ islets is genes associated with ECM signaling and blood vessel interactions (Figure 3G, S5D). These include Metalloproteases, Vascular Adhesion, Collagens, Integrins, Laminins, Glycoproteins and Thrombospondins (Figure 3G). These findings suggested that *Ctnna1*^Δ*Endo/*Δ*Endo*^ islets have augmented ECM signaling and blood vessel interaction. IF for blood vessels confirmed an increase in blood vessel density in *Ctnna1*^Δ*Endo/*Δ*Endo*^ islets at 8 weeks (Figure 3H-3J). The increase in islet vasculature was observed as early as P14 (Figure 3K-3L). These findings demonstrated that the inactivation of *Ctnna1* in endocrine progenitor cells led to a decrease of endocrine cell-cell adhesion and an increase in blood vessel interaction.

Alongside the increased interaction with blood vessels, we observed abnormal organization of F-actin fibers within *Ctnna1^ΔEndo/ΔEndo^* islets (Figure 3E, 3F; S5E, S5F). These alterations in F-actin fiber organization were evident as early as P14 in the *Ctnna1^ΔEndo/ΔEndo^* islets, concurrent with the onset of enhanced interaction with blood vessels (Figure S5G, S5H). These findings demonstrate that the loss of Ctnna1 compromises endocrine cell aggregation and perturbs the organization of the actin cytoskeleton and cell-cell adhesion. Together, these findings suggest that the balance between cell-cell adhesion and blood vessel interaction is crucial for islet cell aggregation during development (Figure S5I).

### Inactivation of Ctnna1 in differentiating endocrine cells affects the assembly and maintenance of non-β endocrine cells in islets

To further investigate the impact of Ctnna1-dependent islet cell aggregation on the islet development process, we compared the gene expression profiles of *Ctnna1*^Δ*Endo/*Δ*Endo*^ and control islets. Our analysis revealed significant misregulation of genes crucial for endocrine cell functions. Specifically, we observed a down-regulation of *Slc27a2*, *Tmsb15b1*, and *Etv1* expression, which are genes required for glycemic control and insulin secretion (Figure 4A Group 1, S6A,). Conversely, we found an up-regulation of *Hk1*, *Ldha*, *Aldh1a1*, and *Cxcl14* expression, which are genes known to be “disallowed” in mature functional β-cells (Lemaire et al., 2016) (Figure 4A Group 2, S6A). These findings suggest that the function of β-cells may be negatively impacted in Ctnna1^ΔEndo/ΔEndo^ mice.

**Figure 4.**
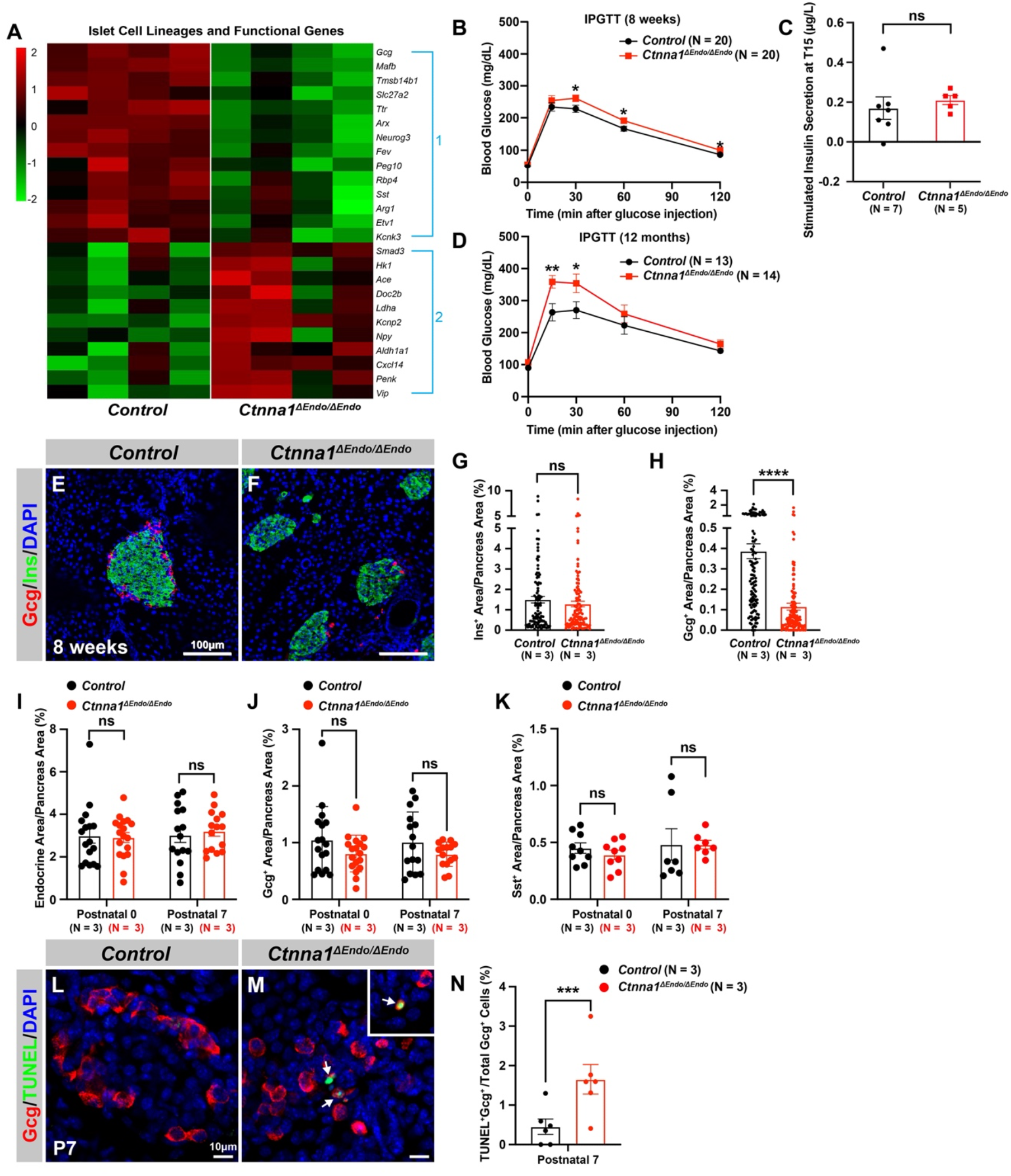
Ctnna1 is required for proper islet functionality and α-cell survival. (A) Heatmap showing the top up-regulated (Group 1) and down-regulated (Group 2) endocrine cell lineage and functional genes in *Ctnna1^ΔEndo/ΔEndo^* islets from 8 weeks old mice. (B-D) The figure shows the results of the intraperitoneal glucose tolerance test (B,D) and stimulated insulin secretion assay (C) performed on control and *Ctnna1^ΔEndo/ΔEndo^* mice at 8 weeks old (B,C) and 12 months old (D). The *Ctnna1^ΔEndo/ΔEndo^* mice exhibited glucose intolerance, but no detectable difference in insulin secretion compared to control littermates. The *Ctnna1^ΔEndo/ΔEndo^*mice are glucose intolerant with non-detectable insulin secretion defects. (E-F) Immunofluorescence staining for Gcg, Ins, and DAPI on 8 weeks old (E) control and (F) *Ctnna1^ΔEndo/ΔEndo^* pancreas sections showing a reduction of α-cells. (G-H) Quantification of (G) Ins^+^ and (H) Gcg^+^ area relative to total pancreas area in 8 weeks old control and *Ctnna1^ΔEndo/ΔEndo^* pancreas sections. (I-K) Quantification of endocrine cell area (I), Gcg^+^ (J), and Sst^+^ (K) relative to total pancreas area in control and *Ctnna1^ΔEndo/ΔEndo^* pancreas sections at P0 and P7. Note that there was no significant difference observed between the control and *Ctnna1^ΔEndo/ΔEndo^* pancreas sections at P0 and P7. (L-N) Immunofluorescence staining for Gcg, TUNEL, and DAPI on P7 control and *Ctnna1^ΔEndo/ΔEndo^*pancreas sections. White arrows indicate TUNEL^+^/Gcg^+^ cells. The percentage of TUNEL^+^/Gcg^+^ co-positive cells relative to total Gcg^+^ cell numbers in P7 control and *Ctnna1^ΔEndo/ΔEndo^* islets is shown in panel (N), and compared to controls, the *Ctnna1^ΔEndo/ΔEndo^* islets exhibited significantly more TUNEL^+^/Gcg^+^ cells at P7. Gcg, Glucagon; Ins, Insulin; DAPI, 4’,6-diamidino-2-phenylindole; IPGTT, intraperitoneal glucose tolerance test; TUNEL, terminal deoxynucleotidyl transferase dUTP nick end labeling; P0, postnatal day 0; P7, postnatal day 7 Data are shown as mean ± SEM. * p < 0.05, ** p < 0.01, *** p < 0.001, **** p < 0.0001.

To support this notion, we conducted glucose tolerance tests, which revealed mild glucose intolerance in *Ctnna1*^Δ*Endo/*Δ*Endo*^ mice at 8 weeks (Figure 4B). However, the quantification of insulin content in serum after glucose challenge did not reveal any noticeable insulin secretion defect in the *Ctnna1*^Δ*Endo/*Δ*Endo*^ mice (Figure 4C). We reasoned that the conventional serum insulin measurement may not reflect the changes in β-cell function, given that the *Ctnna1*^Δ*Endo/*Δ*Endo*^ mice exhibit only mild glucose intolerance. Therefore, we performed quantification of insulin secretion *in vivo* using hyperglycemic clamps, which is a more sensitive detection method. In this assay, we observed a trend towards lower overall insulin secretion in the *Ctnna1*^Δ*Endo/*Δ*Endo*^ mice at 8 weeks (Figure S6B, S6C), providing additional evidence that Ctnna1-dependent islet cell aggregation has an impact on β-cell function. In addition, the glucose intolerance phenotype became even more pronounced at one year of age, as evidenced by glucose tolerance test (Figure 4D). These findings indicate that Ctnna1-dependent islet cell aggregation is crucial for the proper development of β-cells and their function in maintaining glucose homeostasis.

In addition to suggested functional defects in β-cells, we found that genes for non-β endocrine cell lineages, including α-cell genes *Gcg, MafB, Arx, Peg10,* and *Etv1,* and the δ-cell gene *Sst*, were significantly down-regulated in *Ctnna1*^Δ*Endo/*Δ*Endo*^ islets (Figure 4A; S6A,), suggesting the requirement of *Ctnna1* for the presence of non-β endocrine cells in islets. Interestingly, *Ctnna1*^Δ*Endo/*Δ*Endo*^ islets did not show a reduced area of β-cells, as β-cells were still organized loosely in the core of islets (Figure 4E-4G). However, there was a severe reduction of α-cell and δ-cell populations in the *Ctnna1*-deleted islets (Figure 4H; S6D-S6F). To determine whether the loss of non-β endocrine cells was the result of differentiation defects during development, we examined the population of α- and β-cells in *Ctnna1*^Δ*Endo/*Δ*Endo*^ islets at P0 and P7. At these ages, the islets of *Ctnna1*^Δ*Endo/*Δ*Endo*^ mice exhibited normal endocrine cell mass, Gcg^+^ α-cell and Sst^+^ δ- cell areas (Figure 4I-4K; S6G-S6J), demonstrating that the loss of α- and δ-cell was not due to differentiation defects prior to these early stages. However, the detachment of α-cells from the mantle of *Ctnna1*^Δ*Endo/*Δ*Endo*^ islets was observed by P7. By this time point, α-cells had formed tight interactions with other endocrine cells in control islets (Figure S6I,S6K), but *Ctnna1*-deficient α-cells detached from other cell types (Figure S6J,S6L). To investigate whether cell detachment promotes cell death, thus resulting in α-cell loss, TUNEL analysis on P7 *Ctnna1*^Δ*Endo/*Δ*Endo*^ islets was conducted to analyze cell apoptosis. Compared to the control, *Ctnna1*^Δ*Endo/*Δ*Endo*^ islets display 3 times more TUNEL^+^ apoptotic α-cells (Figure 4L-4N), indicating that *Ctnna1* is required for α- cell survival. Thus, the loss of the α-cells in *Ctnna1*^Δ*Endo/*Δ*Endo*^ islets during adulthood is not due to endocrine cell differentiation defects; instead, a drastic increase in apoptosis explains the loss of α-cells in the *Ctnna1*-deleted islets.

### Differential adhesion in endocrine cells controls the formation of islet architecture

Since the reduction of cell-cell adhesion caused distinctive outcomes in different endocrine cell subtypes, we hypothesized that the endocrine cells in the mantle or core of islets may have different adhesiveness. We re-analyzed publicly available gene expression profiles (DiGruccio et al., 2016) to examine the expression of key cell adhesion molecules *E-cadherin*, *N-cadherin*, *α- Catenin*, and *β-Catenin* between the subtypes of endocrine cells. These adhesion genes were expressed at the highest level in β-cells, while their expression was comparatively lower in endocrine cells located in the mantle of islets (α-cells and δ-cells, Figure S7A). These observations suggest that the adhesive properties may differ between endocrine cell subtypes. According to the differential adhesion hypothesis (DAH), a population of cells with different adhesive properties will tend to spontaneously sort themselves to maximize adhesive bonding (Foty and Steinberg, 2005). Stronger adhesion between β-cells may lead to them bundling into islet cores, while the weaker adhesion between non-β-cells results in their dispersion into islet mantles.

Based on the DAH, if the cell adhesion is altered in different endocrine subpopulations, the stereotypical islet architecture will also be affected. To alter the cell adhesion in different endocrine subpopulations, we crossed *Ins1-Cre* and *Gcg-iCre* lines to *Ctnna1-flox* mice and traced the *Ctnna1-*deleted β- or α-cells with the lineage tracer *Rosa26-mT/mG* (*Ins1-Cre*; *Ctnna1^f/f^*; *Rosa26-mT/mG* and *Gcg-iCre; Ctnna1^f/f^*; *Rosa26-mT/mG,* hereafter *Ctnna1^Δβ/Δβ^*and *Ctnna1^Δα/Δα^* mice, respectively). The *Rosa26-mT/mG* lineage-tracing allele faithfully reported the *Ins1-Cre* and *Gcg-iCre* cell lineages at P7 (Figure 5A-5H), allowing examination of the respectively labeled β- and α-cells using mGFP expression. The mGFP^+^ β-cells of *Ctnna1^Δβ/+^* mice are mostly located on the islet core (Figure 5A,5E), and the mGFP^+^ α-cells of *Ctnna1^Δα/+^* mice are mostly located in the islet mantle (Figure 5C,5G). Additionally, we found that β-cells exhibited strong F-actin condensed cellular junctions with well-organized rosette structures (S7B, yellow arrows). In contrast, α-cells displayed loosely bundled F-actin assemblies without focal-condensed junctional structures found in control mice (Figure S7D, yellow arrows), supporting the notion that α-cells may have relatively low adhesive properties. Inactivation of Ctnna1 in β-cells and α-cells significantly diminished the formation of F-actin, as indicated in Figure S7C and S7E (highlighted by yellow arrows). This suggests a substantial decrease in cell adhesion assemblies within the β-cells of *Ctnna1^Δβ/Δβ^* and the α-cells of *Ctnna1^Δα/α^* islets.

**Figure 5.**
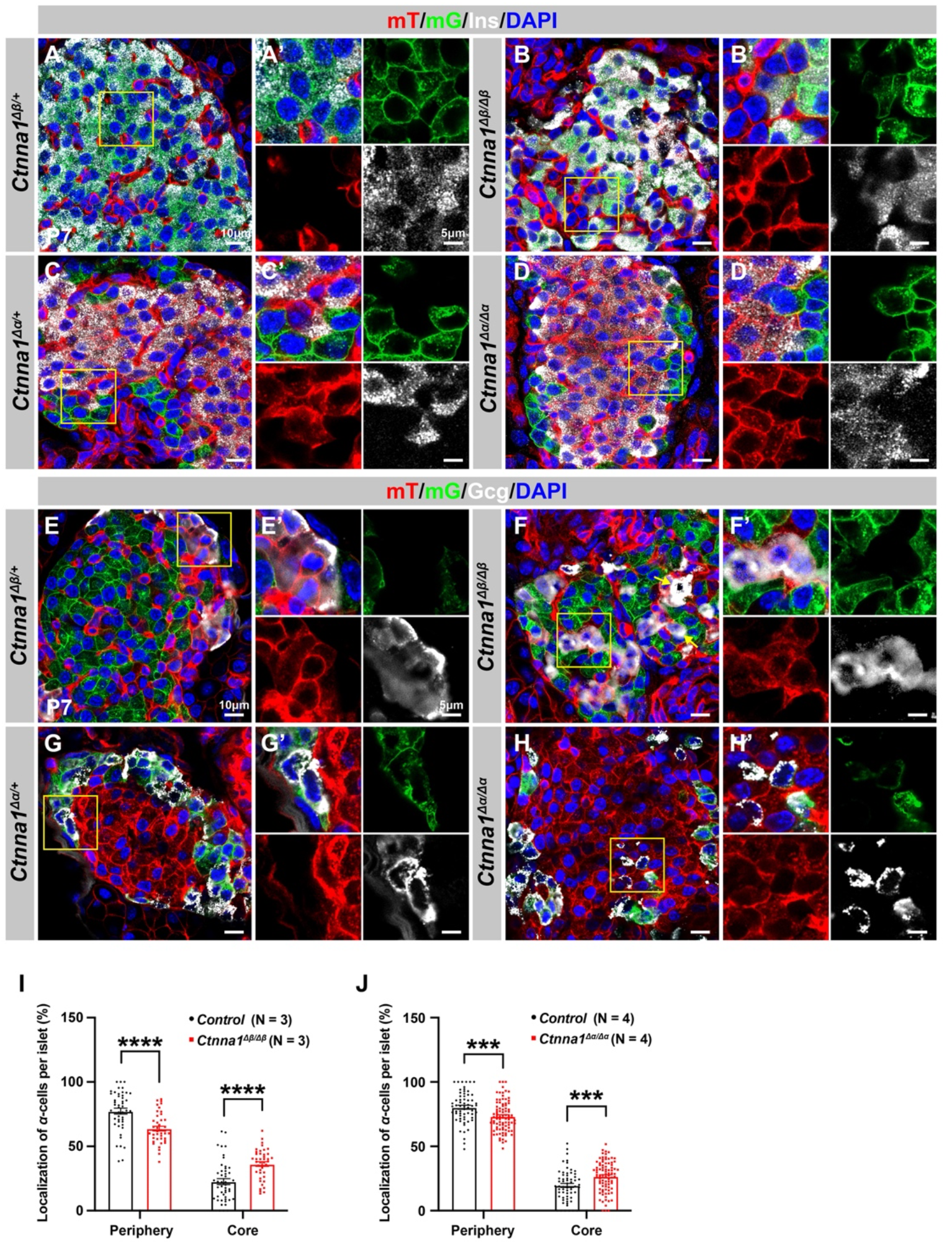
Loss of Ctnna1 in α- and β-cells leads to disrupted islet architecture. (A-H) Airyscan images of immunofluorescence staining for Ins (A-D), Gcg (E-H), and DAPI in pancreas sections of P7 mT/mG reporter mice: Heterozygous *Ctnna1^Δ*β*/+^* (A,E), homozygous *Ctnna1^Δ*β*/Δ*β*^* (B,F), heterozygous *Ctnna1^Δα/+^* (C,G), and homozygous *Ctnna1^Δα/Δα^* (D,H). The yellow arrows in (F) showing α-cells in the islet core in *Ctnna1^Δ*β*/Δ*β*^* mice. The fields demarcated by yellow boxes are shown at higher magnification with individual color channels in the side panels. (I-J) Quantification of α-cell localization in the islets of *Ctnna1^Δ*β*/Δ*β*^* (I) and *Ctnna1^Δα/Δα^* (J) mice. Compared to controls, *Ctnna1^Δ*β*/Δ*β*^* and *Ctnna1^Δα/Δα^* mice exhibit a reduction in peripherally located α-cells and an increase in core-located α-cells. These results suggest that the loss of Ctnna1 in α- and β-cells affects the organization and localization of α-cells within the islet. Gcg, Glucagon; DAPI, 4’,6-diamidino-2-phenylindole; Ins, Insulin; P7, postnatal day 7 Data are shown as mean ± SEM. *** p < 0.001, **** p < 0.0001.

Based on the DAH proposed by previous studies, we expected that Ctnna1-expressing α-cells would tend to cluster together and become more concentrated towards the center of *Ctnna1^Δβ/Δβ^*islets due to weaker adhesion in β-cells lacking Ctnna1. Our observations supported this hypothesis: in *Ctnna1^Δβ/Δβ^* islets, where only β-cell adhesion was affected but not α-cell adhesion, we observed a significant decrease in α-cells at the periphery and an increase in their density at the core of the islets (Figure 5B, 5F, 5I). Additionally, our data revealed that islet core-located α-cells had a higher tendency to form clusters (Figure 5F). These results are consistent with the prediction of DAH that cells with stronger adhesion tend to exclude those with weaker adhesion and to form a cluster with a defined size.

However, in α-cell-specific Ctnna1-deletion (*Ctnna1^Δα/Δα^*) mutants, we observed a reduction in α-cells at the periphery and an increase in their frequency in the islet core (Figure 5D, 5H, 5J), which was unexpected based on the predictions of DAH. Nevertheless, the core-located Ctnna1-deficient α-cells were mostly scattered as single cells without clustering together in the islet of *Ctnna1^Δα/Δα^* mice (Figure 5H). This suggests that the role of Ctnna1 in regulating cellular distribution within the islets may be more complex than the simple attractor-repeller mechanism proposed by DAH. These observations support the idea that differential adhesion between endocrine subtypes is a contributing factor for establishing the islet architecture during development; and the weaker cell adhesion in the peripheral islet layers leads to the specific loss of these cell types in *Ctnna1*^Δ*Endo/*Δ*Endo*^ islets.

### Coordination of ECM and cell-cell adhesion regulates islet vascularization, architecture, and functional maturation

Our studies have shown that the coordination of cell-ECM and cell-cell adhesion plays a vital role in regulating islet development. These adhesions work together to balance interactions between endocrine cells and blood vessels. When one of these adhesions is perturbed, islet endocrine cells may compensate with another interaction to maintain islet aggregation (Figure S5I). However, it is still unclear whether islet clustering relies solely on these two adhesions. To test this, we generated *Ngn3-Cre; Ctnna1^f/f^; Itgb1^f/f^* (hereafter *Ctnna1; Itgb1*^Δ*Endo/*Δ*Endo*^) double knockout (DKO) mice, in which we simultaneously removed cell-ECM and cell-cell adhesion in all endocrine progenitor cells. We compared control, *Ctnna1*^Δ*Endo/*Δ*Endo*^*, Itgb1*^Δ*Endo/*Δ*Endo*^ and DKO islets at P7 to examine their morphology.

At P7, *Ctnna1*^Δ*Endo/*Δ*Endo*^ islets displayed loosely aggregated endocrine cells, increased vascularization (Figure 6A, 6B), reduced F-actin assemblies, and more active-Itgb1^+^ endothelial cells in the loosened islet aggregates (Figure S8A, S8B). Conversely, *Itgb1*^Δ*Endo/*Δ*Endo*^ islets clumped together (Figure 6C), with strong F-actin assemblies and less active-Itgb1^+^ endothelial cells in the tightly aggregated islets (Figure S8C). However, the aggregation of endocrine cells was completely abrogated in the DKO mice (Figure 6D). These DKO endocrine cells became suspended single cells scattered throughout the entire pancreatic epithelium, displaying rounded cell morphology (Figure 6D). The DKO endocrine cells were larger and displayed weak F-actin assemblies (Figure S8D). Despite the drastic changes in endocrine cell aggregation and morphology, the endocrine cell subtypes were still present in the P7 DKO islets (Figure 6D and data not shown), suggesting that the differentiation of endocrine subtypes is independent of the islet aggregation process.

**Figure 6.**
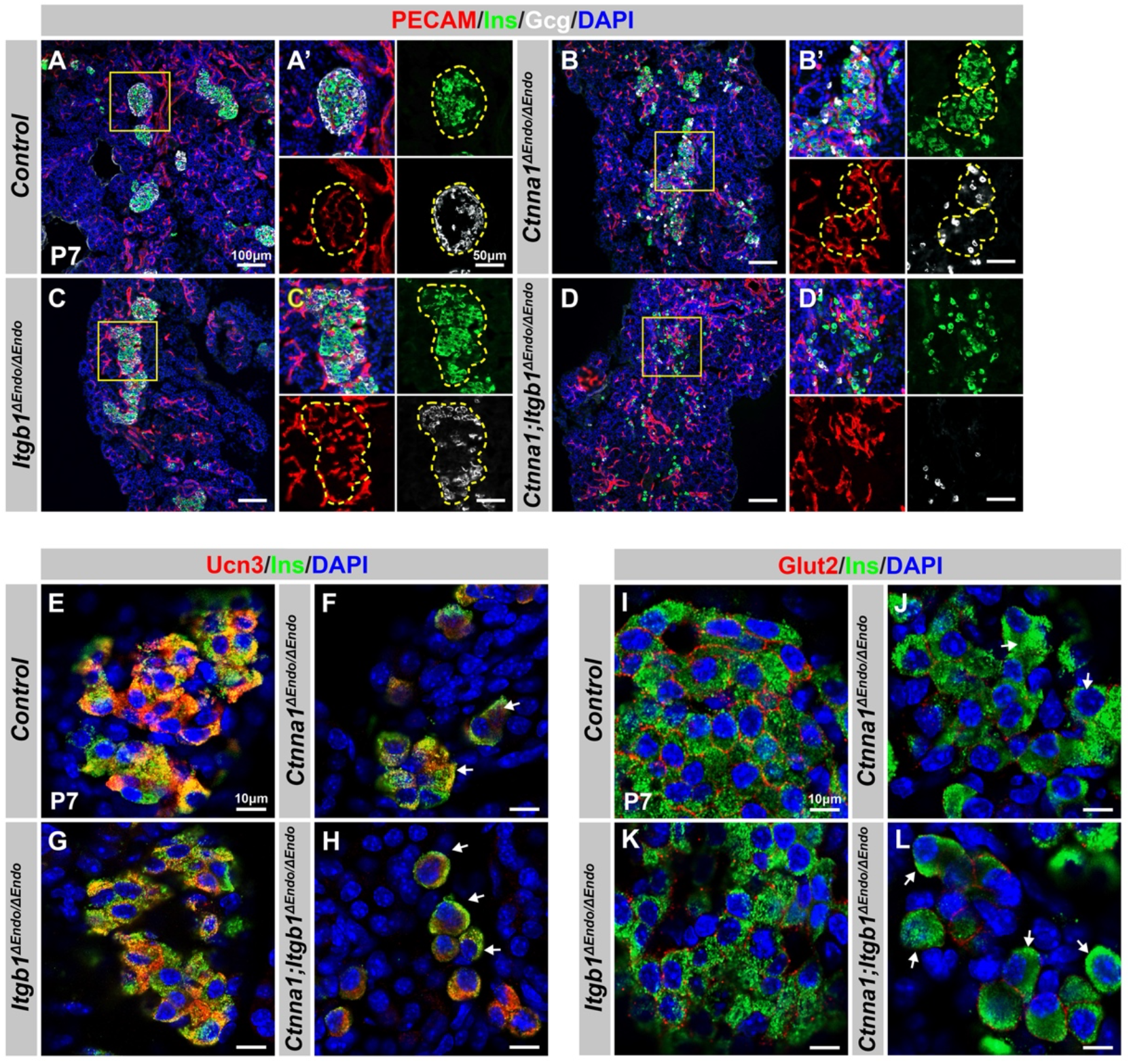
Disruption of cell-ECM and/or cell-cell adhesion results in abnormal islet vascular architecture, endocrine cell aggregation, and decreased expression of β-Cell maturation markers at postnatal day 7. (A-D) Immunofluorescence staining of PECAM, Ins, Gcg, and DAPI in P7 (A) control, (B) *Ctnna1^ΔEndo/ΔEndo^*, (C) *Itgb1^ΔEndo/ΔEndo^*, and (D) *Ctnna1; Itgb1^ΔEndo/ΔEndo^* mice. Fields demarcated by yellow boxes in A-D are shown at higher magnification in A’-D’ side panels. Individual islet shape is outlined by dashed yellow lines in A’-C’. Endocrine cells are suspended and scattered throughout the pancreas of *Ctnna1; Itgb1^ΔEndo/ΔEndo^* mice. (E-L) Airyscan images of immunofluorescence staining for Ucn3, Glut2, Ins, and DAPI in P7 (E,I) control, (F,J) *Ctnna1^ΔEndo/ΔEndo^*, (G,K) *Itgb1^ΔEndo/ΔEndo^*, and (H,L) *Ctnna1; Itgb1^ΔEndo/ΔEndo^* mice. Arrows indicate reduced expression of Ucn3 or Glut2 in *β*-cells of the mutant mice. PECAM, Platelet endothelial cell adhesion molecule-1; Ins, Insulin; Gcg, Glucagon; DAPI, 4’,6-diamidino-2-phenylindole; Ucn3, Urocortin 3; Glut2, Slc2a2; P7, postnatal day 7

We next investigated whether the failure in islet aggregation affected endocrine cell maturation by performing IF for β-cell maturation markers. In P7 control mice, Ucn3 and Glut2 were robustly expressed in the cytosol and cell surface of β-cells, respectively (Figure 6E, 6I). Deletion of *Ctnna1* or *Itgb1* in endocrine progenitor cells led to reduced expression of Ucn3 and Glut2 in β- cells (Figure 6F, 6G, 6J, 6K), which was consistent with gene profiling analysis from 8 week old mice. Finally, DKO islets displayed a severe loss of β-cell maturation markers in P7 β-cells (Figure 6H, 6L). In addition, while endocrine cells in normal islets do not express progenitor markers Sox9 or Spp1 at P7 (Figure S8E, S8G), the endocrine cells in the DKO islets exhibited elevated expression of these markers at P7 (Figure S8F, S8H). These findings suggest that β-cell maturation is linked to the cell adhesion-mediated islet aggregation process.

To investigate whether the islet aggregation and β-cell maturation defects in DKO mice at P7 were due to developmental delays, we examined islets during later adulthood stages. At 8 weeks, normal endocrine cells form tightly aggregated islets (Figure 7A), but in DKO mice, endocrine cells remain mostly scattered, forming smaller and loosely attached clusters (Figure 7B). These smaller clusters of endocrine cells abnormally express progenitor markers Sox9 and Spp1 (Figure 7B; and data not shown). While the endocrine cells appear to be located closely to each other in lower resolution images, Airyscan super-resolution images of F-actin IF on endocrine cells in DKO islets showed that the endocrine cells are completely separated from each other without forming discernible junction structures (Figure 7C, 7D, yellow arrows). Furthermore, the endocrine cells also exhibit separation from the vasculatures in the DKO islets (Figure 7C, 7D, red arrows). These observations indicate that both the islet aggregation and the β-cell maturation defects in DKO mice persist into adulthood. Supporting this, low expression of β-cell maturation markers Ucn3 and Glut2 persisted in 8 week old DKO islets (Figure 7F-7H). These mice were severely glucose intolerant (Figure 7I) and exhibit very low insulin secretion upon glucose stimulation (Figure 7J). Together, our findings support a model in which islet morphogenesis, vasculature interaction, endocrine cell aggregation, and the β-cell maturation process depend on cell-ECM and cell-cell adhesion during development (Figure 7K).

**Figure 7.**
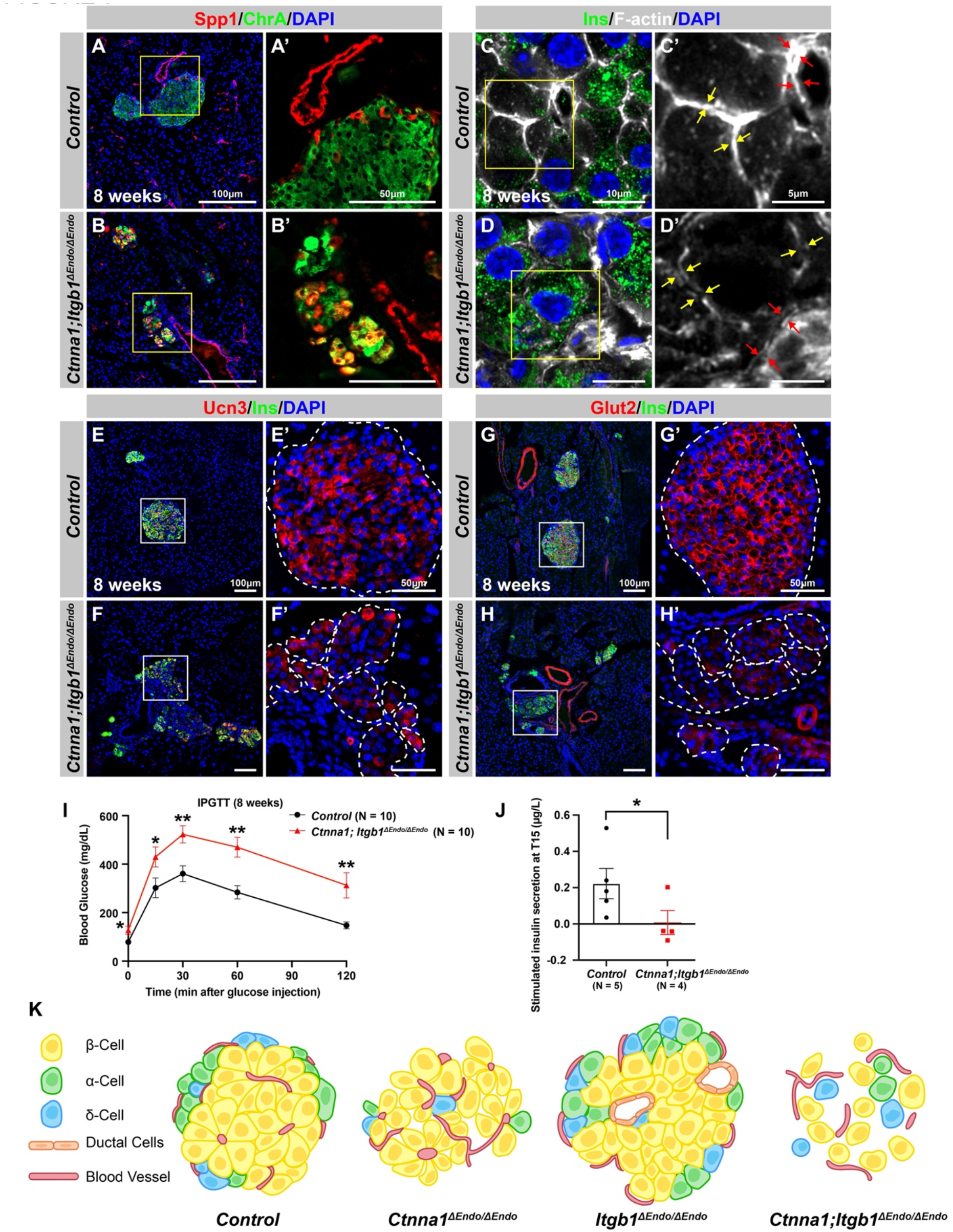
Abnormal islet aggregation, mis-regulation of β-Cell maturation markers, and insulin secretion defects persist in *Ctnna1; Itgb1^ΔEndo/ΔEndo^*mice into adulthood. (A-B) Immunofluorescence staining for endocrine cell marker ChrA and progenitor marker Spp1 on pancreas sections from control (A) and *Ctnna1; Itgb1^ΔEndo/ΔEndo^*(B) mice at 8 weeks old. The yellow boxes in A,B indicate the fields shown at higher magnification in A’,B’ side panels. In the islets of *Ctnna1; Itgb1^ΔEndo/ΔEndo^*mice, a subpopulation of ChrA^+^ endocrine cells expresses progenitor markers Spp1 (yellow cells in B and B’). (C-D) Airyscan images of immunofluorescence staining for Ins, F-actin and DAPI in pancreatic sections from 8-week-old control and *Ctnna1; Itgb1^ΔEndo/ΔEndo^*mice are shown. In control islets (C,C’), the F-actin assemblies appear aligned with cell membranes and condensed in cellular junctions, as indicated by the yellow and red arrows, respectively. In contrast, diffused distribution of F-actin (red arrows in D’) and separation of F-actin assemblies (yellow arrows in D’) between Ins^+^ β-cells are observed in the *Ctnna1; Itgb1^ΔEndo/ΔEndo^* mice. (E-F) Immunofluorescence staining of Ucn3, Ins, and DAPI in pancreatic sections of 8 weeks old (E) control and (F) *Ctnna1; Itgb1^ΔEndo/ΔEndo^* mice. Fields demarcated by white boxes in E and F are shown at higher magnification in E’ and F’, and individual islet shape is outlined by dashed white lines. The reduction of Ucn3 expression persists in adult *Ctnna1; Itgb1^ΔEndo/ΔEndo^* mice. (G-H) Immunofluorescence staining of Glut2, Ins, and DAPI in 8 weeks old (G) control and (H) *Ctnna1; Itgb1^ΔEndo/ΔEndo^* mouse pancreas. Fields demarcated by white boxes in G and H are shown at higher magnification in G’ and H’, and individual islet shape is outlined by dashed white lines. The reduction of Glut2 expression persists in adult *Ctnna1; Itgb1^ΔEndo/ΔEndo^* mice. (I-J) Intraperitoneal glucose tolerance test was performed on 8-week-old mice (I), and glucose-stimulated insulin secretion was measured at T15 in 20-38-week-old mice (J) from control and *Ctnna1; Itgb1^ΔEndo/ΔEndo^* groups. *Ctnna1; Itgb1^ΔEndo/ΔEndo^* mice exhibited severe glucose intolerance and defects in insulin secretion. (K) Graphical summary of the islet phenotypes observed after disruption of cell-cell and cell-ECM adhesion. Ins, Insulin; ChrA, Chromogranin A; Spp1, Secreted Phosphoprotein 1; Ucn3, Urocortin 3; IPGTT, intraperitoneal glucose tolerance test Data are shown as mean ± SEM. * p < 0.05 and ** p < 0.01.

## DISCUSSION

Our study reveals the combinatorial roles of cell-ECM and cell-cell adhesion in regulating the aggregation of endocrine cells into islets. Mechanistically, islet cell aggregation depends on endocrine cell-cell adhesion, which is negatively regulated by cell-ECM adhesion through vascular interaction. These two adhesion properties are reciprocally regulated: lowering cell-ECM adhesion promotes endocrine cell-cell adhesion; and, conversely, lowering cell-cell adhesion promotes cell-ECM adhesion. Importantly, the aggregation and functional maturation processes of endocrine cells are affected by the loss of either adhesion property and are further affected by the loss of both.

In this study, we aimed to elucidate the role of ECM signaling by specifically deleting the essential ECM receptor Itgb1 during endocrine specification. Our findings demonstrate that while the morphology and function of *Itgb1^ΔEndo/ΔEndo^* islets were impacted, the total islet area remained unaffected, suggesting that Itgb1 deletion may not result in overt cell death in islets, unlike in other organ systems (Carlson et al., 2008; Speicher et al., 2014). These observations indicate the involvement of other mediators of cell-ECM interactions in islet maintenance. Although Itgb1 is a key player in cell-ECM interactions, compensatory mechanisms involving other integrins, such as β3, β4, β5, β 6; and α3, α 6, αV integrins, may be operative in Itgb1-deficient islets. Previous studies have highlighted the critical roles of these integrins in cell-ECM interactions and islet biology (Cirulli et al., 2000; Krishnamurthy et al., 2011; Schiesser et al., 2021; Yashpal et al., 2005). It is plausible that these integrins compensate for cell-ECM interactions, ensuring the survival and function of Itgb1-deficient islets. Further investigations are warranted to comprehensively understand the compensatory mechanisms and contributions of other integrins in Itgb1-deficient islets.

Both *Itgb1^ΔEndo/ΔEndo^* and *Itgb1*^Δ*Matureβ/*Δ*Matureβ*^ mice demonstrate reduced vascularization, highlighting the importance of ECM signaling in vascular interactions throughout stages from embryonic endocrine cells to mature β-cells. This aligns with previous research using *MIP-Cre* to create *Itgb1*-knockout mice, which also exhibited decreased islet vascularization (Win et al., 2020). The critical role of Itgb1 signaling in organ vascularization seems universal; studies indicate that Itgb1-inactivation in pituitary endocrine epithelial cells and kidney cells leads to a loss of vasculature (Mohamed et al., 2020; Scully et al., 2016). Reduced vascular interaction can instigate hypoxia responses, which have been tied to β-cell maturation processes (Balboa et al., 2022; Heinis et al., 2010; Zeng et al., 2017). We thus propose that the impaired β-cell maturation observed in *Itgb1^ΔEndo/ΔEndo^* mice could stem from a hypoxic state caused by vascular loss. However, this hypothesis requires further empirical investigation for validation.

The expression of *Vegfa*, an essential growth factor for islet vasculature, is not affected in *Itgb1^ΔEndo/ΔEndo^* islets (Table S1). Thus, the reduced vascularization in Itgb1-deficient islets may not be explained by the lack of Vegfa. Downstream of Integrin signaling, integrin-linked kinase (ILK) is activated upon Integrin-ECM binding to regulate actin organization, cell migration, and other cellular processes (Hannigan et al., 2005). Genetic deletion of ILK in the developing pancreas leads to the failure of ILK-inactivated endocrine cells to adhere to the basement membrane, and drastic reduction of intra-islet blood vessel density (Kragl et al., 2016). This study further showed that cortical actomyosin contraction was significantly increased, thus increasing cortex tension in both the ILK-deleted islets, as well as in Itgb1-inactivated cultured β-cells. Our analyses showed aberrant organization of F-actin fibers and compaction of individual endocrine cells, as well as dysregulated cytoskeleton genes in *Itgb1*-deleted islets. Thus, it is likely that both the *Itgb1^ΔEndo/ΔEndo^* and *Itgb1*^Δ*Matureβ/*Δ*Matureβ*^ islets fail to interact with vasculature resulting in the same alteration of the actomyosin regulation seen in ILK-deleted islet cells. In contrast, the *Ctnna1^ΔEndo/ΔEndo^*model shows increased islet vascularization without affecting the expression of *Vegfa* (Table S2). We posit that weakened adhesion between endocrine cells in *Ctnna1^ΔEndo/ΔEndo^*islets may allow greater penetration of vasculature through islet clusters, leading to augmented islet vascularization. Supporting this notion, salivary gland patterning allows vessel penetration via an epithelial “cleft” created by weakened cell-cell adhesion (Kwon et al., 2017). Interestingly, the initial steps of cleft formation requires ECM-Itgb1 signaling (Sakai et al., 2003); conversely, reducing cell-cell adhesion promotes cell-ECM adhesion during budding morphogenesis (Wang et al., 2021). Thus, our proposed model, in which ECM signaling in concert with cell-cell adhesion controls islet aggregation, may reflect a fundamental mechanism of organ morphogenesis.

In the context of our findings, it is crucial to consider the potential role of vascular alignment in islet cell function. The vasculature system within the islet plays an indispensable role not only in delivering nutrients to the cells but also in facilitating the efficient distribution of hormones into the peripheral circulation (Gan et al., 2018). The functional maturation of β-cells and their efficiency in hormone secretion are likely influenced by their proximity to and interaction with the vasculature. In our *Itgb1* and *Ctnna1* deficient models, we observed an abnormal aggregation of islet cells, resulting in scattered cells that appear misaligned with the vasculature. This disorganization could potentially impair nutrient delivery and hormone distribution, further exacerbating the functional impairments seen in these models. It is plausible that, in addition to their impact on cell-cell adhesion and aggregation, the inactivation of Itgb1 and Ctnna1 may also interfere with the proper integration of β-cells with the islet vasculature. This perspective adds another layer to our understanding of the complex interplay between cellular adhesion, aggregation, and the vasculature in islet morphology and function.

The onset of the proendocrine cell program triggers the expression of Ngn3, a factor thought to stimulate the activation of ECM signaling. This signaling pathway is regarded as a key regulatory mechanism controlling the delamination and migration of proendocrine cells away from ductal progenitor cords (Rosenberg et al., 2010). In the case of *Itgb1^ΔEndo/ΔEndo^* mice, the deactivation of ECM signaling in these endocrine progenitors leads to a noticeable decrease in their migration distances. Consequently, differentiated endocrine cells remain entangled with ducts post-differentiation from as early as E15.5, highlighting the role of ECM signaling in facilitating endocrine progenitor migration during initial developmental stages. Interestingly, this phenomenon of reduced migration is not observed in *Itgb1^ΔMatureβ/ΔMatureβ^* islets, suggesting the impact of ECM signaling on cell migration may be more specific to the progenitor stage. Supporting this hypothesis, studies utilizing *Ins-Cre* or *RIP-Cre* to ablate Itgb1 in post-differentiated insulin-producing cells have reported no discernible migration defects (Diaferia et al., 2013; Win et al., 2020). During the limited time window of endocrine cell development, islet cell radial migration is also modulated by semaphorin signaling (Pauerstein et al., 2017). The remodeling of actin, induced by semaphorin, is associated with changes in the cell’s anchorage to the ECM (Alto and Terman, 2017). This suggests that the chemoattractant system, where semaphorin signaling occurs, could potentially intersect with the activation of Itgb1 signaling. This intersection of signaling pathways may guide the migration of islet progenitor cells toward the periphery and away from the ducts. Thus, the chemoattractant system where semaphorin signaling occurs may involve the activation of Itgb1 signaling to control islet progenitor cell migration toward the periphery, and away from the ducts. Whether the migration of endocrine progenitors toward the periphery affects their function, remains to be tested. However, the role of ECM-Integrin signaling extends beyond the scope of cell migration. In adult stages, ECM-Integrin signaling continues to have a significant impact on differentiated endocrine cells within pancreatic islets (Gan et al., 2018). Therefore, when we use the term “endocrine cells” in this study, we refer to both progenitor and differentiated cells, and the significance of ECM-Integrin signaling spans across multiple cellular processes, not being confined to migration alone.

Despite the endocrine subtype ratios being different in human versus rodent islets, the precise differences in the stereotypical architectures in the two species have been debated. Studies using different imaging modalities have yielded different conclusions (Dybala et al., 2020). In mice, it is proposed that spatiotemporal collinearity leads to the typical core-mantle architecture of the spherical islet in which α-cells, the first to develop, form the peninsular outer layer, and β-cells subsequently form beneath them (Sharon et al., 2019a). Our current study has not directly tested this hypothesis, yet our observations suggest that differential cell adhesion is critical for developing this structure. Our data support a model in which randomly allocated α- and β-cells are specified and sorted into the final islet architecture based on differential adhesion. It remains to be tested whether the differential cell adhesion is also involved in the development of islet architectures in human. In addition, studies have shown that the Roundabout (Robo)/Slit signaling axis plays a role in determining islet stereotypical architecture. The secreted ligand, Slit, is often associated with the ECM, and binds to Robo receptors to control various cellular responses via cytoplasmic kinases and the regulation of actin and microtubule cytoskeleton (Jiang et al., 2022; Tong et al., 2019). Mice lacking *Robo1* and *Robo2* in all endocrine cells, or selectively in β cells, show complete loss of endocrine cell type sorting in the islets (Adams et al., 2021; Adams et al., 2018; Gilbert et al., 2020). Interestingly, Robo1/2 double knockout islets exhibit adhesion defects and are prone to dissociate upon isolation (Adams et al., 2018), similar to the *Ctnna1^ΔEndo/ΔEndo^* islets. Similar observations have been shown in stem cell sorting and cancer cell migration; binding of Slit to Robo receptors leads to the modulation of Ncad- or Ecad-mediated cell adhesion (Stine et al., 2014; Tong et al., 2019). We speculate that Robo/Slit signaling regulates endocrine cell type sorting in islets by controlling cell adhesive properties to establish the islet architecture.

### Limitation of the study

Our study’s potential limitation is the use of the *Ngn3-Cre* driver for targeted gene deletion. While commonly used for pancreatic endocrine progenitor and its lineage-specific gene deletion, *Ngn3* is also expressed in other tissues such as the brain and gastrointestinal tract (Schonhoff et al., 2004; Simon-Areces et al., 2010). Deletion of genes using this driver could impact these other tissues and create unintended consequences. Previous studies show that *Ctnna1* deletion in neuronal progenitor cells in the brain can lead to altered neural cell proliferation and differentiation (Lien et al., 2006). Although we did not investigate the potential effects of *Itgb1* and *Ctnna1* deletion in these other tissues, further studies are necessary to comprehend the role of ECM signaling and cell adhesion in these other tissues and the potential consequences of their deletion using the *Ngn3-Cre* driver.

We acknowledge the intricate interplay between cell-ECM and cell-cell interactions and understand that the loss of one type of interaction does not universally enhance the other. This complexity was exemplified in our double knockout model, which did not exhibit the expected compensatory behaviors. While our *in vitro* model provided some insight, we recognize its limitations in fully mirroring the dynamic and sophisticated changes occurring *in vivo*. Consequently, our observations serve as preliminary findings that pave the way for more comprehensive investigation. Further research, particularly in more complex, dynamic models, will be vital in elucidating the precise nature of these interactions and their role in cell development and function. The dynamic nature of islet aggregation and endocrine subtype sorting would require more in-depth live imaging analysis. Obtaining 3D cellular information in developing islets would provide insights into these dynamic and complex phenomena, and further our understanding of the process of islet development and morphogenesis. We anticipate that using pancreas slice cultures (Panzer et al., 2020) to develop an imaging platform to monitor the development of live neonatal islets would provide a framework for understanding how islets develop, and offer novel ways to harness this understanding in the search for clinical alternatives to treat diabetes.

## AUTHOR CONTRIBUTIONS

W.T., M.M., Y.T.C., W.Y. and H.P.S. conceived the project, designed the experiments, and analyzed the data. W.T., M.M., Y.T.C. performed the analysis of mouse genetic models and live-imaging experiments. Y.T.C performed *in vitro* islet cell aggregation assay. A.C. assembled figures and graphical summaries. N.P. and S.D. analyzed the RNA-seq data and generated heatmaps. M.N. performed segmentation analysis. P.F. performed hyperglycemic clamp experiment. R.H and M.H. analyzed *Ucn3-Cre* mouse model and prepared gene expression genome viewer plots. S.W., W.J.H., and J.L.K. conceived *Ctnna1-flox* mouse breeding and experiments. W.T., M.M., and H.P.S. wrote the manuscript.

## Supporting information

Key Resource Table

List of n and statistics

provided Source Data spreadsheet

## ACKNOWLEDGEMENTS

We acknowledge the support of City of Hope Core Facilities: Dr. Brain Armstrong from the Microscopy Core, Dr. Zhuo Li from the Electron Microscopy Core, Drs. Xiwei Wu and Min-Hsuan Chen from the Integrative Genomics Core, Dr. Patrick Fueger from the Comprehensive Metabolic Phenotyping Core at City of Hope. We thank Dr. Maike Sander for sharing the anti-Ngn3 antiserum and Dr. Rupangi Vasavada for sharing the *Ins-Cre* mouse strain. The California Institute for Regenerative Medicine (CIRM) Training Grant EDUC4-12772 to W.T.; JDRF postdoctoral fellowship 3-PDF-2019-742-A-N to M.M.; NSERC Discovery Grant (RGPIN-2016-04276), CIHR New Investigator Award Msh-147794, and MSFHR Scholar Award 18309 to J.L.K., NIH/NIDDK 1R01DK110276 to M.O.H.; NIH/NIDDK 1R01DK120523, Human Islet Research Network (HIRN) New investigator Award (subaward via UC4DK104162), and a grant from the Wanek Family Foundation to Cure Type 1 Diabetes to S.D.; Wanek Family Foundation to Cure Type 1 Diabetes and NIH/NIDDK 1R01DK119590 to H.P.S.. We acknowledge the assistance of OpenAI’s ChatGPT in refining the language and grammar of our manuscript during resubmission. The contributions of this AI model were strictly limited to textual improvements, with no role in generating or modifying the scientific content and findings of the study.

---

**Figure S1.**
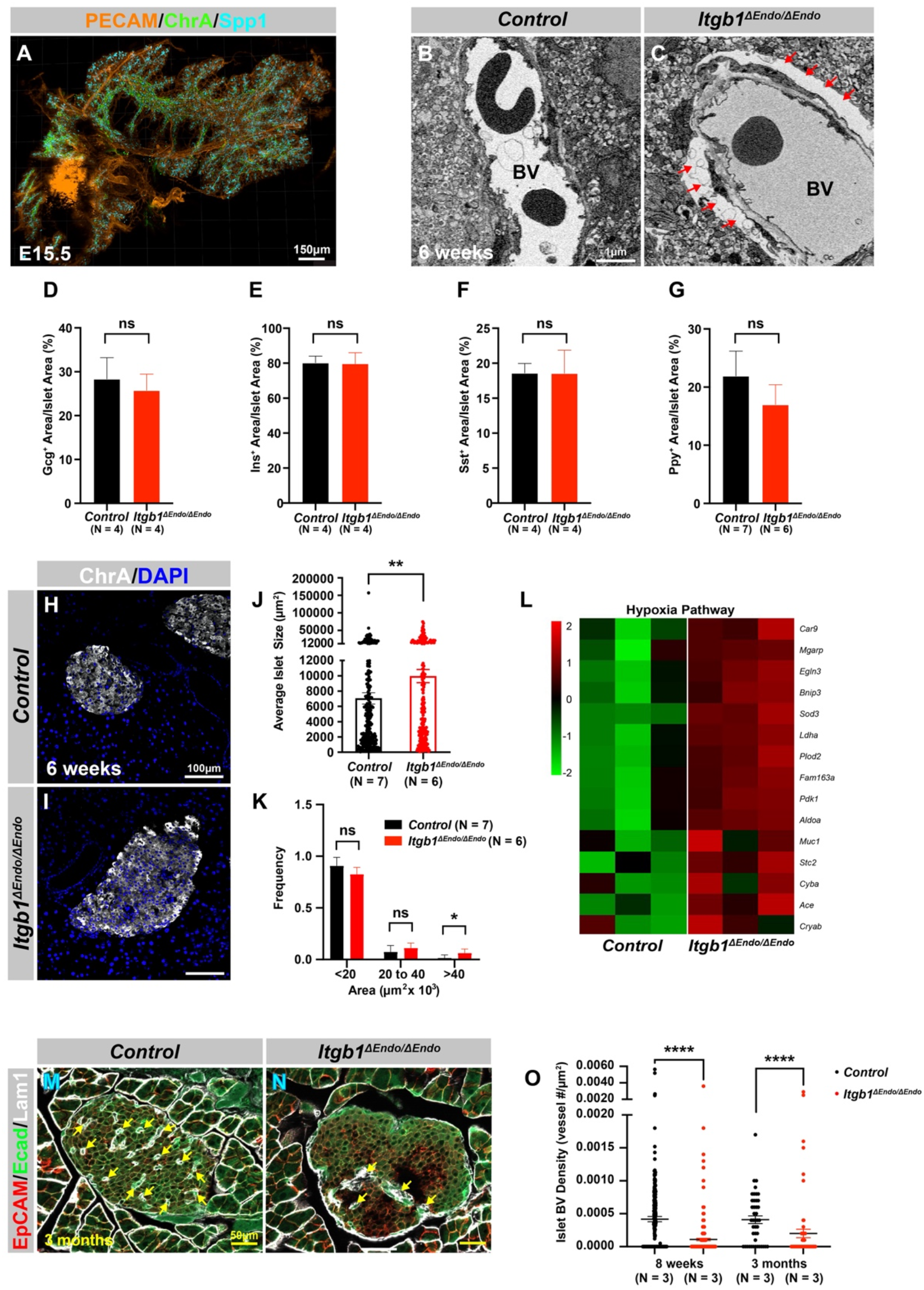
Deletion of *Itgb1* in endocrine progenitors results in abnormal interactions with blood vessels and decreased expression of functional maturation markers within islets. Related to Figure 1. (A) Whole mount immunofluorescence staining for PECAM, ChrA, and Spp1 in a wild-type pancreas at E15.5. (B-C) Transmission electron microscope images showing blood vessels in 6 weeks old islets. Deletion of *Itgb1* disrupts the attachment of endocrine cells to blood vessels (red arrows) in *Itgb1^ΔEndo/ΔEndo^* islets. (D-G) Quantification of the percentage of (D) Gcg^+^, (E) Ins^+^, (F) Sst^+^, and (G) Ppy^+^ cells relative to measured islet area of 6 weeks old control and *Itgb1^ΔEndo/ΔEndo^* pancreas sections. (H-I) Immunofluorescence staining for ChrA, and DAPI in 6 weeks old (H) control and (I) *Itgb1^ΔEndo/ΔEndo^* pancreas sections. (J-K) Quantification of (J) average islet size and (K) islet size distribution based on ChrA staining of 6 weeks old control and *Itgb1^ΔEndo/ΔEndo^* pancreas sections. (L) Heatmap showing hypoxia pathway genes are up-regulated in *Itgb1^ΔEndo/ΔEndo^* islets. (M-N) Immunofluorescence staining for EpCAM, Ecad, and Lam1 on 3 months old pancreas sections showing reduced islet vasculature indicated by Lam1^+^ cells (yellow arrows). (O) Quantification of islet blood vessel density in control or *Itgb1^ΔEndo/ΔEndo^*islet area at 8 weeks and 3 months old. PECAM, Platelet endothelial cell adhesion molecule-1; ChrA, Chromogranin A; Spp1, Secreted Phosphoprotein 1; Gcg, Glucagon; Ins, Insulin; Sst, Somatostatin; Ppy, Pancreatic Polypeptide; Ecad, E-cadherin; Epithelial cellular adhesion molecule, EpCAM; Lam1, Laminin1; BV, blood vessels; E15.5, embryonic day 15.5. Data are shown as mean ± SEM. * p < 0.05, ** p < 0.01, **** p < 0.0001.

**Figure S2.**
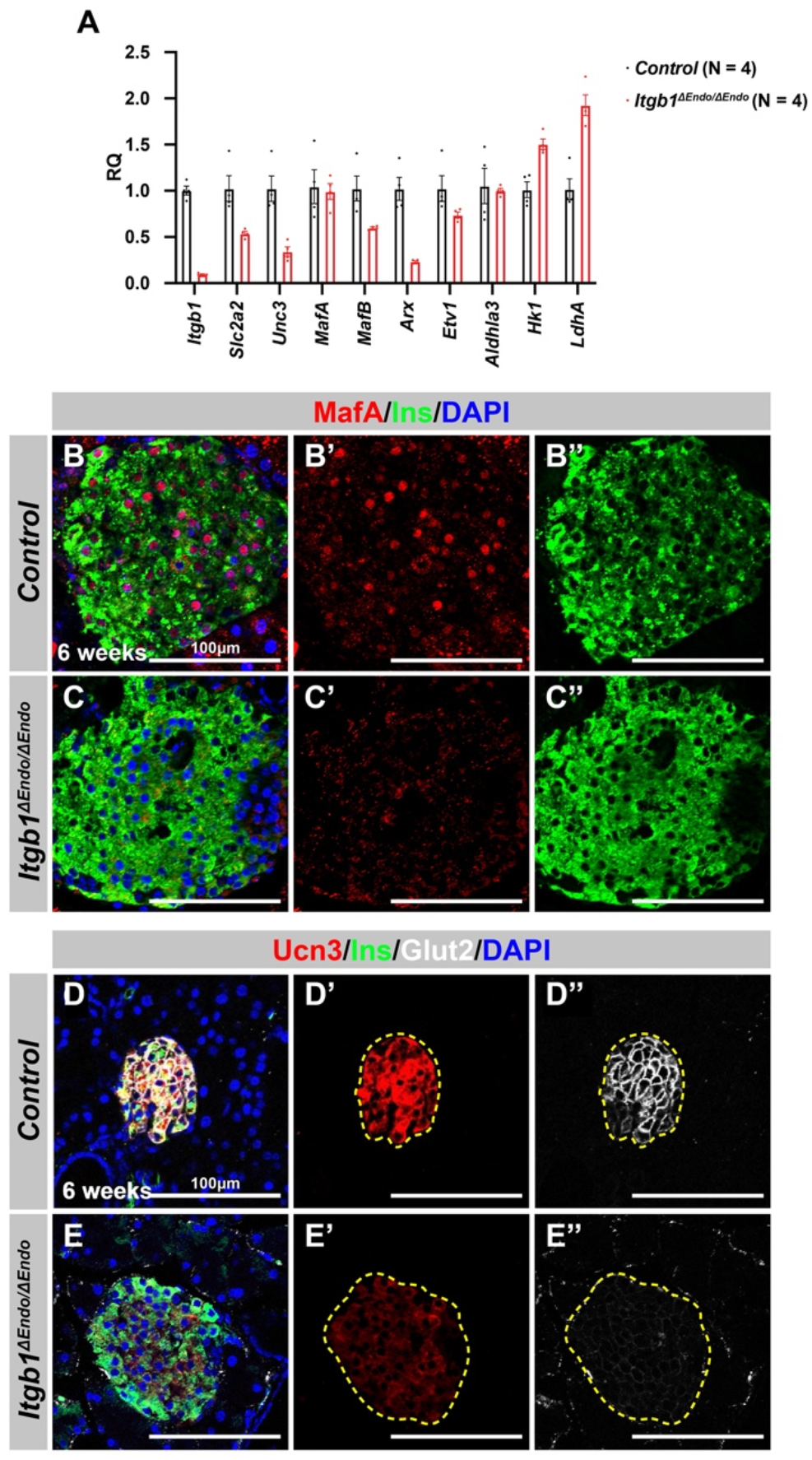
Deletion of *Itgb1* in endocrine progenitors results in abnormal interactions with blood vessels and decreased expression of functional maturation markers within islets. Related to Figure 1. (A) Quantification of selected differentially expressed genes involved in islet functional maturation using real-time qPCR analysis in *Itgb1^ΔEndo/ΔEndo^*islets and compared to controls. The relative quantification of gene expression levels was determined using the 2^-ΔΔCt^ method and is reported as a RQ value relative to the control group. The genes selected for validation were chosen based on their known roles in islet function and maturation. (B-E) Immunofluorescence staining for MafA, Ins, Ucn3, Glut2 and DAPI on 6 weeks old (B,D) control and (C,E) *Itgb1^ΔEndo/ΔEndo^* pancreas sections. Individual islets are outlined by dotted yellow lines in (D’,D”,E’,E”). RQ, Relative quantification; Ins, Insulin; Ucn3, Urocortin 3; PECAM, Platelet endothelial cell adhesion molecule-1; DAPI, 4’,6-diamidino-2-phenylindole; E15.5, embryonic day 15.5. Data are shown as mean ± SEM. * p < 0.05.

**Figure S3.**
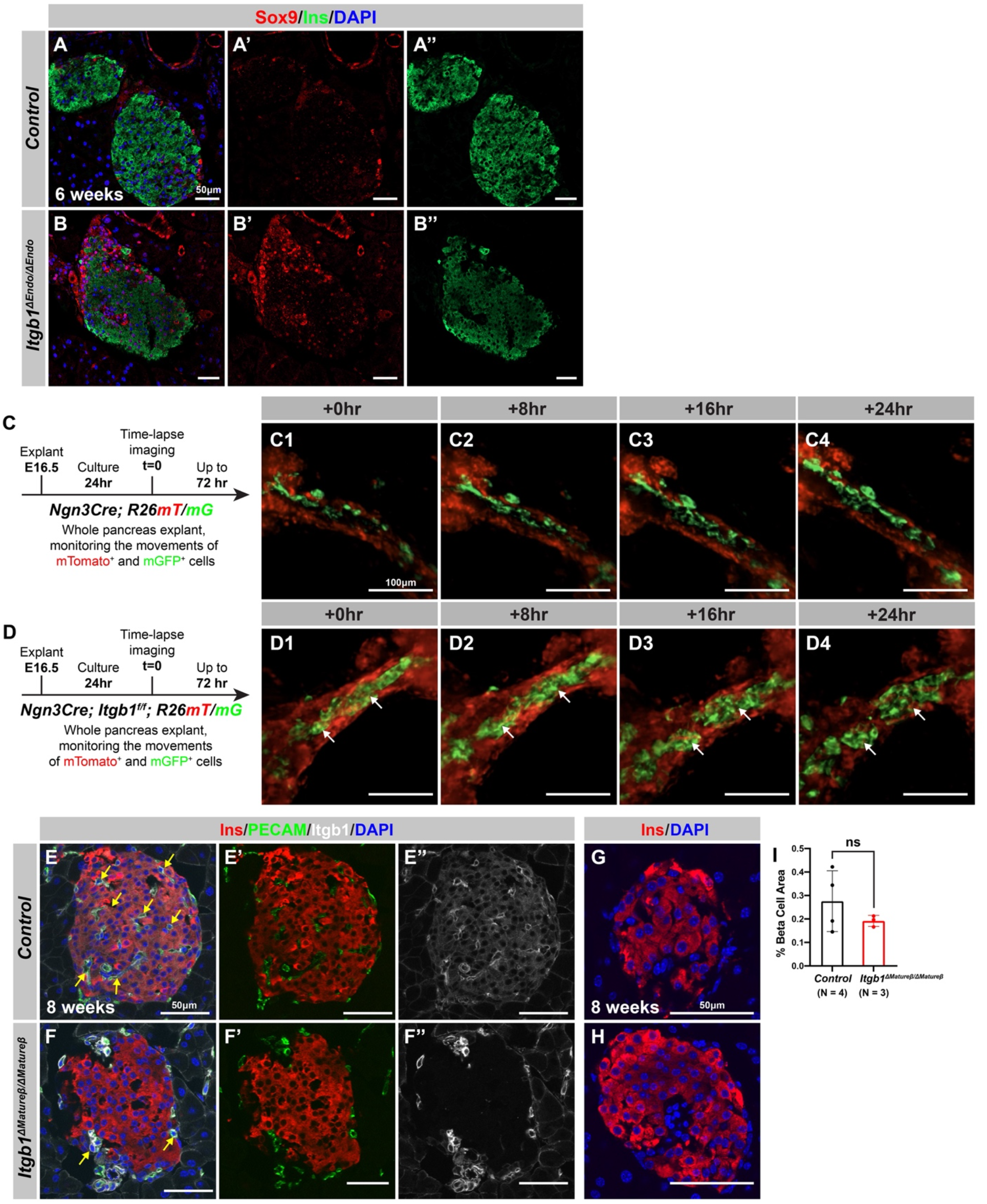
Itgb1 is required for normal islet architecture. Related to Figure 2. (A-B) Immunofluorescence staining for Sox9, Ins, and DAPI on 6 weeks old (A) control and (B) *Itgb1^ΔEndo/ΔEndo^* pancreas sections. The individual channels for Sox9 and Ins are displayed in panels A’ and A”, and B’ and B”, respectively. In the *Itgb1^ΔEndo/ΔEndo^* islets, Sox9-positive ductal cells were observed throughout the sections (B’). (C-D) Schematic of the time-lapse confocal microscope imaging of endocrine progenitor cell behaviors in E16.5 embryonic pancreas explants. The first 24 hours were monitored to observe cell movement in (C1-C4) control and (D1-D4) *Itgb1^ΔEndo/ΔEndo^*pancreas explants. White arrows indicate prematurely clustered islets in *Itgb1^ΔEndo/ΔEndo^*explants. (E-H) Immunofluorescence staining for Ins, PECAM, Itgb1, and DAPI on 8 weeks old (E,G) control and (F,H) *Itgb1^ΔMature*β*/ΔMature*β*^* pancreas sections. Individual channels are displayed as Ins/PECAM in E’,F’ and Itgb1 in E”,F”. In *Itgb1^ΔMature*β*/ΔMature*β*^* islets the presence of Itgb1 and vasculature in the islet is abolished. The overall size and insulin expression are not altered in the *Itgb1^ΔMature*β*/ΔMature*β*^* islets. (I) Quantification of total islet area based on morphometric analysis of 8 weeks old control and *Itgb1^ΔMature*β*/ΔMature*β*^* pancreas sections. Note no significant differences in islet size were observed between the two groups. Ins, Insulin; PECAM, Platelet endothelial cell adhesion molecule-1; DAPI, 4’,6-diamidino-2-phenylindole; E16.5, embryonic day 16.5 Data are shown as mean ± SEM.

**Figure S4.**
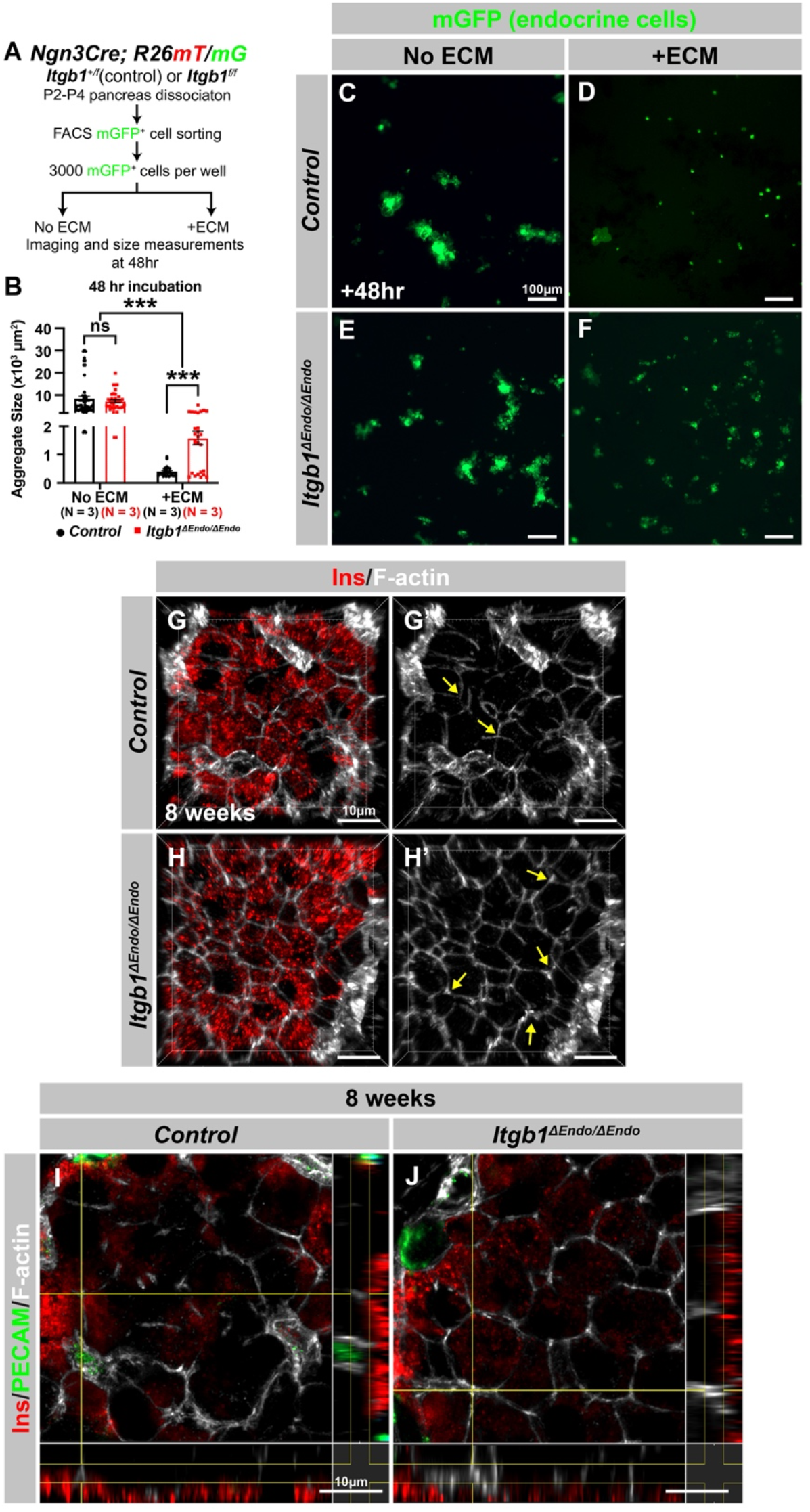
Itgb1 controls cytoskeleton regulation and organization in islets. Related to Figure 2. (A) Schematic of experimental design for *in vitro* aggregation assay. (B) Quantification of aggregate size after 48 hours incubation with or without ECM in control and *Itgb1^ΔEndo/ΔEndo^* endocrine progenitors. (C-F) Fluorescence images of mGFP^+^ endocrine progenitor aggregates after 48 hours culture. mGFP^+^ control endocrine progenitors clustered (C) without ECM but remained single cells (D) with ECM after 48 hours. Alternatively, *Itgb1^ΔEndo/ΔEndo^* endocrine progenitors clustered (E) without ECM or (F) with ECM after 48 hours. (G-J) Airyscan super-resolution 3D projection and cut-view images depict immunofluorescence staining for Ins and F-actin on 8 weeks old control and *Itgb1^ΔEndo/ΔEndo^* pancreas sections. The 3D images (G-H) and cut-view images (I-J) also demonstrated hexagonal cell shapes and increased F-actin assembly in the *Itgb1^ΔEndo/ΔEndo^* islets. Ins, Insulin; PECAM, Platelet endothelial cell adhesion molecule-1 Data are shown as mean ± SEM. *** p < 0.001.

**Figure S5.**
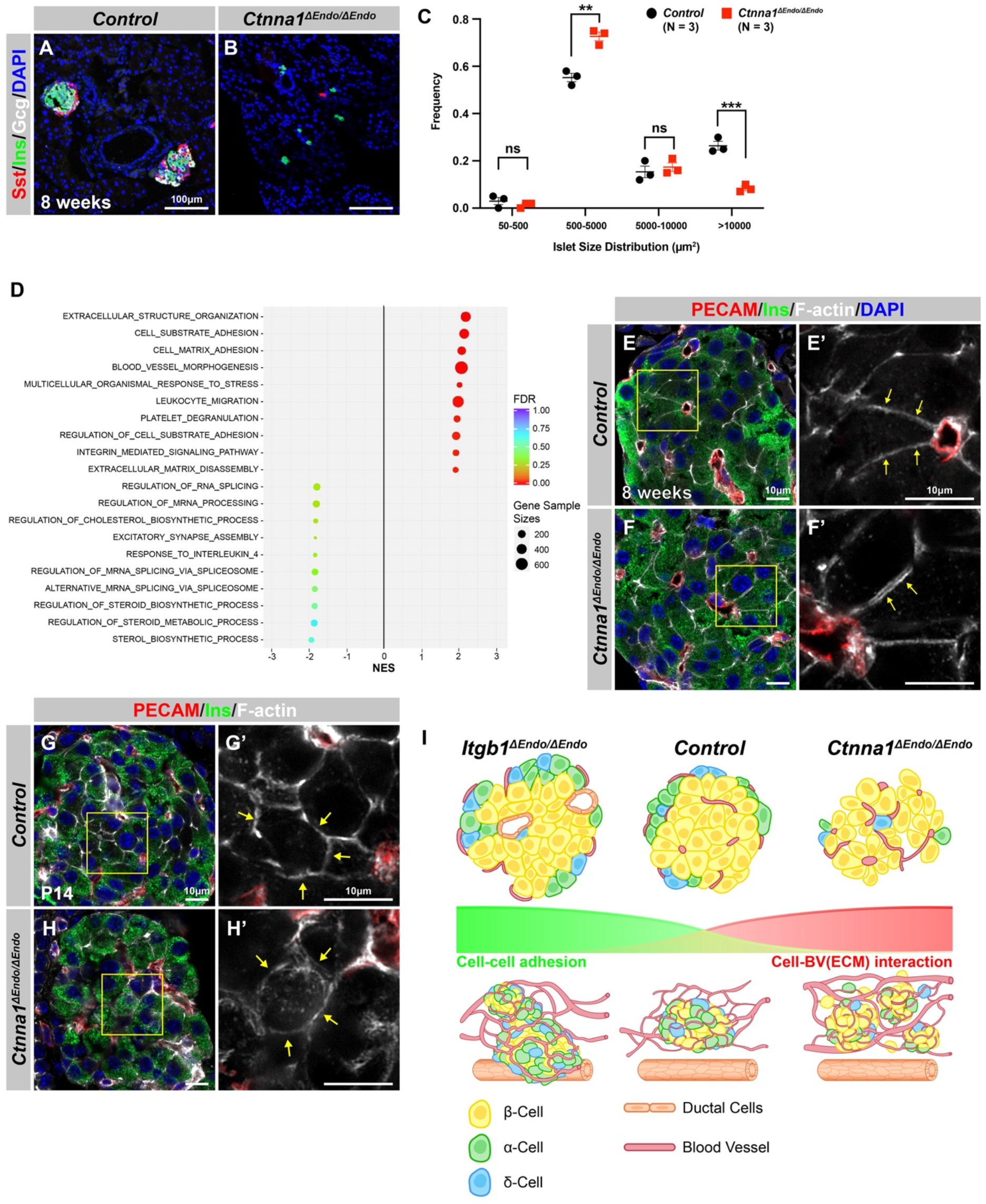
*Ctnna1*-deleted pancreatic islet endocrine cells display cytoskeleton anomalies. Related to Figure 3. (A-B) Immunofluorescence staining for Sst, Ins, and DAPI on pancreas sections from 8 weeks old mice. In *Ctnna1^ΔEndo/ΔEndo^*mice, the islet clusters appear much smaller compared to the control mice. (C) Quantification of islet size distribution based on ChrA staining for 8 weeks old control and *Ctnna1^ΔEndo/ΔEndo^* pancreas sections. The islet size quantification analysis excluded endocrine cell clusters with fewer than five cells. (D) Gene-set enrichment analysis of the differentially expressed genes, indicating top enriched pathways for control versus *Ctnna1^ΔEndo/ΔEndo^* islets at 8 weeks old. Gene sample sizes and FDRs are indicated by the size and color of the dots. (E-F) 3D Airyscan images of immunofluorescence staining for PECAM, Ins, and F-actin on 8 weeks old control and *Ctnna1^ΔEndo/ΔEndo^* pancreas sections. Fields demarcated by yellow boxes in E,F are shown at higher magnification in E’,F’. The images showed the separation of F-actin between Ins-expressing endocrine cells. This separation is highlighted by the yellow arrows in F’. (G-H) Airyscan images of immunofluorescence staining for PECAM, Ins, and F-actin in P14 control and *Ctnna1^ΔEndo/ΔEndo^*pancreas sections. In control islets (G’), the F-actin assemblies appear condensed, as indicated by the yellow arrows. However, in *Ctnna1^ΔEndo/ΔEndo^*islets (H’), the same F-actin assemblies appear to be diffused within the insulin-expressing cells. (I) Graphical summary showing cell-cell adhesion and cell-blood vessel (BV) interaction differences observed in *Itgb1^ΔEndo/ΔEndo^*and *Ctnna1^ΔEndo/ΔEndo^* islets. PECAM, Platelet endothelial cell adhesion molecule-1; Ins, Insulin; P14, postnatal day 14; BV, blood vessels Data are shown as mean ± SEM. ** p < 0.01 and *** p < 0.001.

**Figure S6.**
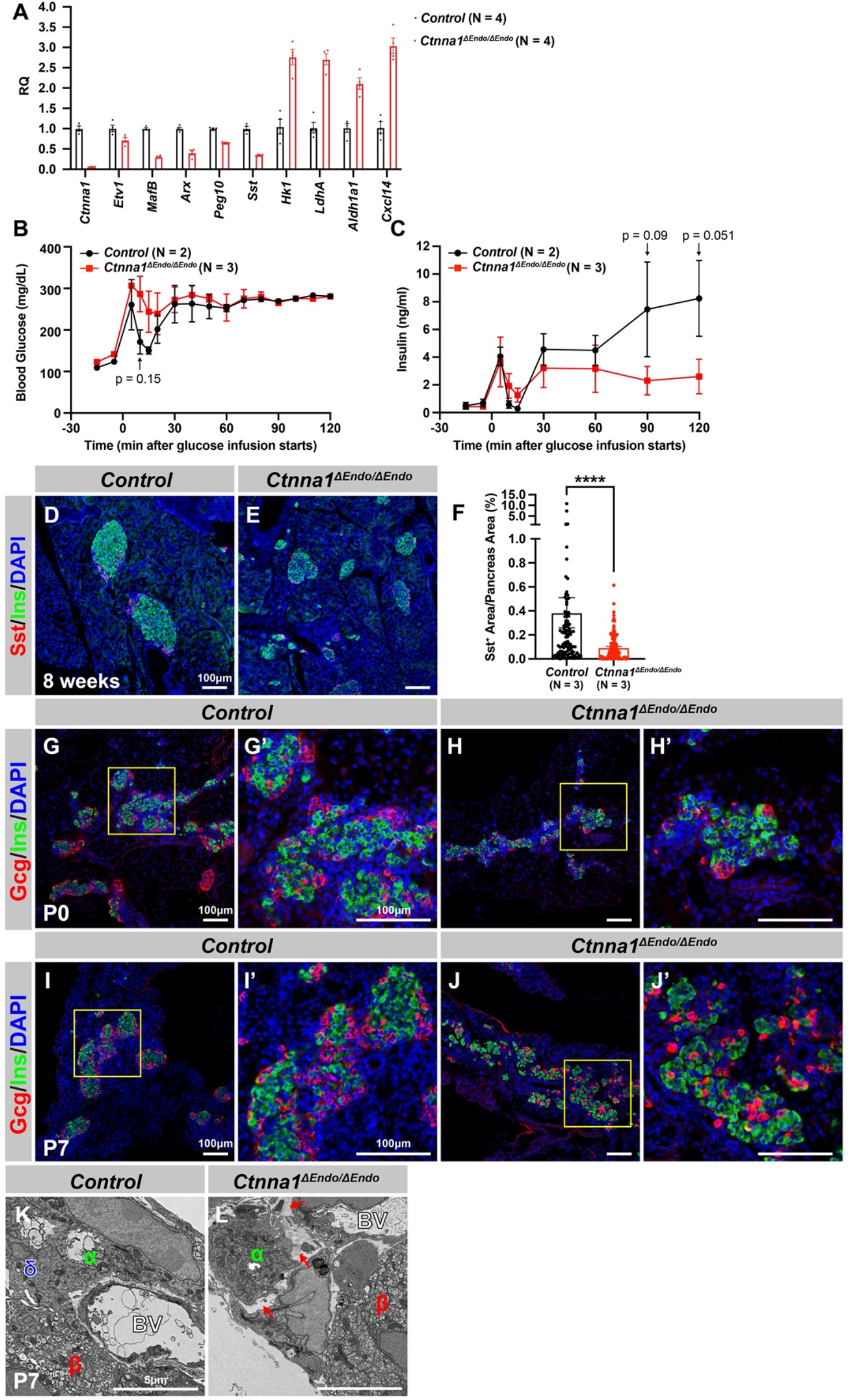
Ctnna1 is required for proper islet functionality and normal islet aggregation. Related to Figure 4. (A) Quantification of selected differentially expressed genes involved in islet functional maturation using real-time qPCR analysis in *Ctnna1^ΔEndo/ΔEndo^*islets and compared to controls. The relative quantification of gene expression levels was determined using the 2^-ΔΔCt^ method and is reported as a RQ value relative to the control group. The genes selected for validation were chosen based on their known roles in islet function and maturation. (B-C) Whole-blood glucose (B; mg/dL) and Insulin secretion (C; ng/ml) during 120 min in euglycemic clamp experiments in 5-h-fasted, surgically catheterized control and *Ctnna1^ΔEndo/ΔEndo^*mice at 8 weeks of age. At 5-15 minutes after glucose infusion, *Ctnna1^ΔEndo/ΔEndo^*mice showed slightly elevated glucose levels compared to controls (P=0.15). The amount of insulin secretion in *Ctnna1^ΔEndo/ΔEndo^* mice decreased at 90 and 120 minutes after glucose infusion (P=0.09 and 0.051, respectively). These results suggest that *Ctnna1^ΔEndo/ΔEndo^* mice may have impaired glucose metabolism and insulin secretion. (D-E) Immunofluorescence staining for Sst (red), Ins (green), and DAPI (blue) in pancreas sections of 8 weeks old control and *Ctnna1^ΔEndo/ΔEndo^* mice. (F) Quantification of the Sst^+^ area relative to the total pancreas area shows a decrease in the number of Sst^+^ cells in *Ctnna1^ΔEndo/ΔEndo^* mice compared with their control. (G-J) Immunofluorescence staining for Gcg (red), Ins (green), and DAPI (blue) in pancreas sections of P0 (G and H) and P7 (I and J) control and *Ctnna1^ΔEndo/ΔEndo^*mice. The fields demarcated by yellow boxes in (G-J) are shown at higher magnification in (G’-J’). (K-L) Transmission electron microscope images of P7 control (K) and *Ctnna1^ΔEndo/ΔEndo^* (L) endocrine areas. Red arrows indicate detachment of an α-cell from an islet aggregate. RQ, Relative quantification; Sst, Somatostatin; Gcg, Glucagon; Ins, Insulin; DAPI, 4’,6-diamidino-2-phenylindole; P0, postnatal day 0; P7, postnatal day 7; BV, blood vessels Data are shown as mean ± SEM. * p < 0.05, **** p < 0.0001.

**Figure S7.**
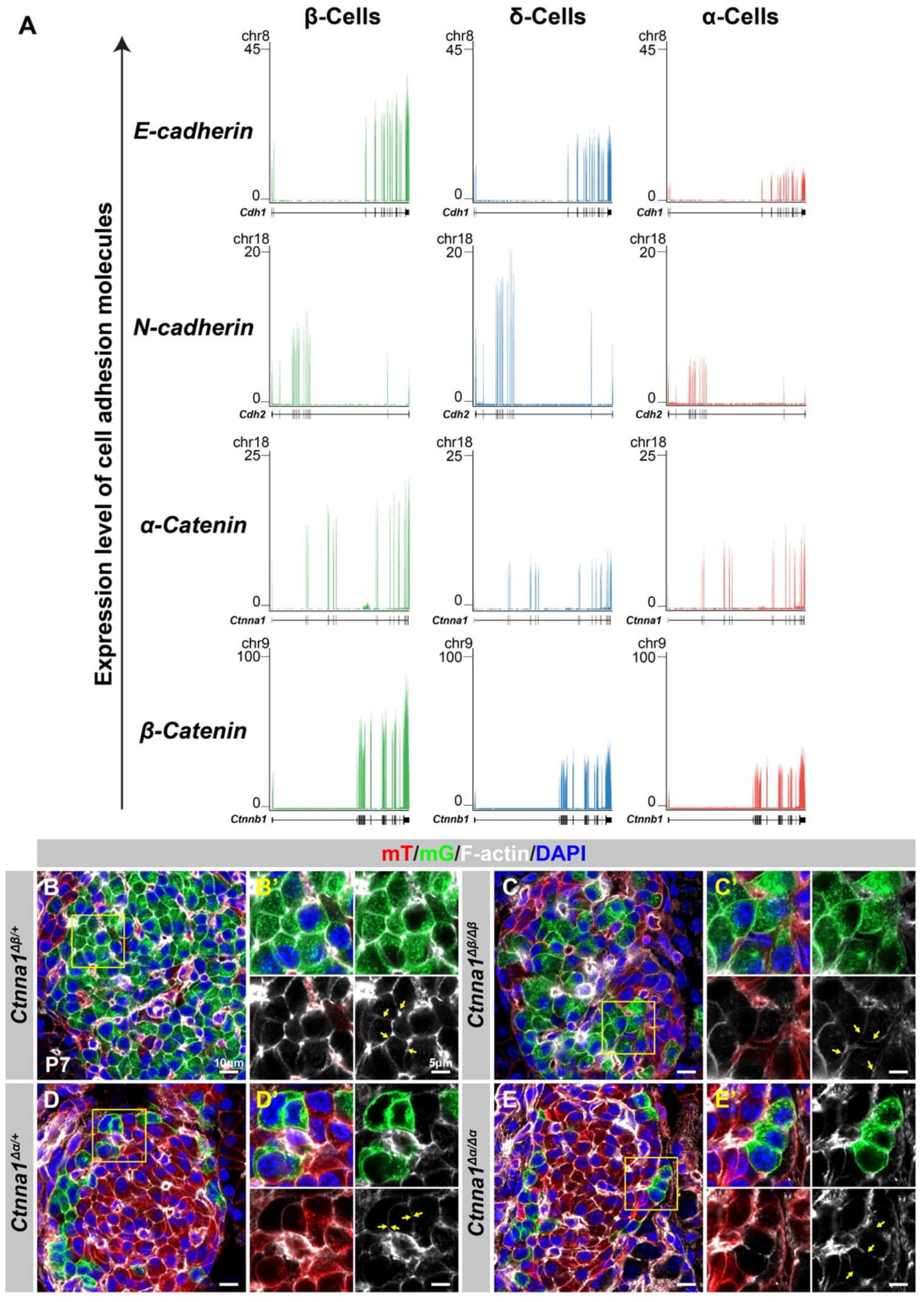
Differential cell-cell adhesion in endocrine cells. Related to Figure 5. (A) Genome viewer plots comparing expression levels of adhesion molecules in α-, *β*-, and δ- cell transcriptomes (DiGruccio et al., 2016). (B-E) Airyscan images of immunofluorescence staining of P7 mT/mG reporter mice stained for F-actin and DAPI in (B) heterozygous *Ctnna1^Δ*β*/+^*, (C) homozygous *Ctnna1^Δ*β*/Δ*β*^*, (D) heterozygous *Ctnna1^Δα/+^*, and (E) homozygous *Ctnna1^Δα/Δα^* mice. Fields demarcated by yellow boxes in B-E are shown at higher magnification with individual color channels in B’-E’ side panels. Yellow arrows indicate expression of F-actin in (B,C) and *β*- (D,E) α-cells. DAPI, 4’,6-diamidino-2-phenylindole; P7, postnatal day 7

**Figure S8.**
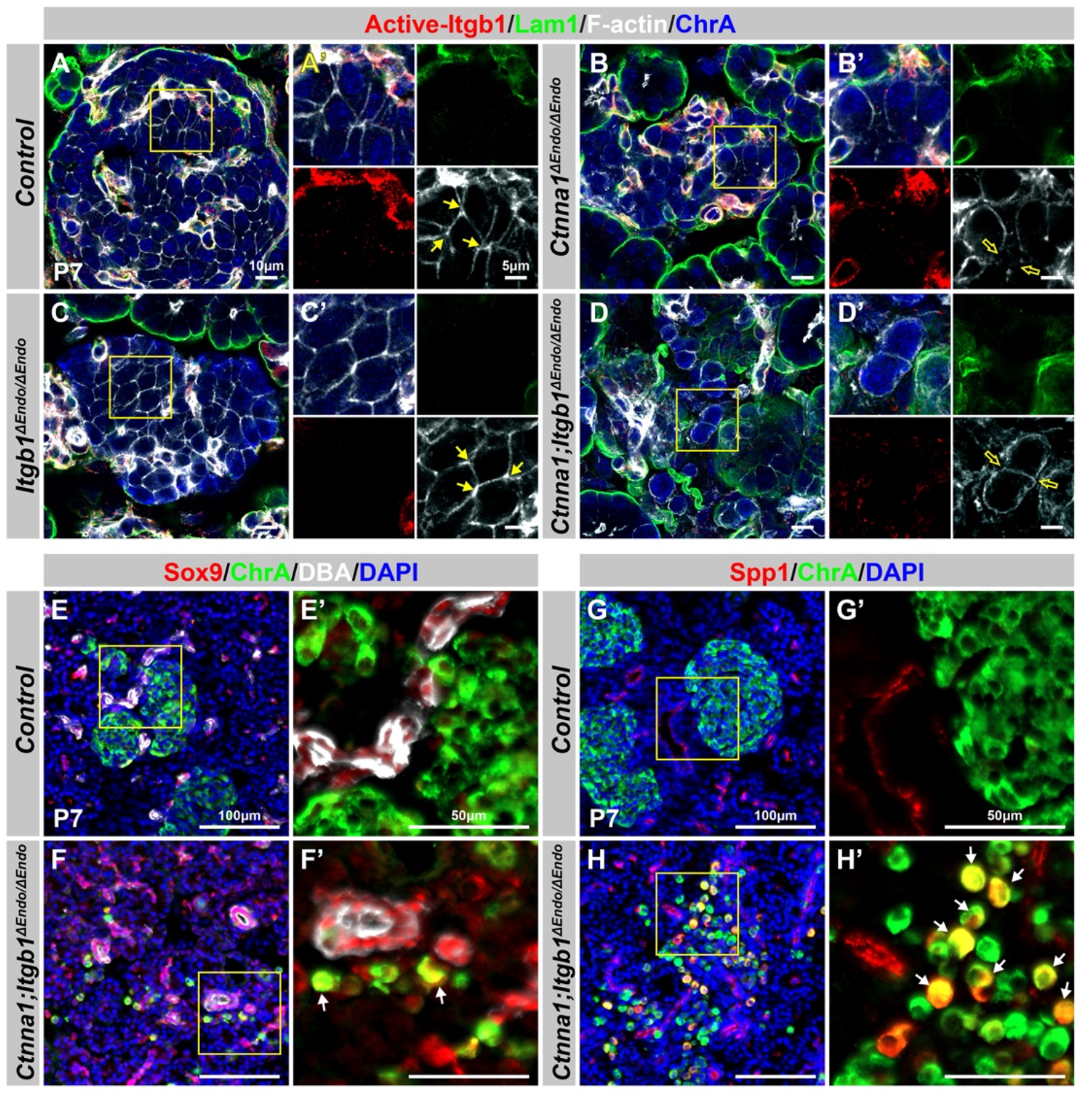
Aberrant endocrine cell clustering and upregulation of islet progenitor markers upon disruption of cell adhesion. Related to Figure 6. (A-D) Airyscan images showing immunofluorescence staining for active-Itgb1, Lam1, F-actin, and ChrA in pancreas sections from (A) control, (B) *Ctnna1^ΔEndo/ΔEndo^*, (C) *Itgb1^ΔEndo/ΔEndo^*, and (D) *Ctnna1; Itgb1^ΔEndo/ΔEndo^* mice at P7. The yellow boxes in A-D indicate the fields shown at higher magnification in A’-D’ side panels. The yellow arrows point to F-actin expression in endocrine cells. The *Ctnna1^ΔEndo/ΔEndo^*(B’) and *Ctnna1; Itgb1^ΔEndo/ΔEndo^* (D’) islet cells exhibit a round cell shape and reduced F-actin-enriched junctional structures. (E-H) Immunofluorescence staining for endocrine cell marker ChrA (green), ductal marker DBA (white in E, F), and progenitor markers Sox9 (red in E, F) and Spp1 (red in G, H) on pancreas sections from control (E, G) and *Ctnna1; Itgb1^ΔEndo/ΔEndo^*(F, H) mice at P7. The yellow boxes in (E-H) indicate the fields shown at higher magnification in (E’-H’) side panels. Compared to controls, in which ChrA^+^ endocrine cells do not express progenitor markers Sox9 and Spp1 (E’ and G’), a subpopulation of ChrA^+^ endocrine cells in the islets of *Ctnna1; Itgb1^ΔEndo/ΔEndo^* mice exhibit upregulated expression of Sox9 and Spp1 (yellow cells indicated by arrows in F’ and H’). Lam1, Laminin1; ChrA; Chromogranin A; P7, postnatal day 7; DBA, *Dolichos biflorus* agglutinin

## EXPERIMENTAL MODEL AND SUBJECTS DETAILS

### Mouse Strains

All animal experiments described herein were approved by the City of Hope Institutional Animal Care and Use Committees. Mice carrying *Ngn3-Cre* (Schonhoff et al., 2004), *R26^mT/mG^* (Muzumdar et al., 2007), *Gcg-iCre* (Shiota et al., 2017), *Itgb1^flox^* (Raghavan et al., 2000), *Ctnna1^flox^* (Vasioukhin et al., 2001), *Ins-Cre* (Thorens et al., 2015), *R26^mT/mG^* (Muzumdar et al., 2007), *R26^EYFP^*(Srinivas et al., 2001), and *Ucn3-cre* (Harris et al., 2014) alleles have been previously described. Additional information for the mouse lines used in this study is listed in Key Resources Table.

## METHOD DETAILS

### Whole Mount Immunofluorescence (WMIF)

A standard whole mount staining protocol was utilized to analyze pancreas three-dimensional (3D) morphology, as previously described (Maldonado et al., 2020). Briefly, pancreas tissue was dissected and fixed overnight in 4% paraformaldehyde (PFA). The tissue was dehydrated with serial methanol washes, bleached with a Dent’s Bleach solution, and rehydrated using an inverse series of methanol washes. Samples were blocked followed by incubation with primary antibodies for Platelet endothelial cell adhesion molecule-1 (PECAM), Chromogranin A (ChrA), and Secreted Phosphoprotein 1 (Spp1). Samples were then washed in a 0.15% triton-X-PBS solution and stained with secondary antibodies overnight. Following additional washes with 0.15% triton-X-PBS, samples were fixed with 4% PFA, and dehydrated with serial ethanol washes. Finally, samples were immersed in a BABB clearing solution (two parts benzyl benzoate and one part benzyl alcohol) and imaged using a Zeiss LSM880 confocal microscope.

### Confocal Microscopy

All images were collected with a Zeiss LSM880 confocal microscope equipped with a 10X 0.45NA plan apochromat objective at 2X magnification, as standard for AiryscanFast collection. All tiled Z-stacks were acquired in AiryscanFast mode, the acquisition method set to Z-stack (per channel), and with a 2.0 micron step size. Scans were conducted using 405nm, 488nm, 561nm, and 633nm laser lines, and emissions were collected using standard Airyscan filters and the Airyscan detector, in Zen Black 2.3 SP1 (Carl Zeiss). Final voxel sizes were 0.22x0.22x1.98 microns, and no digital gain or detector offset was used. Detector gains and laser intensities were consistent across all samples.

### Image Segmentation/Analysis

Data sets were first Airyscan processed in Zen Black using the “Auto” setting using a 2D sample image as a base for Batch mode. After processing, the files were converted to IMS format and analyzed in Imaris 9.5 (Bitplane). Analysis consisted of manually selecting thresholds that could be consistently applied between samples, and best fit the biology. Due to variations in the depth of the sample, amount of tissue the light had to pass through, and orientation, thresholds were chosen which most accurately represented the objects being analyzed. The ducts were first segmented using a “Surface” with a 2.00-micron surface grain size, and a fixed threshold. We limited the analysis to areas where ducts were present by segmenting the duct channel a second time (again as a Surface) with a grain size of 10 microns and a much lower threshold, generating a “bubble” area of interest around the ducts. Within this volume, endocrine positive staining was then segmented as another surface and its distance from the ducts quantified using the median value of the distance transform from the surface of the original ducts. Any endocrine cells contained entirely or mostly within the ducts were considered to have a distance of zero from the ducts.

### Immunofluorescence (IF)

Pancreas tissues were collected from littermate control and mutant mice, ensuring consistency in the sample source. Immunofluorescence staining was performed as previously described (Shih et al., 2007). Briefly, pancreata were fixed in 4% paraformaldehyde in PBS, equilibrated in 30% sucrose in PBS, and cryoembedded in Tissue-Tek OCT. To ensure the reliability and accuracy of the findings, we used littermate control and mutant mice, collecting and processing their pancreas samples concurrently. This included fixing and embedding the samples into the same cryo-block, sectioning, and preparing them for immunofluorescence staining on the same slide. This process ensured identical staining conditions for all samples. Samples were then sectioned at 10 μm intervals from the same cryo-block, facilitating identical staining conditions for all samples. For immunofluorescence staining, sections were blocked for 1 hour before the addition of primary antibodies diluted in the same buffer. Following washes, secondary antibodies, diluted in blocking buffer, were applied to sections and stained for 2 hours at room temperature. The respective dilutions of primary and secondary antibodies are noted in the Key Resources Table.

Images were taken on a Zeiss Axio-Observer-Z1 microscope with Apotome, and figures were created in Adobe Photoshop/Illustrator CC2020. Morphometric analyses were conducted using Image-Pro Premier v.9.2 and Qupath software, available at https://qupath.github.io. High resolution image acquisition was achieved with a Zeiss LSM900 confocal microscope with Airyscan. For images intended for direct comparison, the exposure time was kept constant, and changes in digital gain were avoided to prevent distortion of results. Upon image acquisition, linear adjustment processing was applied uniformly across all comparable samples. These adjustments were made using a linear display with Min/Max rescaling. This adjustment approach was to carefully balance avoiding oversaturation while preserving the accuracy of marker expressions. This approach ensures that the full data range is properly utilized, optimizing the full range of possible intensity values. Specifically, the darkest pixel in the image is adjusted to black (minimum value), the brightest pixel to white (maximum value), and all other pixels are rescaled based on these new extremes.

### α-cell Localization Analysis

The islets were imaged from three serial sections per mouse, taken at 150 μm (P7) or 200 μm (8 weeks) intervals. Manual cell counting was performed using QuPath (version 0.2.3). Islets with a minimum of 40 endocrine cells and at least 10 α-cells were included in the analysis. At the postnatal stage, α-cell localization within the islet was determined according to previously described criteria (Adams et al., 2018). Peripheral α-cells were defined as those within the first two layers, while α-cells in any layers beyond the first two were considered to be in the core.

### Intra Peritoneal Glucose Tolerance Test (IPGTT)

6 to 16 weeks old or 6 to 12 months old mice were fasted for 6-16 hours. Basal blood glucose was measured, and mice were injected intraperitoneally (i.p.) with 2 mg/g body weight dextrose solution. Blood glucose was measured at 15, 30, 60, 90 and 120 minutes after glucose challenge. The experiments were performed by the Comprehensive Metabolic Phenotyping (CMP) Core at City of Hope.

### Insulin secretion *in vivo*

The *β*-cell function was assessed by quantifying insulin secretion *in vivo* with hyperglycemic clamps. Surgical procedures have been described in detail previously(Ayala et al., 2011; Berglund et al., 2008). Briefly, mice were anesthetized with isoflurane, and the carotid artery and jugular vein were catheterized. Free catheter ends were tunneled under the skin to the back of the neck and externalized with a MASA^TM^ (Mouse Antenna for Sampling Access), which permits arterial sampling from an indwelling catheter in conscious mice without handling stress. Mice were individually housed and allowed to recover for 4 days during which body weight returned to within ∼10% of presurgical body weight.

For the hyperglycemic clamps, mice were fasted for 5 h and connected to a swivel. Saline-washed erythrocytes were infused (5–6 μl/min) during the experimental period to prevent a >5% fall in hematocrit. Blood samples were collected from the arterial catheter in tubes containing EDTA and centrifuged, and plasma was stored at −20°C until analyzed. At t = 0 min, mice received a priming glucose infusion rate (GIR, 100 mg/kg/min) for 2 min to stimulate first-phase insulin secretion. The GIR was then adjusted to achieve and maintain blood glucose at ∼15.0 mmol/l. Blood glucose (5 μl) was measured at t = −15, −5, 5, 10, 15, and 20 min and then every 10 min until t = 120 min with an Alpha Trak glucometer (Abbott Laboratories, Abbott Park, IL, USA). Larger blood samples (100 μl) to measure plasma insulin were taken at various timepoints.

### Isolation of Mouse Pancreatic Islets

Pancreatic islets were isolated as previously described (Taylor et al., 2013). Briefly, mouse pancreatic islets were isolated using 2.5 mg/ml collagenase (Sigma Aldrich) as the digestion method. Islets were collected under a stereomicroscope and allowed to recover overnight in PIM(R)® 32A (Prodo Labs) medium with 5% HSA, Gentamicin-AmphotericinB, and 5.8 mmol/L glucose.

### RNA-Sequencing (RNA-seq)

Total RNA was isolated using the RNeasy Micro Kit (Qiagen) with 10 to 30 isolated islets and processed per manufacturer’s instructions. cDNA libraries were prepared using the KAPA RNA HyperPrep Kit with RiboErase (HMR) according to the manufacturer’s instructions. Libraries were subjected to high-throughput sequencing on Illumina HiSeq 2500 (50 bp, single-ended) and subsequent data analysis was completed by the Integrative Genomics Core at City of Hope. For sequence alignment and gene counts, RNA-Seq reads were trimmed to remove sequencing adapters using Trimmomatic (Bolger et al., 2014) and polyA tails using FASTP (Chen et al., 2018). The processed reads were mapped back to the mouse genome (mm10) using STAR software (v. 020201) (Dobin et al., 2013). The featureCount software was applied to generate the count matrix, with default parameters (Liao et al., 2019). Between two sets of samples (Control vs Mutant), differential expression analysis was conducted, in R, by adjusting read counts to normalized expression values using TMM normalization method and by correcting batch effects using a design matrix (Robinson et al., 2010). Genes with an FDR-adjusted p-value less than 0.05 and with a fold change (FC) greater than 2 or less than 0.5 were considered as significant up- and down-regulated genes, respectively. Pathway analysis was conducted using DAVID (v. 6.8) (Huang da et al., 2009a, b) and GSEAPreranked algorithm using GSEA Desktop program in Java (Subramanian et al., 2005). The former requires a list of up- and down-regulated genes, while the latter a ranked list of whole genes according to their log2 fold change and p-values. Raw data and processed data files were available at the GEO database: (GSE153187, https://www.ncbi.nlm.nih.gov/geo/query/acc.cgi?acc=GSE153187). (GSE190788, https://www.ncbi.nlm.nih.gov/geo/query/acc.cgi?acc=GSE190788).

### Quantitative real-time PCR (qPCR)

RNA extraction and purification were performed using the RNeasy Micro Kit (QIAGEN) in accordance with the manufacturer’s protocol. The quality and quantity of the total RNA were assessed using a Bioanalyzer 2100 (Agilent Technologies). Samples with an RNA integrity number (RIN) of at least 6 were used for cDNA synthesis with the SuperScript^TM^ First-Strand Synthesis SuperMix for qRT-PCR (Invitrogen). Real-time PCR was conducted using PowerUp^TM^ SYBR^TM^ Green Master mix (Applied Biosystems) in a QuantStudio 6 Flex machine (Applied Biosystems). Each reaction contained 1 ng cDNA (total RNA equivalent). Relative mRNA expression levels were calculated by normalizing target genes to *Cyclophilin A* (*Ppia*) as an internal control. All primers were designed using the CDS of the target genes by the PrimerBank online tool (https://pga.mgh.harvard.edu/primerbank/).

### Aggregation Assay

Neonatal *Ngn3-Cre; R26mT/mG; Itgb1^+/f^* (control) or *Ngn3-Cre; R26mT/mG; Itgb1^f/f^* (mutant) pancreata were dissociated into a single cell suspension using TrypLE (Thermo Fisher). Enzymatic activity was neutralized using FACS buffer (10% fetal bovine serum (FBS) and 2mM EDTA in CMRL media). Cells were passed through a 40 µm cell strainer and DAPI was used to stain dead cells prior to FACS. Prior to cell seeding, 96-well round bottom plates were coated with either bovine serum albumin (BSA; no ECM) or Matrigel and fibronectin (+ECM). 3000 cells per well were added into no ECM or +ECM wells and cultured for 48 hours.

### *Ex Vivo* Pancreas Explants

Pancreas explants are described previously (Shih and Sander, 2014). After 24 hours of culture, the explants were placed on the microscope stage in 37 °C culture chambers with a controlled atmosphere of humidified 5% CO_2_. Time-lapse imaging was performed using a Zeiss LSM880 inverted confocal microscope with a C-Apochromat 20×/0.75 objective lens. Explants were optically sectioned every 2.5 µm in a 512 × 512 format with up to 80 µm Z-stacks every 10 min for 72 hours. Images were acquired with Zen software and then reconstructed in 3D with Imaris 9.5 (Bitplane) software. The contrast was adjusted and selected optical planes or *z*-projections of sequential optical sections were used to assemble time-lapse movies.

### Electron Microscopy Sample Preparation

Sample blocks were prepared following the NCMIR protocol (T.J. Deerinck, 2010). Briefly, small pieces (1∼2 mm^3^) of mouse pancreatic tissues were dissected and immediately fixed in 0.2M cacodylate buffer pH7.4 containing 2.5% glutaraldehyde and 2% paraformaldehyde with 3mM calcium chloride. After primary aldehyde fixation the samples were rinsed in 0.15M sodium cacodylate pH7.4 containing 2mM calcium chloride and post-fixed in 1.5% potassium ferrocyanide-reduced 2% osmium tetroxide in 0.15M cacodylate buffer for 1 hour. Tissue was then rinsed in distilled water and treated with 0.1% aqueous thiocarbohydrazide for 20 min. After further rinsing in distilled water the tissue was again treated with 2% osmium tetroxide for 30 min, rinsed in distilled water, dehydrated in an ethanol series, and infiltrated with Durcupan ACM resin. Ultra-thin sections (70 nm thick) were acquired by ultramicrotomy and examined on a FEI Tecnai 12 transmission electron microscope equipped with a Gatan OneView CMOS camera.

### Statistics

The statistical analysis was performed using GraphPad Prism. To compare groups, one-way analysis of variance (ANOVA) and a two-tailed student’s t-test were utilized, unless otherwise specified. Each group was evaluated using a minimum of three independent measurements. Unless stated otherwise, all data are presented as mean ± standard error of the mean (SEM).

## Notes

### Competing Interest Statement

The authors have declared no competing interest.

### Summary of Updates

Additional authors; major revision on the manuscript and figures

## REFERENCES

Adams, M.T., and Blum, B. (2022). Determinants and dynamics of pancreatic islet architecture. Islets 14, 82–100.

Adams, M.T., Dwulet, J.M., Briggs, J.K., Reissaus, C.A., Jin, E., Szulczewski, J.M., Lyman, M.R., Sdao, S.M., Kravets, V., Nimkulrat, S.D., et al. (2021). Reduced synchroneity of intra-islet Ca(2+) oscillations in vivo in Robo-deficient beta cells. Elife 10.

Adams, M.T., Gilbert, J.M., Hinojosa Paiz, J., Bowman, F.M., and Blum, B. (2018). Endocrine cell type sorting and mature architecture in the islets of Langerhans require expression of Roundabout receptors in beta cells. Sci Rep 8, 10876.

Alto, L.T., and Terman, J.R. (2017). Semaphorins and their Signaling Mechanisms. Methods Mol Biol 1493, 1–25.

Arrojo e Drigo, R., Ali, Y., Diez, J., Srinivasan, D.K., Berggren, P.O., and Boehm, B.O. (2015). New insights into the architecture of the islet of Langerhans: a focused cross-species assessment. Diabetologia 58, 2218–2228.

Ayala, J.E., Bracy, D.P., Malabanan, C., James, F.D., Ansari, T., Fueger, P.T., McGuinness, O.P., and Wasserman, D.H. (2011). Hyperinsulinemic-euglycemic clamps in conscious, unrestrained mice. J Vis Exp.

Balboa, D., Barsby, T., Lithovius, V., Saarimaki-Vire, J., Omar-Hmeadi, M., Dyachok, O., Montaser, H., Lund, P.E., Yang, M., Ibrahim, H., et al. (2022). Functional, metabolic and transcriptional maturation of human pancreatic islets derived from stem cells. Nat Biotechnol 40, 1042–1055.

Barros, C.S., Franco, S.J., and Muller, U. (2011). Extracellular matrix: functions in the nervous system. Cold Spring Harb Perspect Biol 3, a005108.

Bastidas-Ponce, A., Tritschler, S., Dony, L., Scheibner, K., Tarquis-Medina, M., Salinno, C., Schirge, S., Burtscher, I., Bottcher, A., Theis, F.J., et al. (2019). Comprehensive single cell mRNA profiling reveals a detailed roadmap for pancreatic endocrinogenesis. Development 146.

Bechard, M.E., Bankaitis, E.D., Hipkens, S.B., Ustione, A., Piston, D.W., Yang, Y.P., Magnuson, M.A., and Wright, C.V. (2016). Precommitment low-level Neurog3 expression defines a long-lived mitotic endocrine-biased progenitor pool that drives production of endocrine-committed cells. Genes Dev 30, 1852–1865.

Berglund, E.D., Li, C.Y., Poffenberger, G., Ayala, J.E., Fueger, P.T., Willis, S.E., Jewell, M.M., Powers, A.C., and Wasserman, D.H. (2008). Glucose metabolism in vivo in four commonly used inbred mouse strains. Diabetes 57, 1790–1799.

Blum, B., Roose, A.N., Barrandon, O., Maehr, R., Arvanites, A.C., Davidow, L.S., Davis, J.C., Peterson, Q.P., Rubin, L.L., and Melton, D.A. (2014). Reversal of beta cell de-differentiation by a small molecule inhibitor of the TGFbeta pathway. Elife 3, e02809.

Bolger, A.M., Lohse, M., and Usadel, B. (2014). Trimmomatic: a flexible trimmer for Illumina sequence data. Bioinformatics 30, 2114–2120.

Campbell, J.E., and Newgard, C.B. (2021). Mechanisms controlling pancreatic islet cell function in insulin secretion. Nat Rev Mol Cell Biol 22, 142–158.

Carlson, T.R., Hu, H., Braren, R., Kim, Y.H., and Wang, R.A. (2008). Cell-autonomous requirement for beta1 integrin in endothelial cell adhesion, migration and survival during angiogenesis in mice. Development 135, 2193–2202.

Chen, S., Zhou, Y., Chen, Y., and Gu, J. (2018). fastp: an ultra-fast all-in-one FASTQ preprocessor. Bioinformatics 34, i884–i890.

Cirulli, V., Beattie, G.M., Klier, G., Ellisman, M., Ricordi, C., Quaranta, V., Frasier, F., Ishii, J.K., Hayek, A., and Salomon, D.R. (2000). Expression and function of alpha(v)beta(3) and alpha(v)beta(5) integrins in the developing pancreas: roles in the adhesion and migration of putative endocrine progenitor cells. J Cell Biol 150, 1445–1460.

Cottle, L., Gan, W.J., Gilroy, I., Samra, J.S., Gill, A.J., Loudovaris, T., Thomas, H.E., Hawthorne, W.J., Kebede, M.A., and Thorn, P. (2021). Structural and functional polarisation of human pancreatic beta cells in islets from organ donors with and without type 2 diabetes. Diabetologia 64, 618–629.

Dahl, U., Sjodin, A., and Semb, H. (1996). Cadherins regulate aggregation of pancreatic beta-cells in vivo. Development 122, 2895–2902.

Diaferia, G.R., Jimenez-Caliani, A.J., Ranjitkar, P., Yang, W., Hardiman, G., Rhodes, C.J., Crisa, L., and Cirulli, V. (2013). beta1 integrin is a crucial regulator of pancreatic beta-cell expansion. Development 140, 3360–3372.

DiGruccio, M.R., Mawla, A.M., Donaldson, C.J., Noguchi, G.M., Vaughan, J., Cowing-Zitron, C., van der Meulen, T., and Huising, M.O. (2016). Comprehensive alpha, beta and delta cell transcriptomes reveal that ghrelin selectively activates delta cells and promotes somatostatin release from pancreatic islets. Mol Metab 5, 449–458.

Dobin, A., Davis, C.A., Schlesinger, F., Drenkow, J., Zaleski, C., Jha, S., Batut, P., Chaisson, M., and Gingeras, T.R. (2013). STAR: ultrafast universal RNA-seq aligner. Bioinformatics 29, 15–21.

Dybala, M.P., Butterfield, J.K., Hendren-Santiago, B.K., and Hara, M. (2020). Pancreatic Islets and Gestalt Principles. Diabetes 69, 1864-1874.

Elghazi, L., Gould, A.P., Weiss, A.J., Barker, D.J., Callaghan, J., Opland, D., Myers, M., Cras-Meneur, C., and Bernal-Mizrachi, E. (2012). Importance of beta-Catenin in glucose and energy homeostasis. Sci Rep 2, 693.

Esni, F., Taljedal, I.B., Perl, A.K., Cremer, H., Christofori, G., and Semb, H. (1999). Neural cell adhesion molecule (N-CAM) is required for cell type segregation and normal ultrastructure in pancreatic islets. J Cell Biol 144, 325–337.

Fagotto, F. (2014). The cellular basis of tissue separation. Development 141, 3303–3318.

Foty, R.A., and Steinberg, M.S. (2005). The differential adhesion hypothesis: a direct evaluation. Dev Biol 278, 255–263.

Gan, W.J., Do, O.H., Cottle, L., Ma, W., Kosobrodova, E., Cooper-White, J., Bilek, M., and Thorn, P. (2018). Local Integrin Activation in Pancreatic beta Cells Targets Insulin Secretion to the Vasculature. Cell Rep 24, 2819–2826 e2813.

Gilbert, J.M., Adams, M.T., Sharon, N., Jayaraaman, H., and Blum, B. (2020). Morphogenesis of the islets of Langerhans is guided by extra-endocrine Slit2/3 signals. Mol Cell Biol.

Gonschior, H., Haucke, V., and Lehmann, M. (2020). Super-Resolution Imaging of Tight and Adherens Junctions: Challenges and Open Questions. Int J Mol Sci 21.

Gouzi, M., Kim, Y.H., Katsumoto, K., Johansson, K., and Grapin-Botton, A. (2011). Neurogenin3 initiates stepwise delamination of differentiating endocrine cells during pancreas development. Dev Dyn 240, 589–604.

Gu, G., Dubauskaite, J., and Melton, D.A. (2002). Direct evidence for the pancreatic lineage: NGN3+ cells are islet progenitors and are distinct from duct progenitors. Development 129, 2447–2457.

Halban, P.A., Powers, S.L., George, K.L., and Bonner-Weir, S. (1987). Spontaneous reassociation of dispersed adult rat pancreatic islet cells into aggregates with three-dimensional architecture typical of native islets. Diabetes 36, 783–790.

Hannigan, G., Troussard, A.A., and Dedhar, S. (2005). Integrin-linked kinase: a cancer therapeutic target unique among its ILK. Nat Rev Cancer 5, 51–63.

Harris, J.A., Hirokawa, K.E., Sorensen, S.A., Gu, H., Mills, M., Ng, L.L., Bohn, P., Mortrud, M., Ouellette, B., Kidney, J., et al. (2014). Anatomical characterization of Cre driver mice for neural circuit mapping and manipulation. Front Neural Circuits 8, 76.

Heinis, M., Simon, M.T., Ilc, K., Mazure, N.M., Pouyssegur, J., Scharfmann, R., and Duvillie, B. (2010). Oxygen tension regulates pancreatic beta-cell differentiation through hypoxia-inducible factor 1alpha. Diabetes 59, 662–669.

Huang da, W., Sherman, B.T., and Lempicki, R.A. (2009a). Bioinformatics enrichment tools: paths toward the comprehensive functional analysis of large gene lists. Nucleic Acids Res 37, 1–13.

Huang da, W., Sherman, B.T., and Lempicki, R.A. (2009b). Systematic and integrative analysis of large gene lists using DAVID bioinformatics resources. Nat Protoc 4, 44–57.

Jamora, C., and Fuchs, E. (2002). Intercellular adhesion, signalling and the cytoskeleton. Nat Cell Biol 4, E101–108.

Jiang, L., Sun, J., and Huang, D. (2022). Role of Slit/Robo Signaling pathway in Bone Metabolism. Int J Biol Sci 18, 1303–1312.

Jimenez-Caliani, A.J., Pillich, R., Yang, W., Diaferia, G.R., Meda, P., Crisa, L., and Cirulli, V. (2017). alphaE-Catenin Is a Positive Regulator of Pancreatic Islet Cell Lineage Differentiation. Cell Rep 20, 1295–1306.

Jo, J., Kilimnik, G., Kim, A., Guo, C., Periwal, V., and Hara, M. (2011). Formation of pancreatic islets involves coordinated expansion of small islets and fission of large interconnected islet-like structures. Biophys J 101, 565–574.

Kim, A., Miller, K., Jo, J., Kilimnik, G., Wojcik, P., and Hara, M. (2009). Islet architecture: A comparative study. Islets 1, 129–136.

Kragl, M., Schubert, R., Karsjens, H., Otter, S., Bartosinska, B., Jeruschke, K., Weiss, J., Chen, C., Alsteens, D., Kuss, O., et al. (2016). The biomechanical properties of an epithelial tissue determine the location of its vasculature. Nat Commun 7, 13560.

Krishnamurthy, M., Li, J., Fellows, G.F., Rosenberg, L., Goodyer, C.G., and Wang, R. (2011). Integrin alpha3, but not beta1, regulates islet cell survival and function via PI3K/Akt signaling pathways. Endocrinology 152, 424–435.

Kwon, H.R., Nelson, D.A., DeSantis, K.A., Morrissey, J.M., and Larsen, M. (2017). Endothelial cell regulation of salivary gland epithelial patterning. Development 144, 211–220.

Lammert, E., Gu, G., McLaughlin, M., Brown, D., Brekken, R., Murtaugh, L.C., Gerber, H.P., Ferrara, N., and Melton, D.A. (2003). Role of VEGF-A in vascularization of pancreatic islets. Curr Biol 13, 1070–1074.

Larsen, H.L., and Grapin-Botton, A. (2017). The molecular and morphogenetic basis of pancreas organogenesis. Semin Cell Dev Biol 66, 51–68.

Lemaire, K., Thorrez, L., and Schuit, F. (2016). Disallowed and Allowed Gene Expression: Two Faces of Mature Islet Beta Cells. Annu Rev Nutr 36, 45–71.

Li, J., Zhao, Z., Wang, J., Chen, G., Yang, J., and Luo, S. (2008). The role of extracellular matrix, integrins, and cytoskeleton in mechanotransduction of centrifugal loading. Mol Cell Biochem 309, 41–48.

Liao, Y., Smyth, G.K., and Shi, W. (2019). The R package Rsubread is easier, faster, cheaper and better for alignment and quantification of RNA sequencing reads. Nucleic Acids Res 47, e47.

Lien, W.H., Klezovitch, O., Fernandez, T.E., Delrow, J., and Vasioukhin, V. (2006). alphaE-catenin controls cerebral cortical size by regulating the hedgehog signaling pathway. Science 311, 1609-1612.

Maldonado, M., Serrill, J.D., and Shih, H.P. (2020). Painting the Pancreas in Three Dimensions: Whole-Mount Immunofluorescence Method. Methods Mol Biol 2155, 193–200.

Mohamed, T.H., Watanabe, H., Kaur, R., Belyea, B.C., Walker, P.D., Gomez, R.A., and Sequeira-Lopez, M.L.S. (2020). Renin-Expressing Cells Require beta1-Integrin for Survival and for Development and Maintenance of the Renal Vasculature. Hypertension 76, 458–467.

Mui, K.L., Chen, C.S., and Assoian, R.K. (2016). The mechanical regulation of integrin-cadherin crosstalk organizes cells, signaling and forces. J Cell Sci 129, 1093–1100.

Murtaugh, L.C., Law, A.C., Dor, Y., and Melton, D.A. (2005). Beta-catenin is essential for pancreatic acinar but not islet development. Development 132, 4663–4674.

Muzumdar, M.D., Tasic, B., Miyamichi, K., Li, L., and Luo, L. (2007). A global double-fluorescent Cre reporter mouse. Genesis 45, 593–605.

Nair, G.G., Liu, J.S., Russ, H.A., Tran, S., Saxton, M.S., Chen, R., Juang, C., Li, M.L., Nguyen, V.Q., Giacometti, S., et al. (2019). Recapitulating endocrine cell clustering in culture promotes maturation of human stem-cell-derived beta cells. Nat Cell Biol 21, 263–274.

Panzer, J.K., Cohrs, C.M., and Speier, S. (2020). Using Pancreas Tissue Slices for the Study of Islet Physiology. Methods Mol Biol 2128, 301–312.

Pauerstein, P.T., Tellez, K., Willmarth, K.B., Park, K.M., Hsueh, B., Efsun Arda, H., Gu, X., Aghajanian, H., Deisseroth, K., Epstein, J.A., et al. (2017). A radial axis defined by semaphorin-to-neuropilin signaling controls pancreatic islet morphogenesis. Development 144, 3744–3754.

Raghavan, S., Bauer, C., Mundschau, G., Li, Q., and Fuchs, E. (2000). Conditional ablation of beta1 integrin in skin. Severe defects in epidermal proliferation, basement membrane formation, and hair follicle invagination. J Cell Biol 150, 1149–1160.

Reinert, R.B., Cai, Q., Hong, J.Y., Plank, J.L., Aamodt, K., Prasad, N., Aramandla, R., Dai, C., Levy, S.E., Pozzi, A., et al. (2014). Vascular endothelial growth factor coordinates islet innervation via vascular scaffolding. Development 141, 1480–1491.

Robinson, M.D., McCarthy, D.J., and Smyth, G.K. (2010). edgeR: a Bioconductor package for differential expression analysis of digital gene expression data. Bioinformatics 26, 139–140.

Romer, A.I., and Sussel, L. (2015). Pancreatic islet cell development and regeneration. Curr Opin Endocrinol Diabetes Obes 22, 255–264.

Roscioni, S.S., Migliorini, A., Gegg, M., and Lickert, H. (2016). Impact of islet architecture on beta-cell heterogeneity, plasticity and function. Nat Rev Endocrinol 12, 695–709.

Rosenberg, L.C., Lafon, M.L., Pedersen, J.K., Yassin, H., Jensen, J.N., Serup, P., and Hecksher-Sorensen, J. (2010). The transcriptional activity of Neurog3 affects migration and differentiation of ectopic endocrine cells in chicken endoderm. Dev Dyn 239, 1950–1966.

Rulifson, I.C., Karnik, S.K., Heiser, P.W., ten Berge, D., Chen, H., Gu, X., Taketo, M.M., Nusse, R., Hebrok, M., and Kim, S.K. (2007). Wnt signaling regulates pancreatic beta cell proliferation. Proc Natl Acad Sci U S A 104, 6247–6252.

Sakai, T., Larsen, M., and Yamada, K.M. (2003). Fibronectin requirement in branching morphogenesis. Nature 423, 876–881.

Scheppke, L., Murphy, E.A., Zarpellon, A., Hofmann, J.J., Merkulova, A., Shields, D.J., Weis, S.M., Byzova, T.V., Ruggeri, Z.M., Iruela-Arispe, M.L., et al. (2012). Notch promotes vascular maturation by inducing integrin-mediated smooth muscle cell adhesion to the endothelial basement membrane. Blood 119, 2149–2158.

Schiesser, J.V., Loudovaris, T., Thomas, H.E., Elefanty, A.G., and Stanley, E.G. (2021). Integrin alphavbeta5 heterodimer is a specific marker of human pancreatic beta cells. Sci Rep 11, 8315.

Schonhoff, S.E., Giel-Moloney, M., and Leiter, A.B. (2004). Neurogenin 3-expressing progenitor cells in the gastrointestinal tract differentiate into both endocrine and non-endocrine cell types. Dev Biol 270, 443-454.

Scully, K.M., Skowronska-Krawczyk, D., Krawczyk, M., Merkurjev, D., Taylor, H., Livolsi, A., Tollkuhn, J., Stan, R.V., and Rosenfeld, M.G. (2016). Epithelial cell integrin beta1 is required for developmental angiogenesis in the pituitary gland. Proc Natl Acad Sci U S A 113, 13408–13413.

Seymour, P.A., Bennett, W.R., and Slack, J.M. (2004). Fission of pancreatic islets during postnatal growth of the mouse. J Anat 204, 103-116.

Sharon, N., Chawla, R., Mueller, J., Vanderhooft, J., Whitehorn, L.J., Rosenthal, B., Gurtler, M., Estanboulieh, R.R., Shvartsman, D., Gifford, D.K., et al. (2019a). A Peninsular Structure Coordinates Asynchronous Differentiation with Morphogenesis to Generate Pancreatic Islets. Cell 176, 790–804 e713.

Sharon, N., Vanderhooft, J., Straubhaar, J., Mueller, J., Chawla, R., Zhou, Q., Engquist, E.N., Trapnell, C., Gifford, D.K., and Melton, D.A. (2019b). Wnt Signaling Separates the Progenitor and Endocrine Compartments during Pancreas Development. Cell Rep 27, 2281–2291 e2285.

Shih, H.P., Gross, M.K., and Kioussi, C. (2007). Cranial muscle defects of Pitx2 mutants result from specification defects in the first branchial arch. Proc Natl Acad Sci U S A 104, 5907–5912.

Shih, H.P., Panlasigui, D., Cirulli, V., and Sander, M. (2016). ECM Signaling Regulates Collective Cellular Dynamics to Control Pancreas Branching Morphogenesis. Cell Rep 14, 169-179.

Shih, H.P., and Sander, M. (2014). Pancreas development ex vivo: culturing embryonic pancreas explants on permeable culture inserts, with fibronectin-coated glass microwells, or embedded in three-dimensional Matrigel. Methods Mol Biol 1210, 229–237.

Shih, H.P., Wang, A., and Sander, M. (2013). Pancreas organogenesis: from lineage determination to morphogenesis. Annu Rev Cell Dev Biol 29, 81–105.

Shiota, C., Prasadan, K., Guo, P., Fusco, J., Xiao, X., and Gittes, G.K. (2017). Gcg (CreERT2) knockin mice as a tool for genetic manipulation in pancreatic alpha cells. Diabetologia 60, 2399–2408.

Simon-Areces, J., Membrive, G., Garcia-Fernandez, C., Garcia-Segura, L.M., and Arevalo, M.A. (2010). Neurogenin 3 cellular and subcellular localization in the developing and adult hippocampus. J Comp Neurol 518, 1814–1824.

Speicher, T., Siegenthaler, B., Bogorad, R.L., Ruppert, R., Petzold, T., Padrissa-Altes, S., Bachofner, M., Anderson, D.G., Koteliansky, V., Fassler, R., et al. (2014). Knockdown and knockout of beta1-integrin in hepatocytes impairs liver regeneration through inhibition of growth factor signalling. Nat Commun 5, 3862.

Srinivas, S., Watanabe, T., Lin, C.S., William, C.M., Tanabe, Y., Jessell, T.M., and Costantini, F. (2001). Cre reporter strains produced by targeted insertion of EYFP and ECFP into the ROSA26 locus. BMC Dev Biol 1, 4.

Stine, R.R., Greenspan, L.J., Ramachandran, K.V., and Matunis, E.L. (2014). Coordinate regulation of stem cell competition by Slit-Robo and JAK-STAT signaling in the Drosophila testis. PLoS Genet 10, e1004713.

Subramanian, A., Tamayo, P., Mootha, V.K., Mukherjee, S., Ebert, B.L., Gillette, M.A., Paulovich, A., Pomeroy, S.L., Golub, T.R., Lander, E.S., et al. (2005). Gene set enrichment analysis: a knowledge-based approach for interpreting genome-wide expression profiles. Proc Natl Acad Sci U S A 102, 15545–15550.

Sznurkowska, M.K., Hannezo, E., Azzarelli, R., Chatzeli, L., Ikeda, T., Yoshida, S., Philpott, A., and Simons, B.D. (2020). Tracing the cellular basis of islet specification in mouse pancreas. Nat Commun 11, 5037.

T.J. Deerinck, E.B., A. Thor, M.H. Ellisman (2010). NCMIR Methods for 3D EM: A New Protocol for Preparation of Biological Specimens for Serial Block Face Scanning Electron Microscopy. In National Center for Microscopy and Imaging Research, University of California, San Diego

Taylor, B.L., Liu, F.F., and Sander, M. (2013). Nkx6.1 is essential for maintaining the functional state of pancreatic beta cells. Cell Rep 4, 1262-1275.

Thorens, B., Tarussio, D., Maestro, M.A., Rovira, M., Heikkila, E., and Ferrer, J. (2015). Ins1(Cre) knock-in mice for beta cell-specific gene recombination. Diabetologia 58, 558–565.

Tong, M., Jun, T., Nie, Y., Hao, J., and Fan, D. (2019). The Role of the Slit/Robo Signaling Pathway. J Cancer 10, 2694–2705.

van der Meulen, T., and Huising, M.O. (2014). Maturation of stem cell-derived beta-cells guided by the expression of urocortin 3. Rev Diabet Stud 11, 115–132.

van der Meulen, T., Mawla, A.M., DiGruccio, M.R., Adams, M.W., Nies, V., Dolleman, S., Liu, S., Ackermann, A.M., Caceres, E., Hunter, A.E., et al. (2017). Virgin Beta Cells Persist throughout Life at a Neogenic Niche within Pancreatic Islets. Cell Metab 25, 911–926 e916.

Vasioukhin, V., Bauer, C., Degenstein, L., Wise, B., and Fuchs, E. (2001). Hyperproliferation and defects in epithelial polarity upon conditional ablation of alpha-catenin in skin. Cell 104, 605–617.

Wakae-Takada, N., Xuan, S., Watanabe, K., Meda, P., and Leibel, R.L. (2013). Molecular basis for the regulation of islet beta cell mass in mice: the role of E-cadherin. Diabetologia 56, 856-866.

Wang, S., Matsumoto, K., Lish, S.R., Cartagena-Rivera, A.X., and Yamada, K.M. (2021). Budding epithelial morphogenesis driven by cell-matrix versus cell-cell adhesion. Cell 184, 3702-3716 e3730.

Wang, S., Sekiguchi, R., Daley, W.P., and Yamada, K.M. (2017). Patterned cell and matrix dynamics in branching morphogenesis. J Cell Biol 216, 559–570.

Weber, G.F., Bjerke, M.A., and DeSimone, D.W. (2011). Integrins and cadherins join forces to form adhesive networks. J Cell Sci 124, 1183–1193.

Win, P.W., Oakie, A., Li, J., and Wang, R. (2020). Beta-cell beta1 integrin deficiency affects in utero development of islet growth and vascularization. Cell Tissue Res 381, 163–175.

Yamamoto, H., Ehling, M., Kato, K., Kanai, K., van Lessen, M., Frye, M., Zeuschner, D., Nakayama, M., Vestweber, D., and Adams, R.H. (2015). Integrin beta1 controls VE-cadherin localization and blood vessel stability. Nat Commun 6, 6429.

Yashpal, N.K., Li, J., Wheeler, M.B., and Wang, R. (2005). Expression of beta1 integrin receptors during rat pancreas development--sites and dynamics. Endocrinology 146, 1798–1807.

Zeng, C., Mulas, F., Sui, Y., Guan, T., Miller, N., Tan, Y., Liu, F., Jin, W., Carrano, A.C., Huising, M.O., et al. (2017). Pseudotemporal Ordering of Single Cells Reveals Metabolic Control of Postnatal beta Cell Proliferation. Cell Metab 25, 1160–1175 e1111.

