## Supplementary material for "Coordination between ECM and cell-cell adhesion regulates the development of islet aggregation, architecture, and functional maturation": Key Resource Table

| REAGENT or RESOURCE | | SOURCE | | | IDENTIFIER |
| --- | --- | --- | --- | --- | --- |
| **Antibodies** | | | | | |
| Goat polyclonal anti-Pancreatic Polypeptide/PP  (for IHC 1:1000) | | Novus Biologicals | | | Cat# NB100-1793  RRID: AB_2268669 |
| Mouse monoclonal anti-Glucagon  (for IHC 1:500) | | Sigma-Aldrich | | | Cat# G2654  RRID: AB_259852 |
| Goat polyclonal anti-Glucagon  (for IHC 1:500) | | Santa Cruz Biotechnology | | | Cat# sc-7780  RRID: AB_641025 |
| Guinea polyclonal Pig anti-Insulin  (for IHC 1:2000) | | Dakocytomation | | | Cat# A0564  RRID: AB_10013624 |
| Rabbit monoclonal anti-Insulin  (for IHC & WMIHC 1:1000) | | Cell Signaling Technology | | | Cat# 3014S  RRID: AB_2126503 |
| Rabbit polyclonal anti-Chromogranin A  (for IHC & WMIHC 1:500) | | Novus Biologicals | | | Cat# NB120-15160  RRID: AB_789299 |
| Goat polyclonal anti-Chromogranin A  (for IHC & WMIHC 1:500) | | Santa Cruz Biotechnology | | | Cat# sc-1488  RRID: AB_2276319 |
| Goat polyclonal anti-Somatostatin  (for IHC 1:100) | | Santa Cruz Biotechnology | | | Cat# sc-7819  RRID: AB_2302603 |
| Purified Rat Anti-Mouse CD31  (for IHC & WMIHC 1:200) | | BD Biosciences | | | Cat# 550274  RRID: AB_393571 |
| Goat polyclonal anti-PECAM-1  (for IHC 1:500) | | R&D Systems | | | Cat# AF3628  RRID: AB_2161028 |
| Osteopontin/OPN Antibody (Unconjugated) (Spp1)  (for IHC & WMIHC 1:1000) | | R&D Systems | | | Cat# AF808  RRID: AB_2194992 |
| Guinea Pig anti-Neurogenin3  (for IHC 1:1000) | | A generous gift of Maike Sander  (Henseleit KD, Nelson SB, Kuhlbrodt K, Hennings JC, Ericson J, et al. (2005) NKX6 transcription factor activity is required for alpha- and beta-cell development in the pancreas. | | | Development (2005) 132 (13): 3139–3149. |
| Mouse monoclonal anti-Nkx6.1  (for IHC 1:500) | | DSHB | | | Cat# F55A10  RRID: AB_532378 |
| Rabbit polyclonal anti-Pdx1  (for IHC 1:500) | | Abcam | | | Cat# ab47267  RRID: AB_777179 |
| Rabbit polyclonal anti-Laminin (Lam1)  (for IHC 1:1000) | | Sigma-Aldrich | | | Cat# L9393  RRID: AB_477163 |
| Rat monoclonal anti-EPCAM  (for IHC 1:100) | | DSHB | | | Cat# G8.8  RRID: AB_2098655 |
| Goat polyclonal anti-E-cadherin  (for IHC 1:500) | | R&D Systems | | | Cat# AF748  RRID: AB_355568 |
| Rabbit polyclonal anti-MafA  (for IHC 1:200) | | Abcam | | | Cat# ab26405  RRID: AB_776146 |
| Rabbit anti-Urocortin3 (Ucn3)  (for IHC 1:1000) | | A generous gift of Mark Huising  (van der Meulen, Talitha et al. “Urocortin3 mediates somatostatin-dependent negative feedback control of insulin secretion.” 2015) | | | Nature medicine vol. 21,7 (2015): 769-76. doi:10.1038/nm.3872 |
| Goat polyclonal anti-Glut2  (for IHC 1:500) | | Santa Cruz Biotechnology | | | Cat# sc-7580  RRID: AB_641066 |
| Rat monoclonal anti-CD29 (Itgβ1)  (for IHC 1:500) | | BD Biosciences | | | Cat# 550531  RRID: AB_393729 |
| Rabbit monoclonal anti-Ki67  (for IHC 1:500) | | Cell Signaling | | | Cat# 12202S (D3B5)  RRID: AB_2620142 |
| Rabbit anti-Sox9 antibody  (for IHC 1:500) | | EMD Millipore | | | Cat# AB5535  RRID: AB_2239761 |
| Rabbit anti-ERO1b antiserum  (for IHC 1:300) | | A generous gift of David Ron (Zito, Ester et al. “ERO1-beta, a pancreas-specific disulfide oxidase, promotes insulin biogenesis and glucose homeostasis.”) | | | J Cell Biol. 2010 Mar 22;188(6):821-32.  doi:10.1083/jcb.200911086. |
| Cy™3 AffiniPure Donkey Anti-Goat IgG (H+L)  (for IHC 1:2000) | | Jackson ImmunoResearch | | | Cat# 705-165-147  RRID: AB_2307351 |
| Alexa Fluor® 488 AffiniPure Donkey Anti-Goat IgG (H+L)  (for IHC 1:1000) | | Jackson ImmunoResearch | | | Cat# 705-545-147  RRID: AB_2336933 |
| Alexa Fluor® 647 AffiniPure Donkey Anti-Goat IgG (H+L)  (for IHC 1:200 – 1:500) | | Jackson ImmunoResearch | | | Cat# 705-605-147  RRID: AB_2340437 |
| Cy™3 AffiniPure Donkey Anti-Guinea Pig IgG (H+L)  (for IHC 1:2000) | | Jackson ImmunoResearch | | | Cat# 706-165-148  **RRID:** AB_2340460 |
| Alexa Fluor® 488 AffiniPure Donkey Anti-Guinea Pig IgG (H+L)  (for IHC 1:1000) | | Jackson ImmunoResearch | | | Cat# 706-545-148 **RRID:** AB_2340472 |
| Alexa Fluor® 647 AffiniPure Donkey Anti-Guinea Pig IgG (H+L)  (for IHC 1:200 – 1:500) | | Jackson ImmunoResearch | | | Cat# 706-605-148  **RRID: AB_2340476** |
| Cy™3 AffiniPure Donkey Anti-Rabbit IgG (H+L)  (for IHC 1:2000) | | Jackson ImmunoResearch | | | Cat# 711-165-152 **RRID:** AB_2307443 |
| Alexa Fluor® 488 AffiniPure Donkey Anti-Rabbit IgG (H+L)  (for IHC 1:1000) | | Jackson ImmunoResearch | | | Cat# 711-545-152  **RRID:** AB_2313584 |
| Alexa Fluor® 647 AffiniPure Donkey Anti-Rabbit IgG (H+L)  (for IHC 1:200 – 1:500) | | Jackson ImmunoResearch | | | Cat# 711-605-152 **RRID:** AB_2492288 |
| Cy™3 AffiniPure Donkey Anti-Rat IgG (H+L)  (for IHC 1:2000) | | Jackson ImmunoResearch | | | Cat# 712-165-153  **RRID:** AB_2340667 |
| Alexa Fluor® 488 AffiniPure Donkey Anti-Rat IgG (H+L)  (for IHC 1:1000) | | Jackson ImmunoResearch | | | Cat# 712-545-153  **RRID:** AB_2340684 |
| Alexa Fluor® 647 AffiniPure Donkey Anti-Rat IgG (H+L)  (for IHC 1:200 – 1:500) | | Jackson ImmunoResearch | | | Cat# 712-605-153 **RRID:** AB_2340694 |
| Dolichos Biflorus Agglutinin (DBA), Biotinylated  (for IHC 1:500) | | Vector Laboratories | | | Cat# B1035  RRID: AB_2314288 |
| Dolichos Biflorus Agglutinin (DBA), Rhodamine  (for IHC 1:500) | | Vector Laboratories | | | Cat# RL-1032-2  RRID: AB_2336396 |
| DAPI (4',6-Diamidino-2-Phenylindole, Dihydrochloride)  (for IHC 1:2000) | | ThermoFisher Scientific | | | Cat# D1306  RRID: AB_2629482 |
| Phalloidin-iFluor 647 Reagent  (for IHC 1:1000) | | Abcam | | | Cat# ab176759 |
| Alexa Fluor® 647 Streptavidin  (for IHC 1:500) | | Jackson ImmunoResearch | | | Cat# 016-600-084  **RRID:** AB_2341101 |
| Cy™3 Streptavidin  (for IHC 1:500) | | Jackson ImmunoResearch | | | Cat# 016-160-084 RRID: AB_2337244 |
| **Chemicals, peptides, and recombinant proteins** | | | | | |
| Tween™ 20, Fisher BioReagents™ | | Fisher Scientific | | | BP337-500 |
| Triton™ X-100 | | Sigma | | | X100-1L |
| VectaShield Mounting Medium for Fluorescence | | Vector Laboratories | | | H-1000 |
| Paraformaldehyde | | Sigma-Aldrich | | | P6148-500G |
| Methanol | | VWR Chemical | | | BDH1135-4LP |
| DMSO | | ATCC | | | 4-X-5 |
| Hydrogen Peroxide, ACS, 30%, Stabilized | | VWR Chemical | | | BDH7690-1 |
| Benzyl Benzoate (ACROS organics) | | Fisher Scientific | | | AC105862500 |
| Benzyl Alcohol | | Fisher Scientific | | | 100-51-6 |
| D-Glucose | | Mallinckrodt | | | 4192 |
| Bovine Albumin Fraction V (7.5% Solution) | | Thermo Fisher | | | 15260037 |
| TrypLE Express, with phenol red, Gibco | | Fisher Scientific | | | 12605-010 |
| EDTA, 0.5M, pH 8.0 | | Corning | | | 46-034-CI |
| Fetal Bovine Serum | | Biofluid Technologies | | | BT-101-500-D |
| Heparin sodium salt from porcine intestinal mucosa | | Sigma-Aldrich | | | H3149-100KU |
| CMRL 1066, Supplemented CIT Modification | | Mediatech | | | 98-304-CV |
| RNase-Free DNase Set | | Qiagen | | | 79254 |
| Penicillin-Streptomycin | | Gibco | | | 15070-063 |
| Fibronectin Human Protein, Plasma | | Gibco | | | 33016-015 |
| DMEM/F12, HEPES, no phenol red | | Gibco | | | 11-039-21 |
| Roche Blocking | | Roche | | | 17091700 |
| Donor Donkey Serum | | Gemini Bio | | | 100-151-500 |
| Cacodylic Acid, Sodium Salt, trihydrate | | TedPella Inc | | | 18851 |
| Glutaraldehyde, 25% EM grade | | TedPella Inc | | | 18426 |
| Thiocarbohydrazide ≥98.0% | | VWR International | | | TCT1136-25G |
| Buffer RLT Plus Lysis Buffer | | Qiagen | | | 1053393 |
| 2-Mercaptoethanol | | Sigma-Aldrich | | | M3148-25ML |
| Ethanol, Absolute 200 proof | | Fisher BioReagents | | | BP2818100 |
| **Critical commercial assays** | | | | | |
| AlphaTrak2 Blood Glucose Test Strips | | Zoetis | | | 71681-01 |
| AlphaTrak2 Meter | | Zoetis | | | 71676-01 |
| Streptavidin/Biotin Blocking Kit | | Vector Laboratories | | | SP-2002 |
| Rneasy Micro Kit | | Qiagen | | | 74304 |
| Rnase Zap | | Invitrogen | | | AM9780 |
| KAPA RNA HyperPrep Kit with RiboErase (HMR) | | Roche | | | KK8560 |
| AxyPrep Mag PCR Clean-up kit | | Axigen | | | MAG-PCR-CL-1 |
| NovaSeq 6000 Reagent Kit | | Illumina® | | |  |
| DeadEnd™ Fluorometric TUNEL System | | Promega | | | G3250 |
| M.O.M.® (Mouse on Mouse) Immunodetection Kit, Basic | | Vector Laboratories | | | BMK-2202 |
| PowerUp^TM^ SYBR^TM^ Green Master mix | | Applied Biosystems | | | A25742 |
| SuperScript^TM^ First-Strand Synthesis SuperMix for qRT-PCR | | Invitrogen | | | 11752-050 |
| Ultrasensitive Mouse Insulin ELISA | | Mercodia | | | 10-1249-01 |
| **Deposited data** | | | | | |
| RNA sequencing data of Itgb1 KO murine pancreatic islets. | | https://www.ncbi.nlm.nih.gov/geo/ | | | GSE153187 |
| RNA sequencing data of Ctnna1 KO murine pancreatic islets. | | https://www.ncbi.nlm.nih.gov/geo/ | | | GSE190788 |
| **Experimental models: Organisms/strains** | | | | | |
| Mouse: B6.FVB(Cg)-*Tg(Neurog3-cre)C1Able/J* | | Jackson Laboratory, Bar Harbor, ME | | | RRID:IMSR_JAX:006333 |
| Mouse: B6;129-*Itgb1^tm1Efu^/J* | | Jackson Laboratory, Bar Harbor, ME | | | RRID:IMSR_JAX:004605 |
| Mouse: B6;129-*Ctnna1^tm1Efu^/J* | | Jackson Laboratory, Bar Harbor, ME | | | RRID:IMSR_JAX:004604 |
| Mouse: B6;129S-*Gcg^tm1.1(icre)Gkg^/J* | | Jackson Laboratory, Bar Harbor, ME | | | RRID:IMSR_JAX:030663 |
| Mouse: B6(Cg)-*Ins1^tm1.1(cre)Thor^/J* | | Jackson Laboratory, Bar Harbor, ME | | | RRID:IMSR_JAX:026801 |
| Mouse: B6.129(Cg)-*Gt(ROSA)26Sor^tm4(ACTB-tdTomato,-EGFP)Luo^*/J | | Jackson Laboratory, Bar Harbor, ME | | | RRID:IMSR_JAX:007676 |
| Mouse: B6.129X1-*Gt(ROSA)26Sor^tm1(EYFP)Cos^/J* | | Jackson Laboratory, Bar Harbor, ME | | | RRID:IMSR_JAX:006148 |
| B6.FVB(Cg)-*Tg(Ucn3-cre)KF43Gsat/Mmucd* | | Mutant Mouse Resource & Research Centers, Novi, MI | | | RRID:MMRRC_037417-UCD |
| **PCR Primers** | | | | | |
| **Target** | **Primer Name** | | **Sequence** | **Mouse Strain** | |
| Generic Cre | oIMR1084 | | GCG GTC TGG CAG TAA AAA CTA TC | B6.FVB(Cg)-*Tg(Neurog3-cre)C1Able/J*  And  B6(Cg)-*Ins1^tm1.1(cre)Thor^/J*  And  B6.FVB(Cg)-*Tg(Ucn3-cre)KF43Gsat/Mmucd* | |
| Generic Cre | oIMR1085 | | GTG AAA CAG CAT TGC TGT CAC TT | B6.FVB(Cg)-*Tg(Neurog3-cre)C1Able/J*  And  B6(Cg)-*Ins1^tm1.1(cre)Thor^/J*  And  B6.FVB(Cg)-*Tg(Ucn3-cre)KF43Gsat/Mmucd* | |
| Generic Cre | oIMR7338 | | CTA GGC CAC AGA ATT GAA AGA TCT | B6.FVB(Cg)-*Tg(Neurog3-cre)C1Able/J*  And  B6(Cg)-*Ins1^tm1.1(cre)Thor^/J*  And  B6.FVB(Cg)-*Tg(Ucn3-cre)KF43Gsat/Mmucd* | |
| Generic Cre | oIMR7339 | | GTA GGT GGA AAT TCT AGC ATC ATC C | B6.FVB(Cg)-*Tg(Neurog3-cre)C1Able/J*  And  B6(Cg)-*Ins1^tm1.1(cre)Thor^/J*  And  B6.FVB(Cg)-*Tg(Ucn3-cre)KF43Gsat/Mmucd* | |
| β1 integrin floxed | oIMR1906 | | CGG CTC AAA GCA GAG TGT CAG TC | B6;129-*Itgb1^tm1Efu^/J* | |
| β1 integrin floxed | oIMR1907 | | CCA CAA CTT TCC CAG TTA GCT CTC | B6;129-*Itgb1^tm1Efu^/J* | |
| α-catenin floxed | oIMR1902 | | CAT TTC TGT CAC CCC CAA AGA CAC | B6;129-*Ctnna1^tm1Efu^/J* | |
| α-catenin floxed | oIMR1903 | | GCA AAA TGA TCC AGC GTC CTG GG | B6;129-*Ctnna1^tm1Efu^/J* | |
| Gcg^iCre^ | oIMR7338 | | CTA GGC CAC AGA ATT GAA AGA TCT | B6;129S-*Gcg^tm1.1(icre)Gkg^/J* | |
| Gcg^iCre^ | oIMR7339 | | GTA GGT GGA AAT TCT AGC ATC ATC C | B6;129S-*Gcg^tm1.1(icre)Gkg^/J* | |
| Gcg^iCre^ | oIMR9266 | | AGA TGC CAG GAC ATC AGG AAC CTG | B6;129S-*Gcg^tm1.1(icre)Gkg^/J* | |
| Gcg^iCre^ | oIMR9267 | | ATC AGC CAC ACC AGA CAC AGA GAT C | B6;129S-*Gcg^tm1.1(icre)Gkg^/J* | |
| Rosa^mT/mG^ | 9655 | | CCA GGC GGG CCA TTT ACC GTA AG | *Gt(ROSA)26Sor^tm4(ACTB-tdTomato,-EGFP)Luo^*/J | |
| Rosa^mT/mG^ | oIMR8545 | | AAA GTC GCT CTG AGT TGT TAT | *Gt(ROSA)26Sor^tm4(ACTB-tdTomato,-EGFP)Luo^*/J | |
| Rosa^mT/mG^ | oIMR8546 | | GGA GCG GGA GAA ATG GAT ATG | *Gt(ROSA)26Sor^tm4(ACTB-tdTomato,-EGFP)Luo^*/J | |
| R26R-EYFP | 21306 | | CTG GCT TCT GAG GAC CG | B6.129X1-*Gt(ROSA)26Sor^tm1(EYFP)Cos^/J* | |
| R26R-EYFP | 24500 | | CAG GAC AAC GCC CAC ACA | B6.129X1-*Gt(ROSA)26Sor^tm1(EYFP)Cos^/J* | |
| R26R-EYFP | 24951 | | AGG GCG AGG AGC TGT TCA | B6.129X1-*Gt(ROSA)26Sor^tm1(EYFP)Cos^/J* | |
| R26R-EYFP | 24952 | | TGA AGT CGA TGC CCT TCA G | B6.129X1-*Gt(ROSA)26Sor^tm1(EYFP)Cos^/J* | |
| **qPCR Primers** | | | | | |
| **Gene** | **Primer** | **Sequence** | | | **Primer Bank ID** |
| Itgb1 | mItgb1-F | ATGCCAAATCTTGCGGAGAAT | | | 52722a1 |
|  | mItgb1-R | TTTGCTGCGATTGGTGACATT | | |  |
| Slc2a2 | mSlc2a2-F | TCAGAAGACAAGATCACCGGA | | | 13654262a1 |
|  | mSlc2a2-R | GCTGGTGTGACTGTAAGTGGG | | |  |
| Unc3 | mUcn3-F | AAGCCTCTCCCACAAGTTCTA | | | 21492632a1 |
|  | mUcn3-R | GAGGTGCGTTTGGTTGTCATC | | |  |
| MafA | mMafA-F | AGGAGGAGGTCATCCGACTG | | | 23503735a1 |
|  | mMafA-R | CTTCTCGCTCTCCAGAATGTG | | |  |
| MafB | mMafB-F | TTCGACCTTCTCAAGTTCGACG | | | 23308601a1 |
|  | mMafB-R | TCGAGATGGGTCTTCGGTTCA | | |  |
| Arx | mArx-F | GGCCGGAGTGCAAGAGTAAAT | | | 26024213a1 |
|  | mArx-R | TGCATGGCTTTTTCCTGGTCA | | |  |
| Etv1 | mEtv1-F | TTAAGTGCAGGCGTCTTCTTC | | | 26328055a1 |
|  | mEtv1-R | GGAGGCCATGAAAAGCCAAA | | |  |
| Aldh1a3 | mAldh1a3-F | GGGTCACACTGGAGCTAGGA | | | 31542123a1 |
|  | mAldh1a3-R | CTGGCCTCTTCTTGGCGAA | | |  |
| Hk1 | mHk1-F | CGGAATGGGGAGCCTTTGG | | | 309289a1 |
|  | mHk1-R | GCCTTCCTTATCCGTTTCAATGG | | |  |
| LdhA | mLdhA-F | TGTCTCCAGCAAAGACTACTGT | | | 6754524a1 |
|  | mLdhA-R | GACTGTACTTGACAATGTTGGGA | | |  |
| Ctnna1 | mCtnna1-F | AAGTCTGGAGATTAGGACTCTGG | | | 6753294a1 |
|  | mCtnna1-R | ACGGCCTCTCTTTTTATTAGACG | | |  |
| Slc27a2 | mSlc27a2-F | TCCTCCAAGATGTGCGGTACT | | | 6755548a1 |
|  | mSlc27a2-R | TAGGTGAGCGTCTCGTCTCG | | |  |
| Peg10 | mPeg10-F | TGCTTGCACAGAGCTACAGTC | | | 31376257a1 |
|  | mPeg10-R | AGTTTGGGATAGGGGCTGCT | | |  |
| Sst | mSst-F | ACCGGGAAACAGGAACTGG | | | 6678035a1 |
|  | mSst-R | TTGCTGGGTTCGAGTTGGC | | |  |
| Aldh1a1 | mAldh1a1-F | ATACTTGTCGGATTTAGGAGGCT | | | 7304881a1 |
|  | mAldh1a1-R | GGGCCTATCTTCCAAATGAACA | | |  |
| Cxcl14 | mCxcl14-F | GAAGATGGTTATCGTCACCACC | | | 9625004a1 |
|  | mCxcl14-R | CGTTCCAGGCATTGTACCACT | | |  |
| PPIA (Cycloipin) | mPPIA-Fw | GAGCTGTTTGCAGACAAAGTTC | | | 6679438c1 |
|  | mPPIA-Rv | CCCTGGCACATGAATCCTGG | | |  |
| **Software and algorithms** | | | | | |
| ImageJ Software | | https://imagej.nih.gov/ij/index.html | | | RRID:SCR_003070 |
| Imaris Microscopy Image Analysis Software | | https://imaris.oxinst.com/ | | | RRID:SCR_007370 |
| QuPath Quantitative Pathology & Bioimage Analysis | | https://qupath.github.io/ | | | RRID:SCR_018257 |
| ZEISS ZEN Microscope Software | | http://www.zeiss.com/microscopy/en_us/products/microscope-software/zen.html#introduction | | | RRID:SCR_013672 |
| Star software (v.020201) | | http://code.google.com/p/rna-star/ | | | RRID:SCR_004463 |
| DAVID software (v.6.8) | | http://david.abcc.ncifcrf.gov/ | | | RRID:SCR_001881 |
| Featurecount software | | http://bioinf.wehi.edu.au/featureCounts/ | | | RRID:SCR_012919 |
| R-Project for Statistical Computing | | http://www.r-project.org/ | | | RRID:SCR_001905 |
| GraphPad Prism Statistical Analysis Software | | http://www.graphpad.com/ | | | RRID:SCR_002798 |
| Applied Biosystems QuantStudio^TM^ 6 Real Time PCR System - QuantStudio^TM^ Real-Time PCR Software v1.3 | | https://www.thermofisher.com/us/en/home/global/forms/life-science/quantstudio-6-7-flex-software.html | | | RRID: SCR_020239 |
