## Supplementary movie S1-S3 for "Coordination between ECM and cell-cell adhesion regulates the development of islet aggregation, architecture, and functional maturation"

### Slide 1
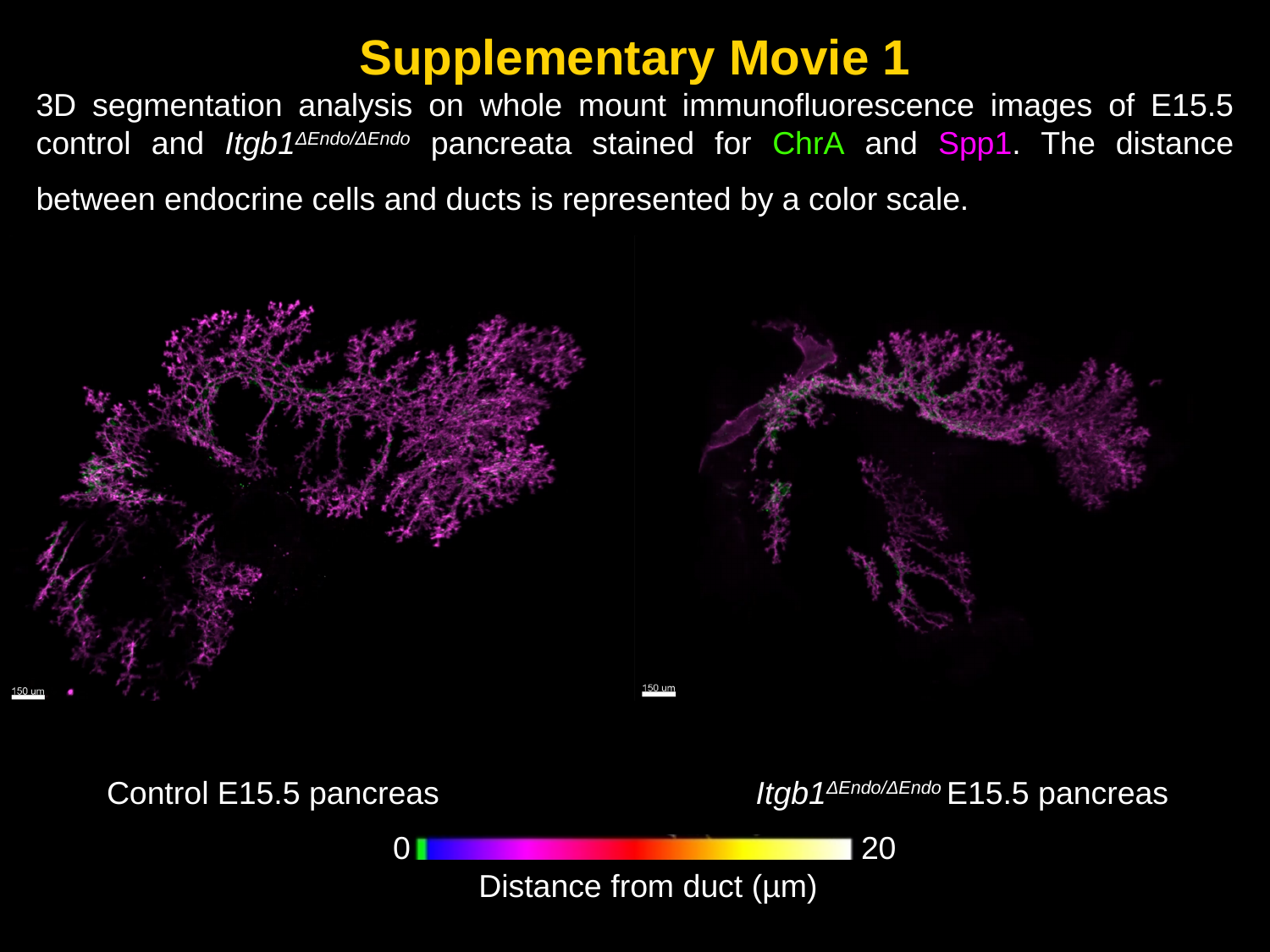

Supplementary Movie 1
3D segmentation analysis on whole mount immunofluorescence images of E15.5 control and Itgb1^ΔEndo/ΔEndo^ pancreata stained for ChrA and Spp1. The distance between endocrine cells and ducts is represented by a color scale.
Control E15.5 pancreas
Itgb1^ΔEndo/ΔEndo^ E15.5 pancreas
0
20
Distance from duct (µm)

### Slide 2
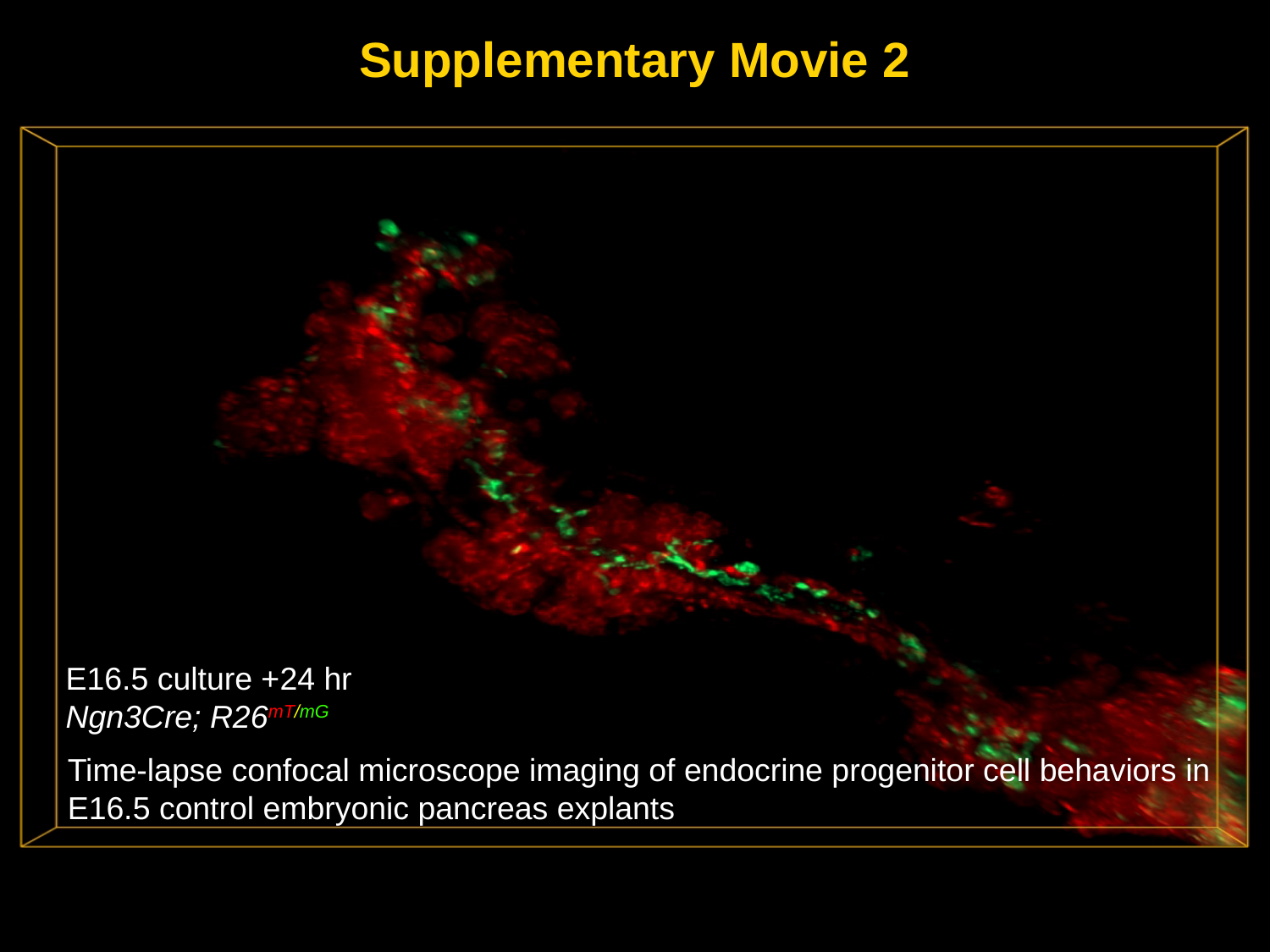

Supplementary Movie 2
E16.5 culture +24 hr
Ngn3Cre; R26^mT/mG^
Time-lapse confocal microscope imaging of endocrine progenitor cell behaviors in E16.5 control embryonic pancreas explants

### Slide 3
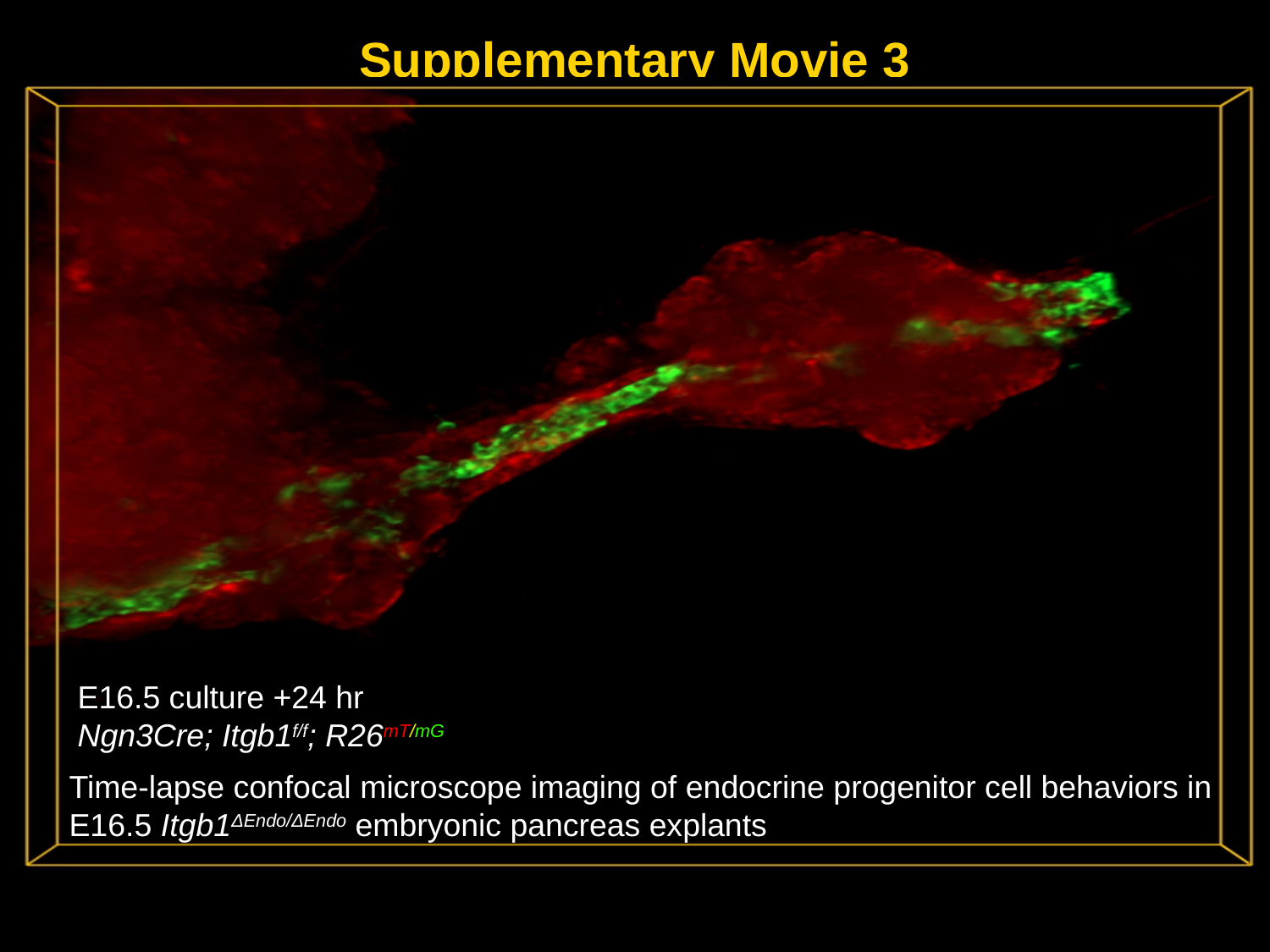

Supplementary Movie 3
E16.5 culture +24 hr
Ngn3Cre; Itgb1^f/f^; R26^mT/mG^
Time-lapse confocal microscope imaging of endocrine progenitor cell behaviors in E16.5 Itgb1^ΔEndo/ΔEndo^ embryonic pancreas explants
